# Evidence for admixture and rapid evolution during glacial climate change in an alpine specialist

**DOI:** 10.1101/2023.09.21.558886

**Authors:** Yi-Ming Weng, David H. Kavanaugh, Peter L. Ralph, Gilia Patterson, Sean D. Schoville

## Abstract

The pace of current climate change is expected to be problematic for alpine flora and fauna, as their adaptive capacity may be limited by small population size. Yet despite substantial genetic drift following post-glacial recolonization of alpine habitats, alpine species are notable for their success in surviving highly heterogeneous environments. Population genomic analyses demonstrating how alpine species have adapted to novel environments with limited genetic diversity remain rare, yet are important in understanding the potential for species to respond to contemporary climate change. In this study, we explored the evolutionary history of alpine ground beetles in the *Nebria ingens* complex, including the demographic and adaptive changes that followed the last glacier retreat. Using whole genome data from hundreds of beetles, to test alternative models of evolutionary divergence in the species complex, we found evidence that the *Nebria ingens* complex has been formed by past admixture of lineages responding to glacial cycles. Recolonization of alpine sites involved a distributional range shift to higher elevation, which was accompanied by a reduction in suitable habitat and the emergence of complex spatial genetic structure. We also used genome-wide association and genotype-environment association methods to look for genetic pathways involved in adaptation to heterogeneous new environments during this range shift. The identified genes were enriched for functions broadly associated with abiotic stress responses, with strong evidence for adaptation to hypoxia-related pathways. The results demonstrate that despite rapid environmental changes, alpine beetles in the *N. ingens* complex have shown rapid physiological evolution.

## Introduction

As the velocity of climate change accelerates (Loarie et al. 2009), a critical question is what role evolution will play in species’ survival under rapidly changing environmental conditions? A general view is that adaptation is unlikely for many species due to the rapid pace of change (Jezkova and Wiens 2016). The stakes appear high for many species because effective population size and mutation rates are generally very low (Ellegren and Galtier 2016; Buffalo 2021), requiring high standing genetic diversity to adapt to rapidly changing environmental conditions (Orr and Unckless 2008). It seems particularly challenging for selection to operate when species undergo extensive spatial displacement during environmental change, or demographic fluctuations, as standing genetic diversity erodes under these conditions (Bell and Collins 2008; Excoffier et al. 2009). Despite these general constraints for evolution under rapid environmental change, the past holds many success stories that can be used to understand the interaction of demography and selection, as well as the mechanisms of adaptation, under rapid environmental change (Hof et al. 2011).

Alpine species are useful for expanding our knowledge of rapid environmental change, as these species experienced extensive distributional and demographic shifts in response to alpine deglaciation (Schoville and Roderick 2009). Following the Last Glacial Maximum (∼26.5 kya), glaciers receded rapidly (starting at ∼17 kya) over the course of 500-1,000 years (Phillips et al. 2009). Many cold-adapted alpine species colonized their current altitudinal range from lowland glacial refugia following retreating glaciers (Porinchu et al. 2003; Segarra-Moragues et al. 2007; Pan et al. 2020; Weng et al. 2021a). This spatial colonization process led to strong genetic drift (Knowles & Massatti, 2017), yet several recent population genomics studies document rapid adaptive evolution in alpine species (Rellstab et al. 2020; Bohutínská et al. 2021; Cheng et al. 2021; McCulloch et al. 2021). Notably, many of these studies demonstrate convergent adaptation in the same molecular pathways, suggesting selection from standing variation to be a common process. However, the generality of adaptive responses to past episodes of rapid climate change is not clear.

Survival in alpine environments involves several significant environmental challenges that allow for predictions of natural selection. First, species experience lower oxygen levels (hypoxia) at higher elevation (Dillon et al. 2006), which has led to adaptive divergence across elevational gradients in a number of animal taxa (Cheviron et al. 2012; Zhao et al. 2013; Foll et al. 2014; Ding et al. 2018). Adaptive phenotypes include enhancements to the oxygenation efficiency of the circulatory and pulmonary systems, as well as cellular adjustments to maintain oxygen homeostasis and reduce cellular stress (Lee et al. 2020). Gross morphological changes can occur. For example, in insects, morphological adaptations to hypoxia include reductions in body size to increase the diffusion of the oxygen, enlarged and multifurcating trachea, and alteration or reduction of wing morphology and musculature (Loudon 1989; Hoback and Stanley 2001; Henry and Harrison 2004; Dillon et al. 2006). Additionally, physiological adaptations involve the regulation of anaerobic metabolism and aerobic respiration involving insulin catabolism, as well as management of reactive oxygen species formation (Zhao et al. 2013; Ding et al. 2018; Harrison et al. 2018). A second important environmental challenge is the decrease in moisture as elevation increases, due to lower water vapor pressure. Drier air coupled with a windier alpine environment causes strong selective pressure for managing desiccation stress (Sømme 1989; Sinclair 2000a; Dillon et al. 2006). Finally, a third important challenge involves thermal conditions in alpine environments, which are cold, diurnally variable, and strongly seasonal. Together, these different environmental stressors can drive synergistic or conflicting selection pressures. For example, the suppression of metabolic rate can be selected by cold, desiccation, and hypoxic environmental factors, whereas selection to increase tracheal capacity to limit hypoxia directly conflicts with the need to close the tracheal system to avoid water loss.

To refine our understanding of how both demography and selection shape genomic variation during rapid climate change, we integrate species paleodistribution models, demographic analyses, and genome scans with whole genome sequencing data to explore the impact of spatial population expansion for a flightless alpine beetle. Specifically, we investigate the evolutionary history of alpine ground beetles in the *Nebria ingens* (Carabidae: Nebriinae) complex, use paleoclimatic data to predict the dynamic of species distributions through time, and employ genome scans (GS) and gene-environment association (GEA) tests to identify signatures of selection in response to high elevation conditions and local environmental variation across the alpine landscape. Our focal taxon, the *N. ingens* species complex colonized high elevation (>3000 meters) from low elevation glacial refugia to occupy patchily distributed seep habitats and snowfields in the nival zone of the Sierra Nevada Mountains of California (**supplementary file 1: Figure S1**) (Weng et al. 2021a). The *N. ingens* species complex is comprised of three distinct genetic lineages, two of which (*N. ingens* and *N. riversi*) are considered distinct species (Weng et al. 2021a). The third lineage is genetically distinct but of unclear taxonomic status due to its intermediate geographical position and morphological features. In *N. ingens*, we expect that metabolic pathways directly tied to low oxygen metabolism are targets of selection, rather than metabolic performance in cold environments, as cold specialization is broadly conserved in the congeners of *N. ingens* (Slatyer and Schoville 2016; Schat et al. 2022). We also expect that variation in snowpack drives divergent selection across the landscape, as snowpack dramatically reshapes the microhabitat (*e.g*. local temperature and water availability) of these alpine beetles.

## Results

### Genomic Diversity and Population Structure

We sequenced and genotyped 382 beetle samples from across the range, which included 100 *N. riversi*, 132 *N. ingens*, and 150 intermediates (**supplementary file 1: Table S1**). We assessed four SNP filtering methods to minimize the genotype mismatch rate (the rate of genotype mismatch between callsets of low-coverage sequence and ultra-deep coverage from an individual beetle, sample SDS06-306A). The best variant quality score recalibration (VQSR) method had a 2.94% mismatch rate and resulted in 5.5 million SNPs with 17.53% missing data (**supplementary file 1: Table S2 and Figure S2–S4**). We then imputed the missing data, which only slightly increased the mismatch rate to 3.5%. After removing loci with a minor allele frequency less than 5% and eight samples with missing rate > 90%, 1,238,058 SNPs remain for downstream analyses (referred to as the “maf dataset”). We also generated a dataset with LD thinning to remove the effect of correlated genotypes as follows. Genome-wide, mean LD decays within roughly 5 Kb. Examining LD within this range, the correlated genotypes were thinned, resulting in a final set of 378,532 SNPs (**supplementary file 1: Figure S5–S6**). This dataset (referred to as “maf + LD filtered”) was used to estimate genetic population structure including principal component analysis (PCA) and sparse non-negative matrix factorization (sNMF).

We found that PCA generates a U-shaped pattern of population structure using the first two components, forming a continuous spread of points in PC space, with *N.ingens* arrayed along PC1 (14.8%); *N. riversi* arrayed along PC2 (11.2%), and intermediates at their nexus. Furthermore, samples from the same sites are clustered together (**supplementary file 1: Figure S7**). Results from clustering algorithms were inconclusive, suggesting continuous patterns of isolation-by-distance and a clinal change in ancestry: for instance, using the clustering algorithm sNMF, the optimal number of clusters was difficult to determine (cross-entropy gradually declines to K=10 in **supplementary file 1: Figure S8**). At K=3, clusters match each lineage, but greater structure (K=5 to K=7) is clearly associated with geography. This mirrors earlier results using genotype-by-sequencing data (Weng et al, 2021a, 311 individuals and 21,238 SNPs), where population ancestry was linked to drainage basins. Examining patterns of pairwise genetic divergence, we found that relative differentiation measured by *F*_ST_ is sensitive to levels of standing genetic diversity in the locations being compared (especially comparisons involving northern *N. riversi*), whereas absolute differentiation, *d_XY,_* is more robust as a tool of pairwise comparison (**supplementary file 1: Figure S9**), as has been observed elsewhere (Cruickshank and Hahn 2014).

### Lineage Formation of the Morphologically Intermediate Populations

To infer the origin of the intermediate beetles, we employed approximate Bayesian computation (ABC) analysis to assess the fit of two competing models: a model of admixture where two main lineages hybridize to form a third lineage, and a model where all three lineages independently diverge (**Figure 1**). The admixture model provides the best fit based on both Bayes factors and the proportion of accepted simulations using a rejection algorithm (both are near 1 for the admixture model). The *d_XY_* summary statistics have a good fit to the observed data under both models, but the observed outgroup-*f3* summary statistic only fits the distribution of the admixture model (**supplementary file 1: Figure S10**). Based on the posterior distribution of the admixture model, we estimated the divergence time (τ_1_) to be around 1.06 mya (95% credible interval 0.79-1.32 mya) and admixture time (τ_2_) to be around 0.8 mya (0.69-0.91 mya). The genetic contribution of each parental population (*m*_1-2_ and *m*_3-2_) was estimated as 0.48 (0.29 – 0.60) and 0.4 (0.15-0.70), respectively, and the mean effective population size was estimated for the ancestral population as *N_A_*=72,400 (2,816-83,123), for *N. riversi* as *N_c_*= 5,668 (708-11,944), for *N. ingens* as *N_a_*=12,575 (12,561-12,586), and for the intermediate lineage as *N_s_*= 5,478 (908-10,484).

**Figure 1.**
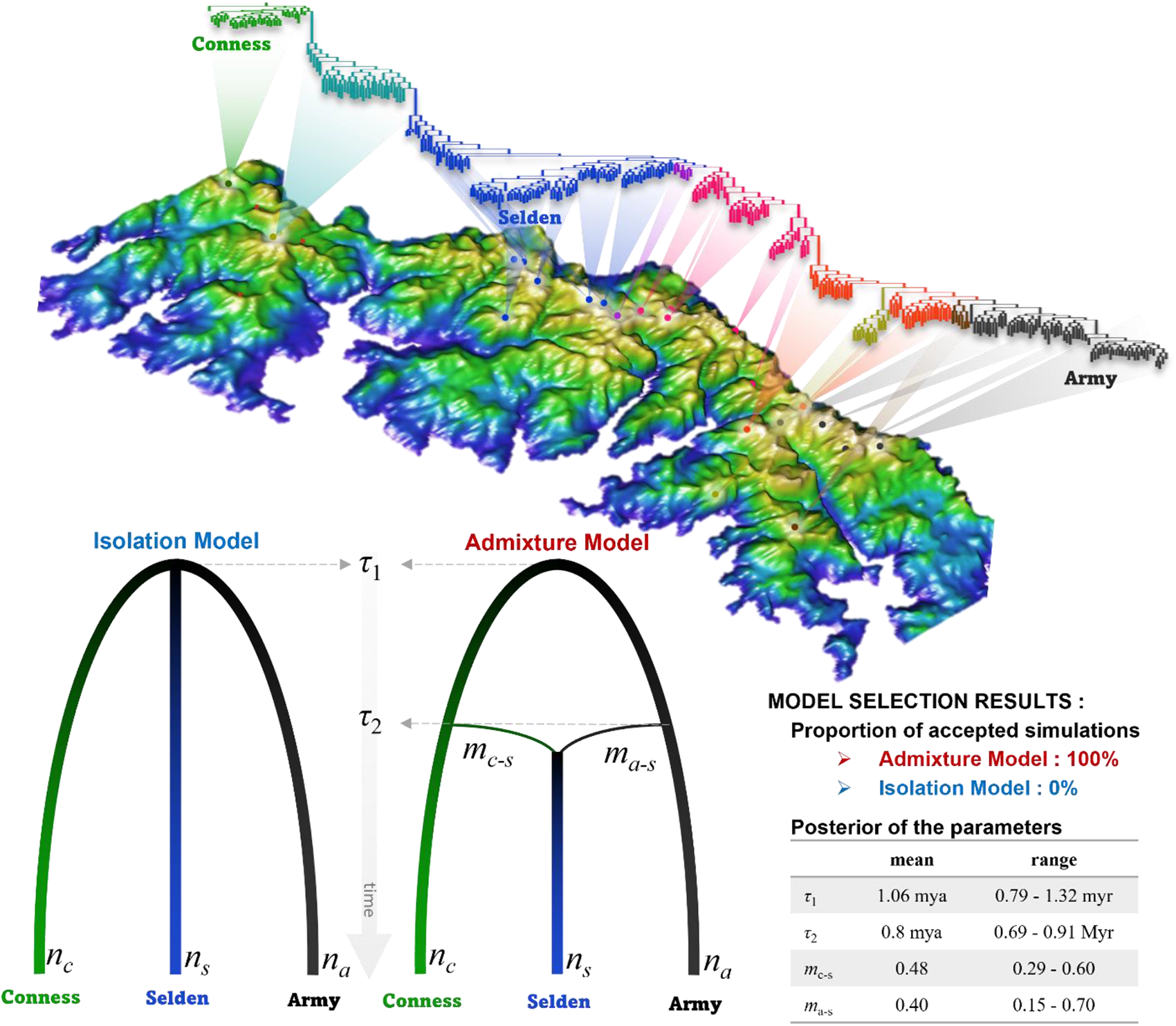
The two competing demographic models (isolation and admixture models). *τ*_1_ denotes the divergence time of the three lineages, *τ*_2_ represents the time of admixture, and *m*_c-s_ is the gene contribution of intermediate lineage (Selden) from *N. riversi* (Conness). The model comparison using Approximate Bayesian Computation shows the intermediate lineage (Selden) in the *Nebria ingens* complex formed as a consequence of nearly equal admixture between ancestors of modern samples of *N. ingens* (at Army) and *N. riversi* (at Conness). The population tree on the top with the 3D map of the Sierra Nevada in California was adapted from Weng et al. 2021a; the 3D map was drawn using the R package *rasterVis* v 0.51.5 (Lamigueiro et al. 2023) with the digital elevation model from the WorldClim database (Fick and Hijmans 2017).

### Recolonization following the last glacial maximum

To reconstruct recent demographic change following the last glacial maximum (LGM), we first generated a series of nine paleoclimate species distribution models spanning the LGM to present day climate, at intervals of ca. 2,000**–**4,000 years (**supplementary file 1: Figure S11**). In the LGM distribution model, the total area of suitable habitat was approximately 10.6-fold greater than at present, and the mean elevation (2,000 meters) was 1,500 meters lower than at present (**supplementary file 1: Figure S12**). A continuous shift from the peripheral low-elevation areas toward the peaks of the mountains can be seen across the time series of distribution models, suggesting a reduction of suitable habitat since the LGM and a continuous shift towards higher elevation (**supplementary file 1: Figure S11–S12**).

### Genomic Scan of Selection

To detect adaptive variation associated with alpine environments (selection following post-glacial recolonization), we employed genome scans to identify genomic signatures of natural selection that were shared across populations using the “maf” dataset (beetles in the *N.* ingens complex live in discrete patches across the landscape, so we refer to distinct sampling locations as “populations”). Using OmegaPlus with a 0.995 quantile threshold for significance (**supplementary file 1: Figure S13**), we identified 7,236 sites corresponding to 533 genes (**supplementary file 2: OmegaPlus top genes**). The same threshold was applied to each of 21 populations using the composite selection test of RAiSD and resulted in 1,415 to 3,201 outlier SNPs per population, or 170 to 479 genes, respectively (**supplementary file 3**). The union of the RAiSD candidate genes from all populations contains 2,741 genes, where 135 are detected in at least one-third of the tested populations (≥ 7 populations) (**supplementary file 2: RAiSD union genes**). Based on the intersection of both outlier approaches, we found 162 genes (the top 20 genes are listed in **Table 1)**, where several of the genes encode products associated with oxygen limitation, including lipid/hydrocarbon metabolic process, insulin regulation, and hypoxia-inducible factor α-subunits (HIFα) (*Foxk1* gene, He et al. 2018) (**supplementary file 2: OmegaPlus RAiSD intersect genes**). A gene ontology (GO) analysis of this set of 162 genes shows that 231 GO terms were significantly enriched. Among 214 biological process terms, notable associations for alpine environmental adaptation include muscle and neuron cell development, insulin-mediated metabolism, response to stress, regulation of homeostasis, and sperm viability (**supplementary file 2: WGS enriched BP terms**).

**Table 1.**
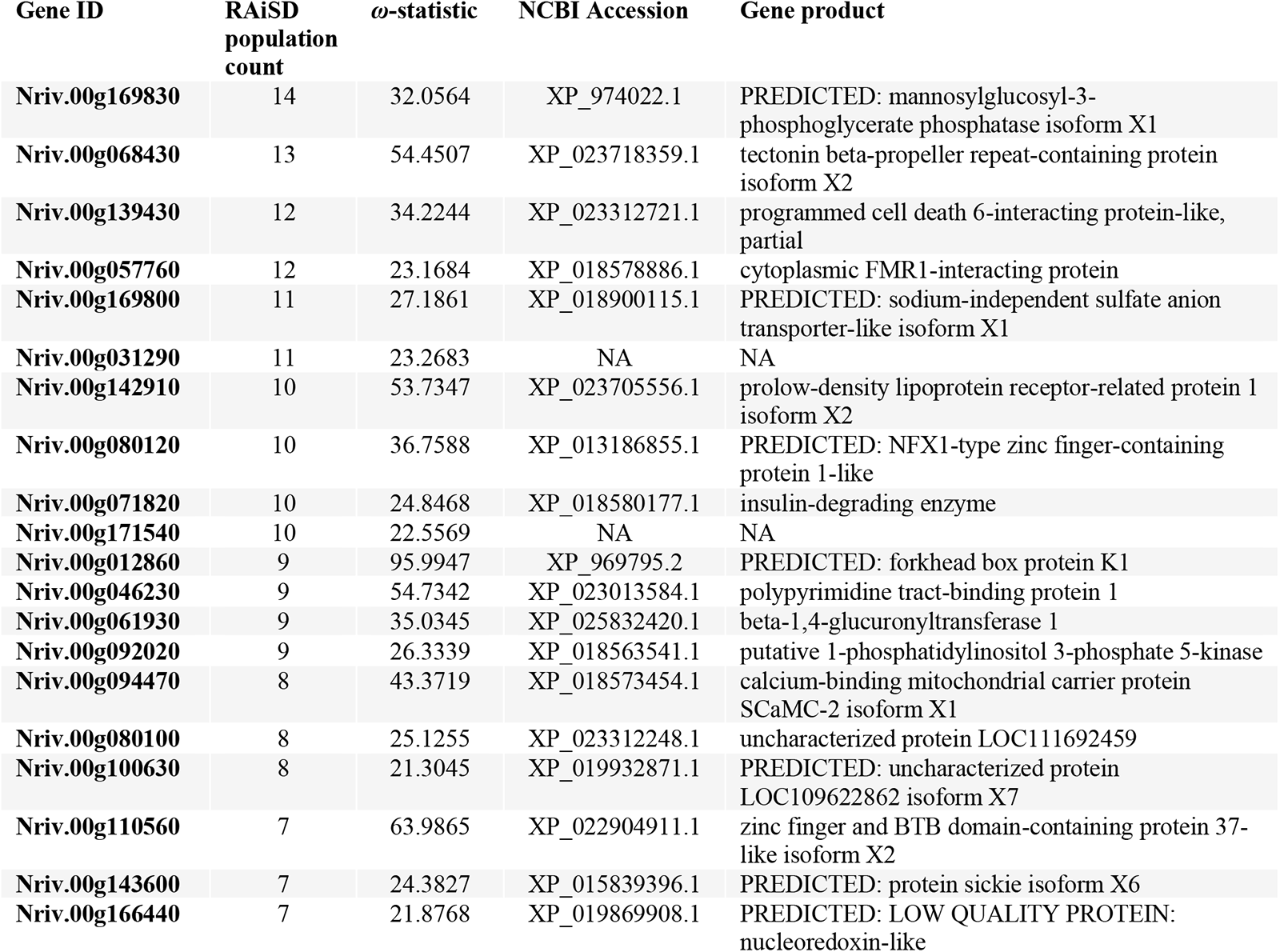
The top 20 genes from the intersection of OmegaPlus and RAiSD genome scans.

### Genotype-environment association (GEA)

We used three environmental variables to scan the genome for genetic variation associated with the environment. Using elevation as a predictor variable, we found 90 SNPs (with Benjamini-Hochberg FDR-adjusted p-value below 0.05) corresponding to 18 genes (13 with gene annotations, **Table 2**). These genes include *daam1* gene (disheveled-associated activator of morphogenesis 1 isoform X2) and *them6* gene (Thioesterase Superfamily Member 6), which are known to be associated with anoxia or hypoxia. A total of 56 GO terms are enriched for elevation (**supplementary file 1: Table S3**), and notable biological process terms include fatty acid metabolism. Using precipitation as a predictor, 15 SNPs corresponding to nine genes are identified (six with gene annotations, **Table 2**). These genes include *tmprss9* (transmembrane protease serine 9) and *bdh1* (D-beta-hydroxybutyrate dehydrogenase, mitochondrial). Insulin catabolic process is a notable biological process enriched for precipitation (**supplementary file 1: Table S4**). Using snow water equivalent as a predictor, 38 significant SNPs and 18 genes were identified (**Table 2**), including *tmprss9*, *AR* (aldose reductase), *futsch* (microtubule-associated protein futsch), *eml2* (echinoderm microtubule-associated protein-like 2), and *Tret1-1* (facilitated trehalose transporter Tret1-like isoform X1). Enriched biological process terms include D-glucuronate catabolic process, microtubule polymerization or depolymerization, and regulation of insulin-like growth factor receptor signaling pathway **(supplementary file 1: Table S5**).

**Table 2.** The list of annotated genes significantly associated with three environmental variables: elevation, precipitation, and snow water equivalent (SWE).

| Gene ID | Environmental variables | Mean adjusted p-value | SNP count | NCBI Accession | Gene products |
| --- | --- | --- | --- | --- | --- |
| Nriv.00g009080 | elevation | 0.039 | 1 | XP_008193114.1 | PREDICTED: uncharacterized protein LOC103312947 (Death domain-containing protein) |
| Nriv.00g015490 | elevation | 0.011 | 1 | XP_019880259.1 | PREDICTED: uncharacterized protein LOC109608276 isoform X2 (immunoglobulin domain-containing protein) |
| Nriv.00g058330 | elevation | 0.040 | 1 | XP_015120571.1 | extensin-like (Chitin_bind_4 domain-containing protein) |
| Nriv.00g060580 | elevation | 0.040 | 3 | XP_022902329.1 | arfaptin-2 |
| Nriv.00g061220 | elevation | 0.038 | 2 | XP_017784081.1 | PREDICTED: hemicentin-1-like isoform X2 |
| Nriv.00g062320 | elevation | 0.011 | 2 | XP_025829314.1 | protein containing domains PDZ_signaling, SH3_DLG-like, and GMPK |
| Nriv.00g062580 | elevation | 0.037 | 1 | XP_008191896.1 | PREDICTED: uncharacterized protein LOC100141693 isoform X3 |
| Nriv.00g072950 | elevation | 0.025 | 1 | XP_012263236.1 | protein THEM6 (thioesterase family protein) |
| Nriv.00g107460 | elevation | 0.018 | 3 | XP_023028942.1 | interferon-inducible double-stranded RNA-dependent protein kinase activator A homolog |
| Nriv.00g124950 | elevation | 0.014 | 3 | XP_018568846.1 | Phospholipase A2 domain-containing protein |
| Nriv.00g129880 | elevation | 0.013 | 2 | XP_017768744.1 | PREDICTED: sorting nexin-13-like isoform X2 |
| Nriv.00g150480 | elevation | 0.017 | 1 | XP_008192803.1 | PREDICTED: disheveled-associated activator of morphogenesis 1 isoform X2 |
| Nriv.00g175620 | elevation | 0.016 | 1 | XP_011860868.1 | PREDICTED: uncharacterized protein LOC105558006 |
| Nriv.00g010710 | precipitation | 0.037 | 1 | XP_018574173.1 | Kruppel-like factor 9 |
| Nriv.00g019110 | precipitation | 0.027 | 2 | XP_001657536.2 | transmembrane protease serine 9 |
| Nriv.00g024510 | precipitation | 0.027 | 1 | XP_023710952.1 | thioredoxin domain-containing protein |
| Nriv.00g025330 | precipitation | 0.005 | 1 | XP_015833364.1 | PREDICTED: chaoptin isoform X2 |
| Nriv.00g066430 | precipitation | 0.024 | 1 | XP_017784383.1 | PREDICTED: D-beta-hydroxybutyrate dehydrogenase, mitochondrial |
| Nriv.00g105190 | precipitation | 0.027 | 1 | XP_015840555.1 | PREDICTED: sushi, von Willebrand factor type A, EGF and pentraxin domain-containing protein 1 isoform X1 |
| Nriv.00g004720 | SWE | 0.011 | 1 | XP_017768153.1 | PREDICTED: actin-binding LIM protein 3-like isoform X3 |
| Nriv.00g015300 | SWE | 0.032 | 1 | XP_022916314.1 | serine/threonine-protein kinase NLK isoform X2 |
| Nriv.00g019110 | SWE | 0.012 | 2 | XP_001657536.2 | transmembrane protease serine 9 |
| Nriv.00g027990 | SWE | 0.017 | 1 | XP_015836986.1 | PREDICTED: putative ferric-chelate reductase 1 homolog |
| Nriv.00g032820 | SWE | 0.030 | 1 | XP_023311431.1 | single-minded homolog 1 |
| Nriv.00g063250 | SWE | 0.003 | 3 | XP_018334317.1 | cancer-related nucleoside-triphosphatase homolog |
| Nriv.00g063520 | SWE | 0.020 | 1 | XP_023020026.1 | thrombospondin type-1 domain-containing protein 4-like |
| Nriv.00g086830 | SWE | 0.021 | 2 | XP_021916483.1 | centrosomal protein of 135 kDa isoform X2 |
| Nriv.00g091800 | SWE | 0.045 | 1 | XP_008549770.1 | PREDICTED: aldose reductase |
| Nriv.00g098910 | SWE | 0.014 | 1 | XP_008201097.1 | PREDICTED: microtubule-associated protein futsch |
| Nriv.00g114090 | SWE | 0.031 | 2 | XP_018579563.1 | echinoderm microtubule-associated protein-like 2 |
| Nriv.00g116540 | SWE | 0.013 | 1 | XP_019867355.1 | PREDICTED: neurotrimin-like isoform X2 |
| Nriv.00g117380 | SWE | 0.012 | 2 | XP_008201486.1 | PREDICTED: protein groucho-1 isoform X2 |
| Nriv.00g133540 | SWE | 0.029 | 1 | XP_017466355.1 | PREDICTED: myb-like protein I |
| Nriv.00g135210 | SWE | 0.036 | 1 | XP_022907726.1 | facilitated trehalose transporter Tret1-like isoform X1 |

## Discussion

With advances in genome sequencing, non-model species are increasingly being studied to identify the genetic basis of adaptation to local environments (Hoban et al. 2016). Genes showing strong evidence for a change in allele frequency above neutral patterns (signatures of a selective sweep or strong environmental association) are candidates for adaptation (Le Corre and Kremer 2012; Barghi and Schlötterer 2020). However, for species experiencing drastic demographic changes such as rapid distributional range shifts, signals of selection in the genome are difficult to disentangle from patterns of genetic drift (Szpiech et al. 2021). In this study, we used whole genome resequencing data to investigate genome-wide patterns of adaptation in an alpine ground beetle, the *N. ingens* complex, following rapid distributional shifts to alpine habitats induced by post-glacial climate change. We first used the genomic data to reconstruct the demographic history including the formation of hybrid populations from an early (∼0.8 mya) admixture event (**Figure 1**), then reinforced previous results showing that genetic structure and distributional models are consistent with glacial refugia in the lower elevation areas of the Sierra Nevada. We then employed whole genome scans to understand how species adapt to novel alpine conditions despite strong demographic change. As phenotypic traits involved in adaption are often polygenic, it can be challenging to detect subtle changes in allele frequency across many loci while controlling for false positive rates (Pritchard and Di Rienzo 2010; Le Corre and Kremer 2012; Rees et al. 2020). Enrichment analyses of gene functions can partially resolve the problem of identifying polygenic adaptation (Fagny and Austerlitz 2021), although many genes identified from non-model organisms are not well annotated for their species-specific function. As a result, uncertainties about false positives accentuate the importance of using a strong hypothesis-testing framework to identify putative candidate genes in the context of evolution following rapid environmental change. The series of paleodistribution models demonstrate an increasing elevational distribution (>1500 meters) approaching present day conditions, which leads to the hypothesis that populations experienced strong environmental selection. Specifically, alpine environments are expected to exert strong selection on regulation of metabolism, hypoxia, and desiccation resistance. We tested for evidence of selection in these pathways using whole genome scan approaches and conservative testing procedures, finding strong evidence for known genes that are associated with these physiological stressors.

### The demographic history of <u>N. ingens</u> complex and its implications for climate adaptation

It is well known that glacial cycles have shaped the distribution of genetic variation in alpine species and lead to allopatric diversification, but few studies have examined population genomic resequencing detail in fine geographical resolution (Boucher et al. 2016; Sim et al. 2016; Weng et al. 2016; Huang et al. 2017; Rovito and Schoville 2017; Capblancq et al. 2019; He et al. 2019; Pan et al. 2020; Weng et al. 2020; Ortego and Knowles 2022). Here, our work shows a dynamic pattern of spatial recolonization, with a more ancient episode of lineage formation following population admixture. In previous work, mitochondrial and morphological data suggested the possible presence of hybrid populations (Schoville et al. 2012), and SNP data suggested that the current genetic structure of the *N. ingens* complex could be explained by distributional shifts within drainage basins during the last glacial maximum (Weng et al. 2021a). In this study, whole genome resequencing clearly shows that species known as *N. riversi* and *N. ingens* diverged 1.06 mya (0.79-1.32 mya) and hybridized around 0.8 mya (0.69-0.91 mya) to form a third distinct lineage. This admixture event coincides with the timing of the most extensive Pleistocene glacial event in the Sierra Nevada, the Sherwin glaciation (ca. 0.82 mya) (Moore and Moring 2013), suggesting that major alpine glacial events facilitate low-elevation gene flow of alpine organisms across the Sierra Nevada. The Sherwin glaciation also resulted in a gene flow event for the sub-alpine Foxtail Pine, *Pinus balfouriana*, between the Sierra Nevada and the northern California Trinity Mountains (Eckert et al. 2008).

Furthermore, simulations coupled to paleoclimate models show how an increase in elevational distribution and a reduction in suitable habitat since the last glacial maximum drove rapid spatial shifts accompanied by population declines. The result is supported by our previous finding that the effective population size of this species complex has declined since the last glacial maximum (Weng et al. 2021a).

### Testing for alpine adaptation following postglacial recolonization

A distributional range shift to higher elevation habitats following retreating glaciers involves novel alpine environmental conditions that could exert strong selective pressure. For nocturnally foraging beetles, the extreme environment in the alpine zone creates adverse conditions that include reduced atmospheric oxygen and low water vapor (arid conditions). Furthermore, alpine microhabitats of the *N. ingens* complex vary considerably across the span of Sierra Nevada, where microhabitats of the northern species, *N. riversi*, are covered with more snow for longer periods of time but occur at lower elevations on average compared to southern populations of *N. ingens*. To test for adaptation to the alpine environment, we employed genome scans of selection to identify signatures of selective sweeps that are shared across populations, and we also examined how genetic variation is statistically associated with three microhabitat variables, snow water equivalent (SWE), precipitation, and elevation. For elevation, we expected candidate genes to be linked to the reduced atmospheric oxygen (hypoxic conditions). For SWE and precipitation, selection to improve desiccation resistance is expected for populations in more arid regions.

Based on shared signatures of selective sweeps across populations, we identified 162 candidate genes using conservative testing criteria. The enriched functions from these genes include muscle and neuron development, regulation of growth, insulin-mediated metabolic pathways, response to stress, regulation of homeostasis, inorganic cation transmembrane transport, and sperm viability (**supplementary file 2: WGS enriched BP terms**). The biological terms of muscle and neuron development are mostly from the three candidate genes: *LRP1* (prolow-density lipoprotein receptor-related protein 1 isoform X2), *msn* (Non-specific serine/threonine protein kinase) and *PAK3* (serine/threonine-protein kinase PAK 3). Additionally, these two serine/threonine protein kinase genes together with *IDE* (insulin-degrading enzyme) provide more specific connections to the regulation of metabolic rate through insulin catabolism. The insulin-mediated metabolic pathways have been found to respond to hypoxic conditions, including in alpine locusts in Tibet where it maintains high aerobic respiration (Zhang et al. 2013; Zhao et al. 2013; Ding et al. 2018). It is also noteworthy that the putative hypoxia gene, *COX* (cytochrome c oxidase assembly protein COX15 homolog) was identified from nine of the 21 populations in RAiSD, suggesting widespread convergence across populations. Among the 18 elevation-associated genes, *DAAM1*, *THEM6*, and *PLA2* (Phospholipase A2 domain-containing protein) are potentially linked to hypoxia adaptation (Zhang et al. 2019; Fang et al. 2020). In addition, the enriched biological process terms from these 18 genes comprise functions in fatty acid and lipoprotein metabolism. Thus, selection from environmental stressors appears to drive alpine adaptation.

A second important environmental challenge in alpine habitats is the decrease in water vapor (humidity) as elevation increases and temperature declines to near freezing. Drier air coupled with a windier alpine environment could cause desiccation stress (Sømme 1989; Sinclair 2000b; Dillon et al. 2006). The physiological response to desiccation stress in insects generally involves the regulation of cuticular hydrocarbon composition, prevention of protein denaturation (e.g. late embryogenesis abundant proteins and heat shock proteins), and the maintenance of homeostats through trehalose, mannitol, and glycerol metabolism (Clark et al. 2009; Cornette et al. 2010; Storey and Storey 2012; Nesmelov et al. 2018; Thorat and Nath 2018). Among the genes identified as selective sweeps, we found enriched terms of cellular chemical homeostasis and inorganic cation transmembrane transport that may be linked to desiccation-resistant phenotypes. We also examined genetic variation associated with SWE and precipitation. Among the 18 SWE associated genes, *Tret1-1* gene is most likely to play a part in adaptation to desiccation stress as it has been reported to regulate trehalose levels in the hemolymph against desiccation, cold, and other stressors (Kikawada et al. 2007; Kanamori et al. 2010; Kikuta et al. 2012). However, the *Tret1-1* gene may alternatively be selected by temperature due to its well-known cryoprotectant function (Uchida et al. 2019; Zhao et al. 2021). More generally, many enriched terms from our genome scans are associated with stress adaptation. For example, other SWE associated genes, such as *futsch, eml2,* and *AR* gene (aldose reductase), could be involved in cold acclimation, as, for example, *AR* functions in converting glucose to sorbitol in the polyol pathway (Storey 1990; Joanisse and Storey 1994; Wolfe et al. 1998; Teets et al. 2012; Yamamoto et al. 2020). The nine precipitation associated genes have no clear connection to desiccation resistance, and instead are mostly linked to tissue and organ development. Interestingly, the *tmprss9* gene was identified with both SWE and precipitation. This gene has enriched functions of dorsal/ventral pattern formation and insulin catabolic process, but how these functions might link to desiccation is unclear.

## Conclusions

In summary, we found evidence that populations of the *N. ingens* complex have responded to glacial cycles in a dynamic pattern of spatial movement that has allowed for occasional admixture, but has also reduced population size and generated complex spatial genetic structure. Shifts to alpine conditions have involved rapid spatial shifts and large changes in elevation (>1500 meters), and indeed, we see clear evidence that selection to alpine conditions acts on hypoxia-related pathways. Although selection scans also identified genes that are broadly associated with a range of stress responses, clear functional evidence for the drivers of selection on these pathways is needed to improve our understanding of alpine adaptation in the *N. ingens* complex.

## Materials and Methods

### Study System, Sampling and Sequencing

A total of 384 adult beetles were sampled with permits granted by Sequoia and Kings National Parks (Study# SEKI-0091), Yosemite National Park (Study# YOSE-00093), and the California Fish and Game Department (#SC-006997). Details of sample sizes and localities were described in (Weng et al, 2021a). Genomic DNA was extracted from the thoracic muscle tissue of each individual using either the QIAamp 96 DNA Blood Kit or the DNeasy Blood & Tissue Kits (Qiagen, LLC). Libraries were prepared for low-coverage whole genome sequencing (2×150 bp reads) using an Illumina Nextera® DNA Library Preparation Kit by the Biotechnology Center of University of Wisconsin-Madison. Each sample was sequenced on the Illumina Novaseq 6000 to moderate coverage (∼5x).

### Variant Calling

We examined the quality of sequencing read data using FASTQC (Andrews 2010) and trimmed low quality sequences, adaptors, and putative contaminants based on the UniVec database from NCBI (www.ncbi.nlm.nih.gov/VecScreenUniVec.html) using the BBduk function in the BBtools package (Bushnell 2014). The trimmed reads were subsequently mapped to a high-quality reference genome of *N. riversi* using BWA-mem (Li 2013; Weng et al. 2021b). The mapped reads were then extracted and sorted with Samtools (Li et al. 2009). To call the variants from the mapped reads, we followed the GATK pipeline (McKenna et al. 2010; DePristo et al. 2011; Van der Auwera et al. 2013; Poplin et al. 2017). Briefly, we employed HaplotypeCaller and GenotypeGVCFs to generate a raw cohort SNP callset (**supplementary file 1: Figure S2**; details of variant calling process are described in the **supplementary file 1: Supplementary Methods**. For these raw variants, we evaluated multiple combinations of base-recalibration and variant-filtering procedures to minimize the genotype calling error, by comparing the genotype and genotype dosage for one individual with high coverage sequence data (171.4x coverage; data is available on NCBI with SRA number SRR12215150) to the genotypes generated from low coverage sequencing of the same individual. Genotypes of this individual were called from the high coverage data using a strict hard-filtered process implemented in the GATK variant calling pipeline (**supplementary file 1: Figure S2**), as a nearly 100% accuracy is expected for genotype calls when ultra-deep whole-genome sequencing data is available (Kishikawa et al. 2019). This strict hard-filtered raw cohort SNP callset, as well as a cohort SNP callset from GBS data (19.2x coverage) published by Weng *et al*. (2021), was used as a training dataset for Variant Quality Score Recalibration (VQSR). We employed VQSR at 99.9% target sensitivity, as it retains more SNPs with fewer genotype mismatches (**supplementary file 1: Table S2**). As we had large sample sizes at each collection site, we subsequently imputed missing genotypes using BEAGLE 4.1 (Browning and Browning 2016), after dropping samples with > 90% missing data (**supplementary file 1: Figure S4**). Finally, we applied a filter to trim the loci with a minor allele frequency less than 0.05 for downstream analyses.

### Genetic structure of the <u>N. ingens</u> complex

To investigate the genetic structure of the *N. ingens* species complex with the genome-wide SNP markers, we first examined the decay of linkage disequilibrium (LD) by calculating pairwise correlations between SNPs within a range of 1 Mb using *plink* v1.90b6.21 (Zheng et al. 2012). Since the reference genome is not a chromosomal level assembly, we calculated the LD decay for each of the 1,345 putative autosomal contigs (determined by the male-female ratio of mean contig read coverage, **supplementary file 1: Figure S5**) that are larger than 10 kbps and ignored inter-contig LD. The genome-wide LD decay is then represented by the average LD decay across the contigs (**supplementary file 1: Figure S6**; see **Supplementary Methods** for more detail), and shows that the mean LD decays within 5 Kb. We used the snpgdsLDpruning function in the R package *SNPRelate* (Zheng et al. 2012) to remove SNP loci with *R*^2^ higher than 0.5 in this 5 Kb range. The resulting dataset (378,532 SNPs) was used to estimate genetic population structure with principal component analysis (PCA) and sparse non-negative matrix factorization (sNMF). For PCA, we used the function snpgdsPCA in *SNPRelate* to visualize the genetic structure for the first 15 principal components (PCs). For sNMF, we used function snmf in the R package *LEA* to calculate the ancestry coefficients for three to twelve clusters (K), with 10 replicates each (Frichot et al. 2014; Frichot and François 2015). In addition to the clustering analyses, we calculated heterozygosity of each population and the pairwise *F*_ST_ and *d*_XY_ (*π*_XY_ as it was originally described) to describe the genetic structure using the R package *PopGenome* (Nei and Li 1979; Pfeifer et al. 2014).

### Testing Admixture in Lineage Formation

The intermediate lineage of the *N. ingens* species complex might represent an ancestral hybridization event of *N. riversi* and *N. ingens*, as beetles in this lineage show intermediate color pattern and genetic evidence of admixture (Schoville et al. 2012; Weng et al. 2021a). To identify the origin of this intermediate lineage, we performed ABC analysis to compare two competing evolutionary models. An isolation model (three independent lineages: *N. riversi*, intermediate, and *N. ingens*) was proposed as a null model, as it assumes that the intermediate population arose as the result of genetic divergence with limited gene flow. As an alternative, an admixture model depicts the intermediate lineage forming as the result of an admixture event between *N. riversi* and *N. ingens* (**Figure 1**). To simplify the simulations, we selected three populations representing each lineage (Conness, Selden, and Army), based on their isolation and lack of recent gene flow (Weng et al. 2021a; Lamigueiro et al. 2023). The presence of recent gene flow could confound the ABC analysis by biasing the summary statistics towards the admixture model, as well as skew parameter estimates. The details of the priors and other parameters used to simulate data in *msprime* v. 1.0.4 are described in the **Supplementary Methods** (Kelleher et al. 2016). Notably, we set the timing of admixture to a wide range (from ca. 115 to 960 kyr) to allow for admixture over several glacial episodes. Model selection was performed using the R package “abc” (Csilléry et al. 2012) based on two summary statistics: the outgroup-*f*3 statistic and absolute pairwise genetic divergence (*d*_XY_) (Nei and Li 1979; Patterson et al. 2012). These summary statistics were chosen to efficiently capture information that can distinguish the independent lineage model from the admixture model with minimum complexity. For the summary statistics from the simulation data, we used tskit v. 2.0.2 to calculate the outgroup-*f*3 statistic and pairwise *d*_XY_ statistics (Kelleher et al. 2018), and for the observed data, we calculated the outgroup-*f*3 statistic using Treemix and pairwise *d*_XY_ statistics with the R package “PopGenome” (Pickrell and Pritchard 2012; Pfeifer et al. 2014). Both statistics were calculated using the maf-filtered SNPs with window size of 10,000 base pairs.

### Recolonization following the last glacial maximum

To reconstruct the process of post-glacial colonization, we generated raster maps of the Sierra Nevada Mountains using areas higher than 1,000 meters in elevation (this range spans 38 to 40 degrees latitude, -118 to -120 longitude). In order to generate shifting habitat suitability through time, species distribution models were generated from the last glacial maximum (LGM, 21,000 yr) to present day in seven time periods, using paleoclimatic models downloaded from the PaleoClim database (Dolan et al. 2015; Hill 2015; Brown et al. 2018; Karger and Zimmermann 2019), and a contemporary climatic model (1970-2000 AD) from the WorldClim database (Fick and Hijmans 2017; Karger and Zimmermann 2019). Modern beetle occurrence data (45 sites) was used to generate a modern SDM using the R package ENMeval v2.0.3 (Kass et al. 2021), and the suitable range of climate variables was then used to fit each paleoclimate model. To reduce overprediction of suitable habitat at the earliest time interval, we applied the LGM glacier boundary to the SDM to remove suitable patches within the boundary of the glacial ice sheet (Rood et al. 2011), as the environment within this range was extremely harsh and unlikely to provide habitat for beetles (**supplementary file 1: Figure S11**). The aforementioned SDMs were selected based on the minimum AICc using “ENMevaluate” in the R package ENMeval v2.0.3 (Kass et al. 2021). Details of procedures used to generate species distribution models are described in the **Supplementary Methods**.

### Genome-wide scan for selective sweeps

To identify the genes involved in adaptation following post-glacial colonization, we used two genome scan approaches to detect selective sweeps. Each method relies on different measures of genomic diversity, and by combining them, we reduce the likelihood of false positives (Lotterhos et al. 2017). We first employed OmegaPlus v2.2.2 (Alachiotis et al. 2012) to measure the omega statistic (*ω*), which measures linkage disequilibrium among variants in defined genomic windows (Kim and Nielsen 2004). We measured the *ω*-statistic in genomic windows ranging from 1kb to 10kb (minwin=1,000 and maxwin=10,000) and set the threshold for outlier detection at the 0.995 quantile (top 0.5%) of the *ω*-statistic distribution (**supplementary file 1: Figure S13**). SNPs were filtered to remove contigs smaller than 10kbp, multiallelic variants, and loci with maf less than 5%. This method has been shown to perform well as a global measure of selection in species with structured populations (Crisci et al. 2013; Montano et al. 2015). Second, using the same filtered SNP dataset, we employed the method RAiSD (Alachiotis and Pavlidis 2018). This approach detects putative selective sweeps by calculating the *μ*-statistic, a composite likelihood that measures genetic diversity, the site allele frequency, and linkage disequilibrium. Since the *μ*-statistic takes the site allele frequency into account, we performed this analysis on individual populations (we ignore populations with fewer than five beetles sampled) to avoid any confounding effect of population structure (**supplementary file 1: Table S1**), as the introduction of foreign alleles could increase the proportion of minor alleles and skew the site allele frequency spectrum to the left. The outlier regions from each population were defined to be the 0.995 quantile of the scanned SNPs (Quinlan and Hall 2010). Outlier regions from both methods were further thinned to a candidate gene set based on whether they intersected annotated genes. We used bedtools v2.30.0 (Quinlan and Hall 2010) to generate these intersections. Finally, we found the intersection of outliers from each method to generate the most reliable set of candidate selective sweeps.

### Genotype-environment association (GEA)

To investigate genome-wide signatures of adaptation to the local environment, we conducted genotype-environment association (GEA) analysis on three uncorrelated environmental factors that relate to desiccation stress and hypoxia. Higher levels of snow water equivalent (SWE) and precipitation are important measures of water availability in nival habitats, and this species complex relies on flowing water throughout the summer. For the SWE data (representing daily measures across 31 years, 1985-2015; (Margulis et al. 2016)), we chose to calculate the mean for the month of May (**supplementary file 1: Figure S15**) because SWE typically peaks at this time and overwintering adult beetles do not become active until late May. Precipitation data were downloaded from the WorldClim global climate database (Fick and Hijmans 2017). Finally, we used elevation from the same database as it relates directly to atmospheric oxygen content and hypoxic stress.

To detect environmentally associated genes, we used a latent factor mixed modeling approach implemented in LFMM2 (Caye et al. 2019) to control for population structure while testing for environmental associations with the three environmental factors. The “maf” dataset was used and the number of latent factors (K=10) was determined by examining the scree plot of a PCA. Subsequently, we applied ridge and lasso regression methods to control model complexity during statistical tests. We selected candidates as those SNPs below a Benjamini-Hochberg adjusted p-value = 0.05 in both ridge and lasso regressions, and further focused on those SNPs that intersected annotated genes.

### Gene ontology (GO) enrichment

For all candidate genes of selection, we performed GO enrichment analysis using the R package clusterProfiler (Yu et al. 2012) v3.16.1. We used a *p*-value threshold of 0.01 in enrichment analyses for the gene sets from both genomic scans and GEA to all three environmental variables. The GO terms of the genes were extracted from the annotated genome assembly of *N. riversi* (Weng et al. 2021b).

## Data Availability Statement

The data underlying this article are publicly available at NCBI (Bioproject PRJNA937745) and as Supplementary Material Additional File 1, 2, and 3.

## Acknowledgments and Funding Information

We are grateful to Yosemite, and Sequoia and Kings Canyon National Parks for providing permission to conduct research in the parks. Funding was provided by the University of Wisconsin-Madison to SDS and the Ministry of Education Republic of China (Taiwan) to YMW. This research is also supported by a grant from the Valentine Eastern Sierra Reserve (UC Santa Barbara Natural Reserve System) to YMW. We thank Benton Veire and Matthew Medeiros for helping us to conduct the field work, and the Ph.D. committee of YMW for providing guidance. The authors have no conflicts of interests to declare.

## References

Alachiotis N, Pavlidis P. 2018. RAiSD detects positive selection based on multiple signatures of a selective sweep and SNP vectors. Communications Biology 1:79.

Alachiotis N, Stamatakis A, Pavlidis P. 2012. OmegaPlus: a scalable tool for rapid detection of selective sweeps in whole-genome datasets. Bioinformatics 28:2274–2275.

Andrews S. 2010. FastQC: a quality control tool for high throughput sequence data. In: Babraham Bioinformatics, Babraham Institute, Cambridge, United Kingdom.

Barghi N, Schlötterer C. 2020. Distinct Patterns of Selective Sweep and Polygenic Adaptation in Evolve and Resequence Studies. Genome Biology and Evolution 12:890–904.

Barghi N, Tobler R, Nolte V, Jakšić AM, Mallard F, Otte KA, Dolezal M, Taus T, Kofler R, Schlötterer C. 2019. Genetic redundancy fuels polygenic adaptation in Drosophila. PLoS Biology 17:e3000128.

Bell G, Collins S. 2008. Adaptation, extinction and global change. Evolutionary Applications 1:3–16.

Bohutínská M, Vlček J, Yair S, Laenen B, Konečná V, Fracassetti M, Slotte T, Kolář F. 2021. Genomic basis of parallel adaptation varies with divergence in *Arabidopsis* and its relatives. Proceedings of the National Academy of Sciences 118.

Boucher FC, Zimmermann NE, Conti E. 2016. Allopatric speciation with little niche divergence is common among alpine Primulaceae. Journal of Biogeography 43:591–602.

Brown JL, Hill DJ, Dolan AM, Carnaval AC, Haywood AM. 2018. PaleoClim, high spatial resolution paleoclimate surfaces for global land areas. Scientific Data 5:180254.

Browning BL, Browning SR. 2016. Genotype Imputation with Millions of Reference Samples. The American Journal of Human Genetics 98:116–126.

Buffalo V. 2021. Quantifying the relationship between genetic diversity and population size suggests natural selection cannot explain Lewontin’s paradox. Elife 10.

Bushnell B. 2014. BBTools software package. URL http://sourceforge.net/projects/bbmap.

Capblancq T, Mavárez J, Rioux D, Després L. 2019. Speciation with gene flow: Evidence from a complex of alpine butterflies (*Coenonympha*, Satyridae). Ecology and Evolution 9:6444–6457.

Caye K, Jumentier B, Lepeule J, François O. 2019. LFMM 2: Fast and Accurate Inference of Gene-Environment Associations in Genome-Wide Studies. Molecular Biology and Evolution 36:852–860.

Cheng Y, Miller MJ, Zhang D, Xiong Y, Hao Y, Jia C, Cai T, Li SH, Johansson US, Liu Y et al. 2021. Parallel genomic responses to historical climate change and high elevation in East Asian songbirds. Proceedings of the National Academy of Sciences 118.

Cheviron ZA, Bachman GC, Connaty AD, McClelland GB, Storz JF. 2012. Regulatory changes contribute to the adaptive enhancement of thermogenic capacity in high-altitude deer mice. Proceedings of the National Academy of Sciences 109:8635–8640.

Clark MS, Thorne MA, Purać J, Burns G, Hillyard G, Popović ZD, Grubor-Lajsić G, Worland MR. 2009. Surviving the cold: molecular analyses of insect cryoprotective dehydration in the Arctic springtail Megaphorura arctica (Tullberg). BMC Genomics 10:328.

Cornette R, Kanamori Y, Watanabe M, Nakahara Y, Gusev O, Mitsumasu K, Kadono-Okuda K, Shimomura M, Mita K, Kikawada T et al. 2010. Identification of anhydrobiosis-related genes from an expressed sequence tag database in the cryptobiotic midge Polypedilum vanderplanki (Diptera; Chironomidae). Journal of Biological Chemistry 285:35889–35899.

Crisci JL, Poh YP, Mahajan S, Jensen JD. 2013. The impact of equilibrium assumptions on tests of selection. Frontiers in genetics 4:235.

Cruickshank TE, Hahn MW. 2014. Reanalysis suggests that genomic islands of speciation are due to reduced diversity, not reduced gene flow. Molecular Ecology 23:3133–3157.

Csilléry K, François O, Blum MGB. 2012. abc: an R package for approximate Bayesian computation (ABC). Methods in Ecology and Evolution 3:475–479.

DePristo MA, Banks E, Poplin R, Garimella KV, Maguire JR, Hartl C, Philippakis AA, del Angel G, Rivas MA, Hanna M et al. 2011. A framework for variation discovery and genotyping using next-generation DNA sequencing data. Nat Genet 43:491–498.

Dillon ME, Frazier MR, Dudley R. 2006. Into thin air: physiology and evolution of alpine insects. Integrative and Comparative Biology 46:49–61.

Ding D, Liu G, Hou L, Gui W, Chen B, Kang L. 2018. Genetic variation in PTPN1 contributes to metabolic adaptation to high-altitude hypoxia in Tibetan migratory locusts. Nature Communications 9:4991.

Dolan AM, Haywood AM, Hunter SJ, Tindall JC, Dowsett HJ, Hill DJ, Pickering SJ. 2015. Modelling the enigmatic late Pliocene glacial event—Marine Isotope Stage M2. Global and Planetary Change 128:47–60.

Eckert AJ, Tearse BR, Hall BD. 2008. A phylogeographical analysis of the range disjunction for foxtail pine (Pinus balfouriana, Pinaceae): the role of Pleistocene glaciation. Molecular Ecology 17:1983–1997.

Ellegren H, Galtier N. 2016. Determinants of genetic diversity. Nature Reviews Genetics 17:422–433.

Excoffier L, Foll M, Petit RJ. 2009. Genetic Consequences of Range Expansions. Annual Review of Ecology, Evolution, and Systematics 40:481–501.

Fagny M, Austerlitz F. 2021. Polygenic Adaptation: Integrating Population Genetics and Gene Regulatory Networks. Trends in Genetics 37:631–638.

Fang X, Zhang D, Zhao W, Gao L, Wang L. 2020. Dishevelled Associated Activator Of Morphogenesis (DAAM) Facilitates Invasion of Hepatocellular Carcinoma by Upregulating Hypoxia-Inducible Factor 1α (HIF-1α) Expression. Medical Science Monitor: International Medical Journal of Experimental and Clinical Research 26:e924670–924671.

Fick SE, Hijmans RJ. 2017. WorldClim 2: new 1-km spatial resolution climate surfaces for global land areas. International journal of climatology 37:4302–4315.

Foll M, Gaggiotti OE, Daub JT, Vatsiou A, Excoffier L. 2014. Widespread signals of convergent adaptation to high altitude in Asia and america. The American Journal of Human Genetics 95:394–407.

Frichot E, François O. 2015. LEA: An R package for landscape and ecological association studies. Methods in Ecology and Evolution 6:925–929.

Frichot E, Mathieu F, Trouillon T, Bouchard G, François O. 2014. Fast and efficient estimation of individual ancestry coefficients. Genetics 196:973–983.

Haller BC, Messer PW. 2019. SLiM 3: Forward genetic simulations beyond the Wright–Fisher model. Molecular biology and evolution 36:632–637.

Harrison JF, Greenlee KJ, Verberk WCEP. 2018. Functional Hypoxia in Insects: Definition, Assessment, and Consequences for Physiology, Ecology, and Evolution. Annual Review of Entomology 63:303–325.

He L, Gomes AP, Wang X, Yoon SO, Lee G, Nagiec MJ, Cho S, Chavez A, Islam T, Yu Y. 2018. mTORC1 promotes metabolic reprogramming by the suppression of GSK3-dependent Foxk1 phosphorylation. Molecular cell 70:949–960. e944.

He X, Burgess KS, Gao LM, Li DZ. 2019. Distributional responses to climate change for alpine species of *Cyananthus* and *Primula* endemic to the Himalaya-Hengduan Mountains. Plant Divers 41:26–32.

Henry JR, Harrison JF. 2004. Plastic and evolved responses of larval tracheae and mass to varying atmospheric oxygen content in *Drosophila melanogaster*. Journal of Experimental Biology 207:3559–3567.

Hill DJ. 2015. The non-analogue nature of Pliocene temperature gradients. Earth and Planetary Science Letters 425:232–241.

Hoback WW, Stanley DW. 2001. Insects in hypoxia. Journal of Insect Physiology 47:533–542.

Hoban S, Kelley JL, Lotterhos KE, Antolin MF, Bradburd G, Lowry DB, Poss ML, Reed LK, Storfer A, Whitlock MC. 2016. Finding the Genomic Basis of Local Adaptation: Pitfalls, Practical Solutions, and Future Directions. The American Naturalist 188:379–397.

Hof C, Levinsky I, Araujo MB, Rahbek C. 2011. Rethinking species’ ability to cope with rapid climate change. Global Change Biology 17:2987–2990.

Huang ZS, Yu FL, Gong HS, Song YL, Zeng ZG, Zhang Q. 2017. Phylogeographical structure and demographic expansion in the endemic alpine stream salamander (Hynobiidae: Batrachuperus) of the Qinling Mountains. Sci Rep 7:1871.

Jezkova T, Wiens JJ. 2016. Rates of change in climatic niches in plant and animal populations are much slower than projected climate change. Proceedings of the Royal Society B 283.

Joanisse D, Storey KB. 1994. Enzyme activity profiles in an overwintering population of freeze-tolerant larvae of the gall fly, Eurosta solidaginis. Journal of Comparative Physiology B 164:247–255.

Johri P, Charlesworth B, Jensen JD. 2020. Toward an Evolutionarily Appropriate Null Model: Jointly Inferring Demography and Purifying Selection. Genetics 215:173–192.

Jones MR, Mills LS, Jensen JD, Good JM. 2020. The Origin and Spread of Locally Adaptive Seasonal Camouflage in Snowshoe Hares. American Naturalist 196:316–332.

Kanamori Y, Saito A, Hagiwara-Komoda Y, Tanaka D, Mitsumasu K, Kikuta S, Watanabe M, Cornette R, Kikawada T, Okuda T. 2010. The trehalose transporter 1 gene sequence is conserved in insects and encodes proteins with different kinetic properties involved in trehalose import into peripheral tissues. Insect Biochemistry and Molecular Biology 40:30–37.

Karger DN, Zimmermann NE. 2019. Climatologies at high resolution for the earth land surface areas CHELSA V1. 2: Technical specification. Scientific Data.

Kass JM, Muscarella R, Galante PJ, Bohl CL, Pinilla-Buitrago GE, Boria RA, Soley-Guardia M, Anderson RP. 2021. ENMeval 2.0: redesigned for customizable and reproducible modeling of species’ niches and distributions. Methods in Ecology and Evolution.

Kelleher J, Etheridge AM, McVean G. 2016. Efficient Coalescent Simulation and Genealogical Analysis for Large Sample Sizes. PLOS Computational Biology 12:e1004842.

Kelleher J, Thornton KR, Ashander J, Ralph PL. 2018. Efficient pedigree recording for fast population genetics simulation. PLOS Computational Biology 14:e1006581.

Kikawada T, Saito A, Kanamori Y, Nakahara Y, Iwata K, Tanaka D, Watanabe M, Okuda T. 2007. Trehalose transporter 1, a facilitated and high-capacity trehalose transporter, allows exogenous trehalose uptake into cells. Proceedings of the National Academy of Sciences 104:11585–11590.

Kikuta S, Hagiwara-Komoda Y, Noda H, Kikawada T. 2012. A novel member of the trehalose transporter family functions as an h(+)-dependent trehalose transporter in the reabsorption of trehalose in malpighian tubules. Frontiers in Physiology 3:290.

Kim Y, Nielsen R. 2004. Linkage disequilibrium as a signature of selective sweeps. Genetics 167:1513–1524.

Kishikawa T, Momozawa Y, Ozeki T, Mushiroda T, Inohara H, Kamatani Y, Kubo M, Okada Y. 2019. Empirical evaluation of variant calling accuracy using ultra-deep whole-genome sequencing data. Scientific Reports 9:1784.

Lamigueiro OP, Hijmans R, Lamigueiro MOP. 2023. Package ‘rasterVis’.

Le Corre V, Kremer A. 2012. The genetic differentiation at quantitative trait loci under local adaptation. Molecular Ecology 21:1548–1566.

Lee P, Chandel NS, Simon MC. 2020. Cellular adaptation to hypoxia through hypoxia inducible factors and beyond. Nature Reviews Molecular Cell Biology 21:268–283.

Leffler EM, Bullaughey K, Matute DR, Meyer WK, Ségurel L, Venkat A, Andolfatto P, Przeworski M. 2012. Revisiting an old riddle: what determines genetic diversity levels within species? PLoS Biol 10:e1001388.

Li H. 2013. Aligning sequence reads, clone sequences and assembly contigs with BWA-MEM. arXiv preprint arXiv:1303.3997.

Li H, Handsaker B, Wysoker A, Fennell T, Ruan J, Homer N, Marth G, Abecasis G, Durbin R. 2009. The sequence alignment/map format and SAMtools. Bioinformatics 25:2078–2079.

Loarie SR, Duffy PB, Hamilton H, Asner GP, Field CB, Ackerly DD. 2009. The velocity of climate change. Nature 462:1052–1055.

Lotterhos KE, Card DC, Schaal SM, Wang L, Collins C, Verity B. 2017. Composite measures of selection can improve the signal-to-noise ratio in genome scans. Methods in Ecology and Evolution 8:717–727.

Loudon C. 1989. Tracheal hypertrophy in mealworms: design and plasticity in oxygen supply systems. Journal of Experimental Biology 147:217–235.

Margulis SA, Cortés G, Girotto M, Durand M. 2016. A Landsat-Era Sierra Nevada Snow Reanalysis (1985–2015). Journal of Hydrometeorology 17:1203–1221.

McCulloch GA, Guhlin J, Dutoit L, Harrop TWR, Dearden PK, Waters JM. 2021. Genomic signatures of parallel alpine adaptation in recently evolved flightless insects. Molecular Ecology 30:6677–6686.

McKenna A, Hanna M, Banks E, Sivachenko A, Cibulskis K, Kernytsky A, Garimella K, Altshuler D, Gabriel S, Daly M. 2010. The Genome Analysis Toolkit: a MapReduce framework for analyzing next-generation DNA sequencing data. Genome research 20:1297–1303.

Montano V, Didelot X, Foll M, Linz B, Reinhardt R, Suerbaum S, Moodley Y, Jensen JD. 2015. Worldwide Population Structure, Long-Term Demography, and Local Adaptation of Helicobacter pylori. Genetics 200:947–963.

Moore JG, Moring BC. 2013. Rangewide glaciation in the Sierra Nevada, California. Geosphere 9:1804–1818.

Nei M, Li WH. 1979. Mathematical model for studying genetic variation in terms of restriction endonucleases. Proceedings of the National Academy of Sciences 76:5269–5273.

Nesmelov A, Cornette R, Gusev O, Kikawada T. 2018. The Antioxidant System in the Anhydrobiotic Midge as an Essential, Adaptive Mechanism for Desiccation Survival. Advances in Experimental Medicine and Biology 1081:259–270.

Orr HA, Unckless RL. 2008. Population extinction and the genetics of adaptation. American Naturalist 172:160–169.

Ortego J, Knowles LL. 2022. Geographical isolation versus dispersal: Relictual alpine grasshoppers support a model of interglacial diversification with limited hybridization. Molecular Ecology 31:296–312.

Pan D, Hülber K, Willner W, Schneeweiss GM. 2020. An explicit test of Pleistocene survival in peripheral versus nunatak refugia in two high mountain plant species. Molecular Ecolology 29:172–183.

Patterson N, Moorjani P, Luo Y, Mallick S, Rohland N, Zhan Y, Genschoreck T, Webster T, Reich D. 2012. Ancient admixture in human history. Genetics 192:1065–1093.

Pfeifer B, Wittelsbürger U, Ramos-Onsins SE, Lercher MJ. 2014. PopGenome: an efficient Swiss army knife for population genomic analyses in R. Molecular Biology and Evolution 31:1929–1936.

Phillips FM, Zreda M, Plummer MA, Elmore D, Clark DH. 2009. Glacial geology and chronology of Bishop Creek and vicinity, eastern Sierra Nevada, California. Geological Society of America Bulletin 121:1013–1033.

Pickrell JK, Pritchard JK. 2012. Inference of population splits and mixtures from genome-wide allele frequency data. PLOS Genetics 8:e1002967.

Poplin R, Ruano-Rubio V, DePristo MA, Fennell TJ, Carneiro MO, Van der Auwera GA, Kling DE, Gauthier LD, Levy-Moonshine A, Roazen D. 2017. Scaling accurate genetic variant discovery to tens of thousands of samples. BioRxiv:201178.

Porinchu DF, MacDonald GM, Bloom AM, Moser KA. 2003. Late Pleistocene and early Holocene climate and limnological changes in the Sierra Nevada, California, USA inferred from midges (Insecta: Diptera: Chironomidae). Palaeogeography, Palaeoclimatology, Palaeoecology 198:403–422.

Pritchard JK, Di Rienzo A. 2010. Adaptation–not by sweeps alone. Nature Reviews Genetics 11:665–667.

Quinlan AR, Hall IM. 2010. BEDTools: a flexible suite of utilities for comparing genomic features. Bioinformatics 26:841–842.

Rees JS, Castellano S, Andrés AM. 2020. The genomics of human local adaptation. Trends in Genetics 36:415–428.

Rellstab C, Zoller S, Sailer C, Tedder A, Gugerli F, Shimizu KK, Holderegger R, Widmer A, Fischer MC. 2020. Genomic signatures of convergent adaptation to Alpine environments in three Brassicaceae species. Molecular Ecology 29:4350–4365.

Rood DH, Burbank DW, Finkel RC. 2011. Chronology of glaciations in the Sierra Nevada, California, from 10Be surface exposure dating. Quaternary Science Reviews 30:646–661.

Rovito SM, Schoville SD. 2017. Testing models of refugial isolation, colonization and population connectivity in two species of montane salamanders. Heredity (Edinb) 119:265–274.

Schat J, Weng Y-M, Dudko RY, Kavanaugh DH, Luo L, Schoville SD. 2022. Evidence for niche conservatism in alpine beetles under a climate-driven species pump model. Journal of Biogeography n/a.

Schoville SD, Roderick GK. 2009. Alpine biogeography of Parnassian butterflies during Quaternary climate cycles in North America. Molecular Ecology 18:3471–3485.

Schoville SD, Roderick GK, Kavanaugh DH. 2012. Testing the ‘Pleistocene species pump’in alpine habitats: lineage diversification of flightless ground beetles (Coleoptera: Carabidae: Nebr*ia) in* relation to altitudinal zonation. Biological Journal of the Linnean Society 107:95–111.

Segarra-Moragues JG, Palop-Esteban M, González-Candelas F, Catalán P. 2007. Nunatak survival vs. tabula rasa in the Central Pyrenees: a study on the endemic plant species Borderea pyrenaica (Dioscoreaceae). Journal of Biogeography 34:1893–1906.

Sim Z, Hall JC, Jex B, Hegel TM, Coltman DW. 2016. Genome-wide set of SNPs reveals evidence for two glacial refugia and admixture from postglacial recolonization in an alpine ungulate. Mol Ecol 25:3696–3705.

Sinclair BJ. 2000a. Water relations of the freeze-tolerant New Zealand alpine cockroach *Celatoblatta quinquemaculata* (Dictyoptera: Blattidae). Journal of Insect Physiologya 46:869–876.

Sinclair BJ. 2000b. Water relations of the freeze-tolerant New Zealand alpine cockroach Celatoblatta quinquemaculata (Dictyoptera: Blattidae). Journal of Insect Physiology 46:869–876.

Slatyer RA, Schoville SD. 2016. Physiological limits along an elevational gradient in a radiation of montane ground beetles. PloS One 11:e0151959.

Storey K. 1990. Biochemical adaptation for cold hardiness in insects. Philosophical Transactions of the Royal Society of London. B, Biological Sciences 326:635–654.

Storey KB, Storey JM. 2012. Insect cold hardiness: metabolic, gene, and protein adaptation. Canadian Journal of Zoology 90:456–475.

Szpiech ZA, Novak TE, Bailey NP, Stevison LS. 2021. Application of a novel haplotype-based scan for local adaptation to study high-altitude adaptation in rhesus macaques. Evolution Letters 5:408–421.

Sømme L. 1989. Adaptations of terrestrial arthropods to the alpine environment. Biological Reviews 64:367–407.

Teets NM, Peyton JT, Ragland GJ, Colinet H, Renault D, Hahn DA, Denlinger DL. 2012. Combined transcriptomic and metabolomic approach uncovers molecular mechanisms of cold tolerance in a temperate flesh fly. Physiological Genomics 44:764–777.

Thorat L, Nath BB. 2018. Insects with survival kits for desiccation tolerance under xxtreme water deficits. Frontiers in Physiology 9:1843.

Uchida T, Furukawa M, Kikawada T, Yamazaki K, Gohara K. 2019. Trehalose uptake and dehydration effects on the cryoprotection of CHO–K1 cells expressing TRET1. Cryobiology 90:30–40.

Van der Auwera GA, Carneiro MO, Hartl C, Poplin R, Del Angel G, Levy-Moonshine A, Jordan T, Shakir K, Roazen D, Thibault J. 2013. From FastQ data to high-confidence variant calls: the genome analysis toolkit best practices pipeline. Current protocols in bioinformatics 43:11.10. 11-11.10. 33.

Weng YM, Kavanaugh DH, Schoville SD. 2021a. Drainage basins serve as multiple glacial refugia for alpine habitats in the Sierra Nevada Mountains, California. Molecular Ecology 30:826–843.

Weng YM, Francoeur CB, Currie CR, Kavanaugh DH, Schoville SD. 2021b. A high-quality carabid genome assembly provides insights into beetle genome evolution and cold adaptation. Molecular Ecology Resources 21:2145–2165.

Weng YM, Veire BM, Dudko RY, Medeiros MJ, Kavanaugh DH, Schoville SD. 2020. Rapid speciation and ecological divergence into North American alpine habitats: the Nippononebria (Coleoptera: Carabidae) species complex. Biological Journal of the Linnean Society 130:18–33.

Weng YM, Yang MM, Yeh WB. 2016. A comparative phylogeographic study reveals discordant evolutionary histories of alpine ground beetles (Coleoptera, Carabidae). Ecology and Evolution 6:2061–2073.

Wolfe GR, Hendrix DL, Salvucci ME. 1998. A thermoprotective role for sorbitol in the silverleaf whitefly, Bemisia argentifolii. Journal of Insect Physiology 44:597–603.

Yamamoto K, Yamaguchi M, Endo S. 2020. Functional characterization of an aldose reductase (bmALD1) obtained from the silkworm Bombyx mori. Insect Molecular Biology 29:490–497.

Yu G, Wang L-G, Han Y, He Q-Y. 2012. clusterProfiler: an R package for comparing biological themes among gene clusters. Omics: a journal of integrative biology 16:284–287.

Zhang Y, Hu Y, Wang X, Jiang Q, Zhao H, Wang J, Ju Z, Yang L, Gao Y, Wei X et al. 2019. Population Structure, and Selection Signatures Underlying High-Altitude Adaptation Inferred From Genome-Wide Copy Number Variations in Chinese Indigenous Cattle. Frontiers in Genetics 10:1404.

Zhang ZY, Chen B, Zhao DJ, Kang L. 2013. Functional modulation of mitochondrial cytochrome c oxidase underlies adaptation to high-altitude hypoxia in a Tibetan migratory locust. Proceedings of the Royal Society B: Biological Sciences 280:20122758.

Zhao D, Zhang Z, Cease A, Harrison J, Kang L. 2013. Efficient utilization of aerobic metabolism helps Tibetan locusts conquer hypoxia. BMC Genomics 14:631.

Zhao D, Zheng C, Shi F, Xu Y, Zong S, Tao J. 2021. Expression analysis of genes related to cold tolerance in *Dendroctonus valens*. PeerJ 9:e10864.

Zheng X, Levine D, Shen J, Gogarten SM, Laurie C, Weir BS. 2012. A high-performance computing toolset for relatedness and principal component analysis of SNP data. Bioinformatics 28:3326–3328.

