## supplementary_file_1 for "Evidence for admixture and rapid evolution during glacial climate change in an alpine specialist"

**Additional File 1**

**Selection and drift of an alpine specialist following rapid deglaciation**

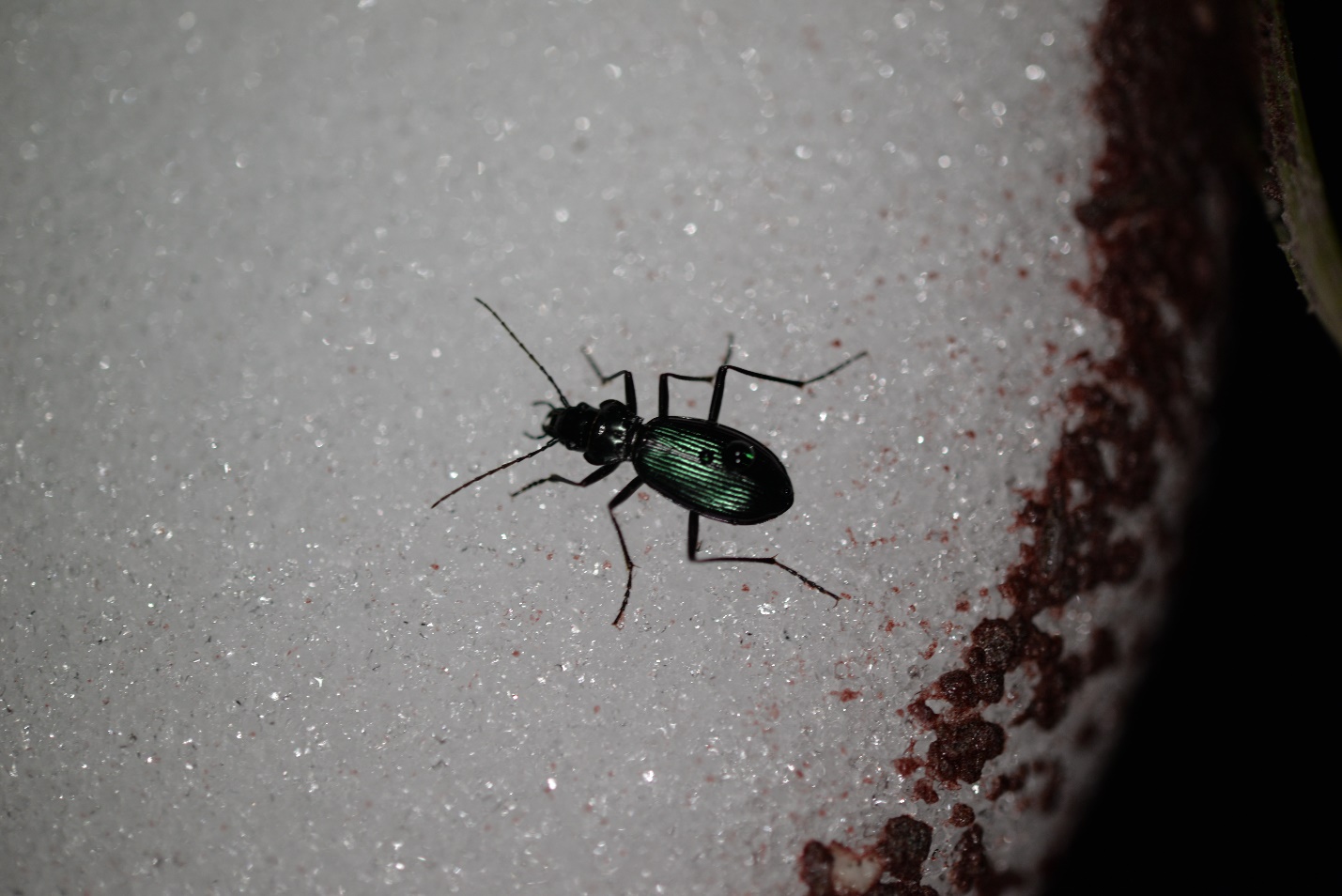

**Figure S1.** *Nebria ingens* complex (*N. riversi*) is foraging on the snowpack during summer night in the nival zone of Sierra Nevada in the Yosemite National Park. Photo credit: Yi-Ming Weng, August 07, 2017

### **Supplementary Results**

#### *Accuracy of four different variant calling methods*

With the low-coverage whole genome sequence reads of 382 beetle samples from 27 sample sites (**Table S1**), we were able to identify large number of single nucleotide polymorphisms (SNPs) by mapping the reads to the reference genome (Weng, Francoeur, et al. 2021). To filter the low-quality SNPs accurately and efficiently, we accessed four different SNPs calling methods to optimize the SNP callset (**Figure S2**). Among the four callsets, the original bam (without BQSR) + VQSR kept the most SNPs among the four filtering approaches and had only a marginal increase in mismatch rate (2.94%) compared to the lowest observed rate (2.74% under hard filtering) (**Table S2 and Figure S3**). We therefore selected this method for downstream analyses, which comprises 5.5 million SNPs. However, missing genotype data remained high across individuals (17.53%), so we used *beagle* version 4.1 to impute missing genotypes after dropping eight samples with > 90% missing genotype data (**Figure S4**) (Browning and Browning 2016). After imputation, the mismatch rate in genotype calls increased to 3.5%. We applied a final filtering step to remove alleles with minor allele frequency less than 5%, resulting in a dataset of 1,238,058 SNPs for downstream analyses.

**Table S1.** Sample localities, sample size, and number of sequences used in the analyses.

| Site name | Site code | no. beetle sequenced | no. sequences used in analyses | latitude | longitude |
| --- | --- | --- | --- | --- | --- |
| Conness Lake | Conness | 57 | 53 | 37.97251 | -119.31404 |
| Kuna Lake | Kuna | 1 | 1 | 37.84874 | -119.25929 |
| Donohue Pass | Donohue | 16 | 16 | 37.75289 | -119.24847 |
| Lyell Peak | Lyell | 15 | 15 | 37.74956 | -119.25991 |
| Ritter Range | Ritter | 8 | 8 | 37.71442 | -119.20355 |
| Ottoway Lake | Ottoway | 3 | 3 | 37.63967 | -119.39713 |
| Recess Lakes | FRecess | 9 | 9 | 37.43124 | -118.78462 |
| Ruby Lake | Ruby | 29 | 28 | 37.41151 | -118.77009 |
| Italy Lake | Italy | 10 | 10 | 37.34084 | -118.76735 |
| Selden Pass | Selden | 13 | 13 | 37.30436 | -118.91287 |
| Piute Pass | Piute | 22 | 22 | 37.23683 | -118.679944 |
| Lamarck Lakes | Lamarck | 16 | 16 | 37.20691 | -118.65303 |
| Hungry Packer Lake | HungryPacker | 10 | 10 | 37.16094 | -118.64077 |
| Treasure Lake | Treasure | 20 | 17 | 37.14056 | -118.57619 |
| Sam Mack Lake | SamMack | 16 | 16 | 37.11646 | -118.51212 |
| Dusy Pass | Dusy | 5 | 5 | 37.09534 | -118.53421 |
| Taboose Pass | Taboose | 13 | 13 | 36.97993 | -118.39877 |
| Sixty Lakes | Sixty | 4 | 4 | 36.82776 | -118.43623 |
| Sphinx Lakes | Sphinx | 2 | 2 | 36.71427 | -118.51487 |
| North Forester | NForester | 4 | 4 | 36.71056 | -118.3711 |
| Milly's Footpass | Millys | 13 | 13 | 36.69039 | -118.43453 |
| South Forester | SForester | 16 | 16 | 36.68882 | -118.37648 |
| Wright Lakes | Wright | 14 | 14 | 36.63116 | -118.34489 |
| Pear Lake | Pear | 12 | 12 | 36.59707 | -118.6658 |
| Crabtree Lakes | Crabtree | 13 | 13 | 36.54445 | -118.32363 |
| Army Pass | Army | 35 | 35 | 36.50159 | -118.24711 |
| Monarch Lake | Monarch | 6 | 6 | 36.41429 | -118.54783 |

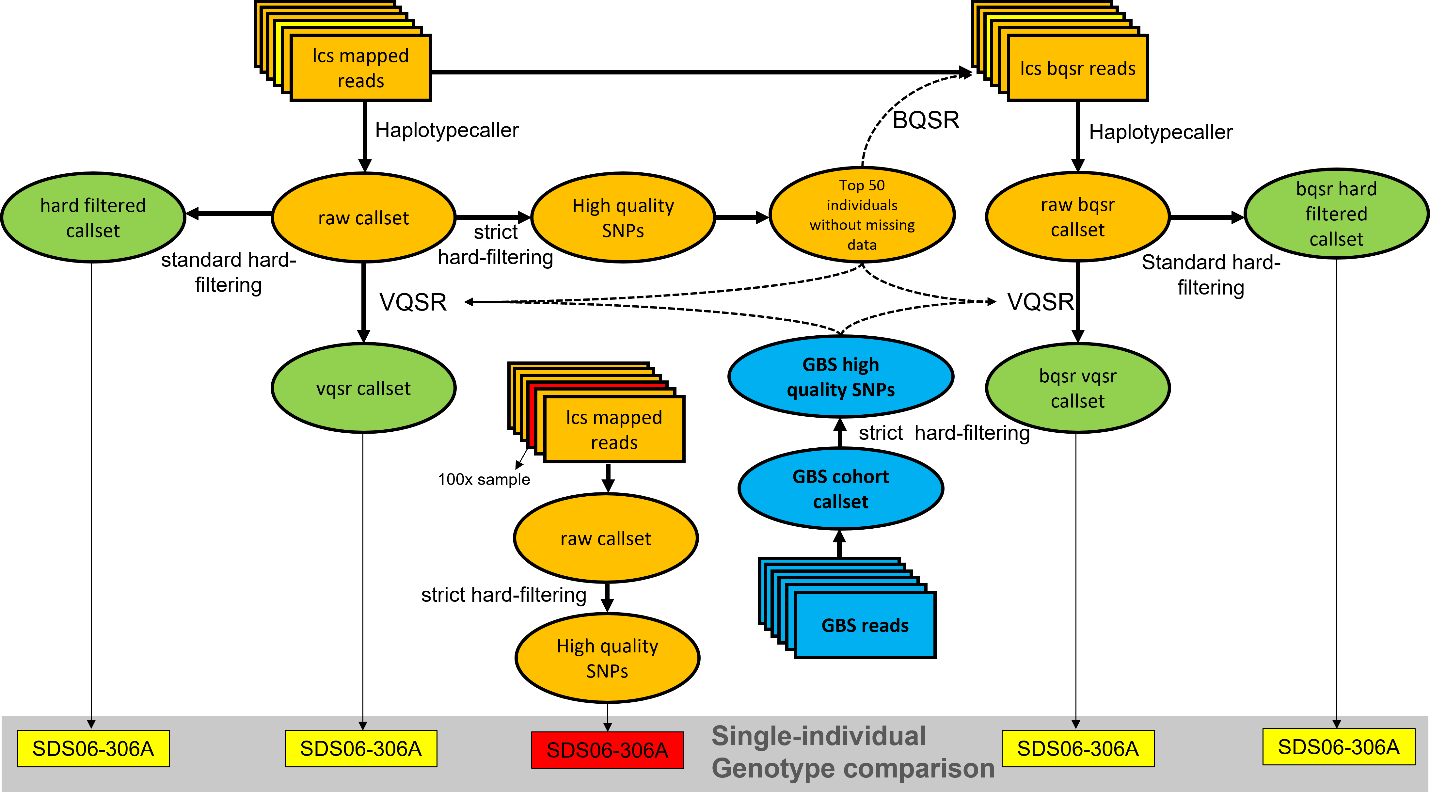

**Figure S2.** The workflow of variant calling for this study. A calibration step (BQSR) and two SNP filtering steps (VQSR and hard filtering) were considered in assessing variant calls. We examined a single individual (SDS06-306A) sequenced at both a low (5x) and a high (171.4x) depth of coverage. The comparison (mismatch rate) between the low and high coverage genotypes provided a performance measure for the accuracy of variant calling. Abbreviation: lcs=low coverage sequence; GBS=genotype by sequence; VQSR= Variant Quality Score Recalibration; BQSR= Base Quality Score Recalibration.

**Table S2.** The accuracy of the final SNP call sets with different calibration and filtering methods

|  | vqsr trainning set (from GBS) | bqsr/vqsr trainning set | raw callset | strict hard filtered | standard hard filtered | vqsr at 99.9 | bqsr plus standard hard filtered | bqsr plus vqsr at 99.9 |
| --- | --- | --- | --- | --- | --- | --- | --- | --- |
| number of snp loci | 21602 | 169953 | 5880298 | 2422050 | 4766328 | 5543251 | 3848354 | 4581821 |
| number of non-missing snp loci | - | - | 4831308 | 2000883 | 3937844 | 4571606 | 3177297 | 3777213 |
| missing rate (%) | - | - | 17.84 | 17.39 | 17.38 | 17.53 | 17.44 | 17.56 |
| matched genotype loci | - | - | 1934534 | 1921325 | 1934453 | 1924944 | 1625792 | 1619303 |
| mismatch genotype loci | - | - | 61212 | 52586 | 61131 | 56602 | 63829 | 59830 |
| mismatch rate (%) | - | - | 3.16 | 2.74 | 3.16 | 2.94 | 3.93 | 3.67 |

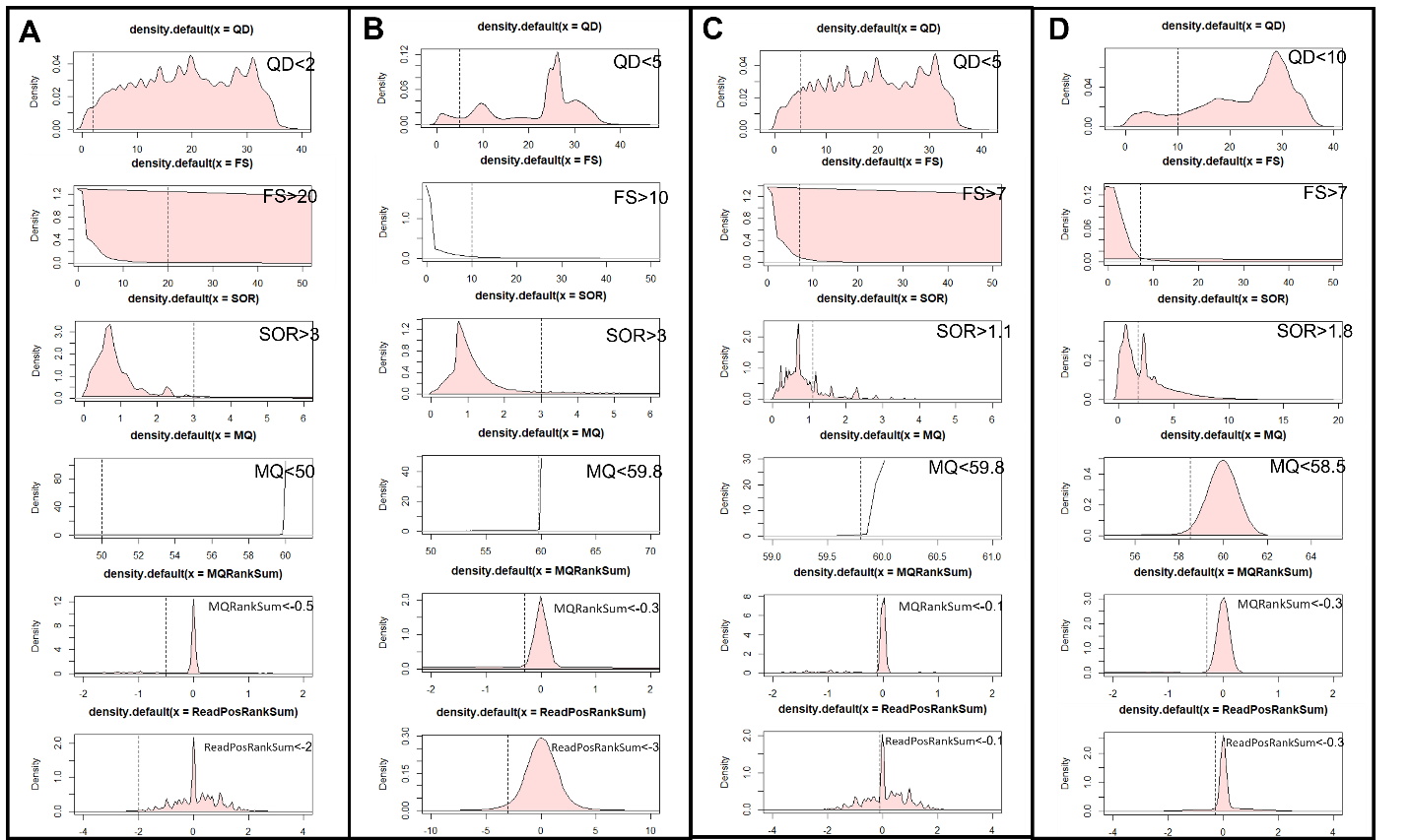

**Figure S3.** The distribution and cutoff for various variant calling metrics. Datasets were generated by (A) filtering the original callset, (B) strict filtering the high coverage single-sample callset, (C) use of the strict filtered original call set for a training reference in BQSR and VQSR, and (D) use of the strict filtered call set from GBS reads for a training reference in VQSR.

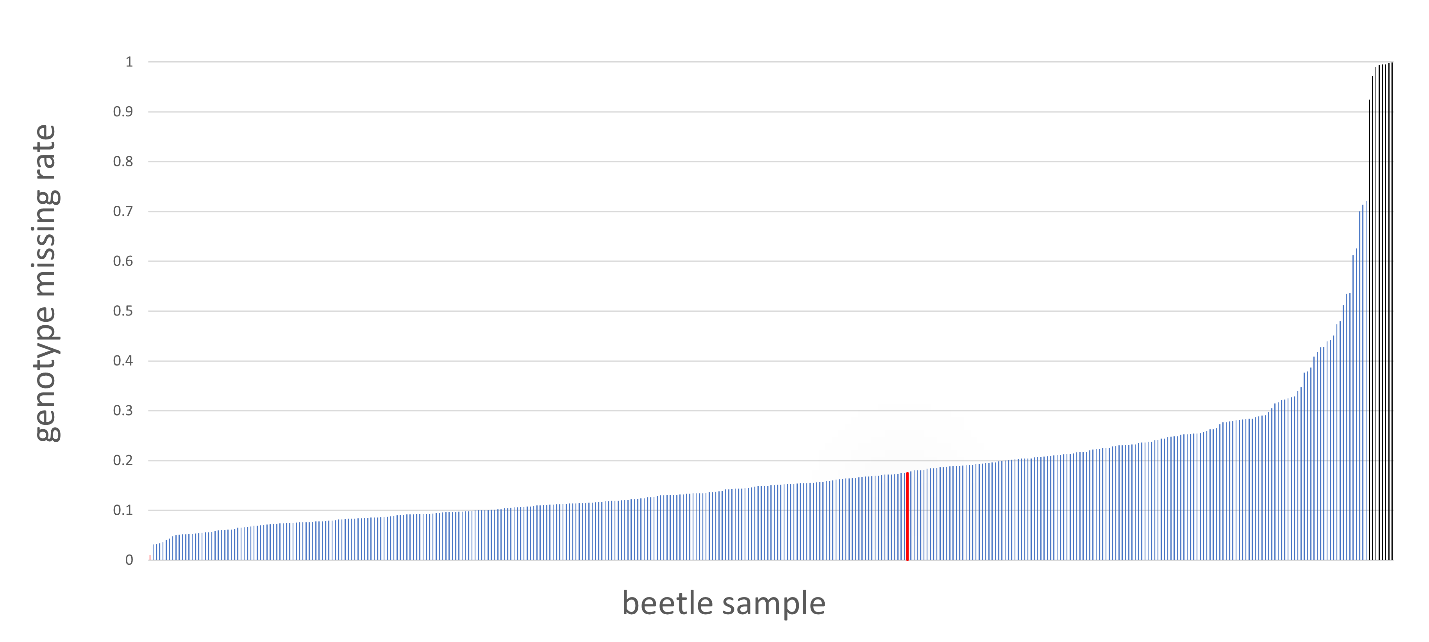

**Figure S4.** The rate of missing genotype data across beetle samples. The black bars denote the 8 samples with more than 90% of missing genotype data, which were dropped prior to imputation. The red bar denotes the sample (SDS06-306A) low coverage dataset, which was used to assess the genotyping error rate relative to a high coverage dataset for the same individual.

#### *Identifying autosomal contigs and their LD decay plot*

To accurately estimate the diploid genomic *F*_ST_ and *d_XY_* of paired populations, we exclude the SNPs locate on putative sex linkage chromosome. The sex linkage contig with mean male-female read depth ratio are expected to be 0.5 since the XY and XO sex-determination systems are primarily present in many other carabid species. Based on the read depth ratio distribution (**Figure S5**), we apply a cutoff for the ratio of 0.65 for sex-linked chromosome. As a result, 110 contigs (8,874,574 bps, ca. 6% of the genome) were identified as putative sex linkage chromosome and 2,027 contigs as autosomal (138,474,936 bps, ca. 94% of the genome) (**supplementary file 2: contig read depth ratio**). Among the 110 putative sex linkage contigs, 80% (88 contigs) were found in a previously published list of putative sex linkage chromosome contigs (Weng et al., 2021). The mean LD decay plots from the autosomal contigs shows a significant decline in the *R*^2^ when the SNPs are more than 3 kbp away from each other, and the decay line reaches an equilibrium at *R*^2^=0.02 starting from the distance of the paired SNPs being 5 kbp (**Figure S6**).

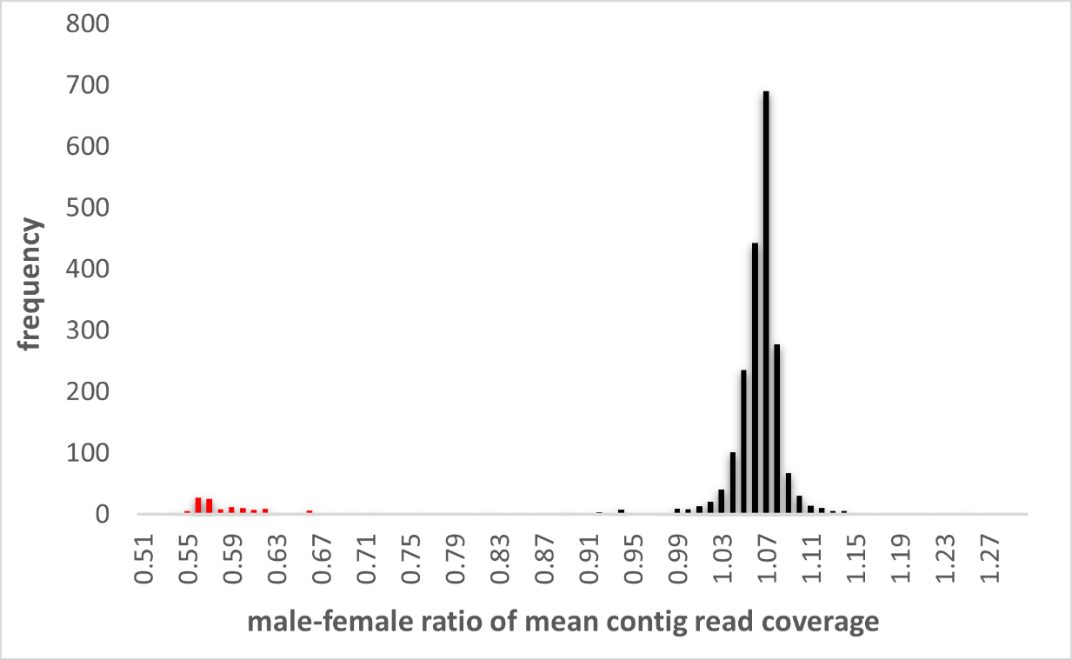

**Figure S5.** The frequency plot of the male-female ratio of mean contig read coverage. The autosomal contigs are expected to have a 1.0 male-female ratio (black bars). Of 2,027 autosomal contigs, 1,345 contigs are larger than 10kbp. The sex linkage chromosome contigs are expected to have 0.5 male-female ratio (red bars).

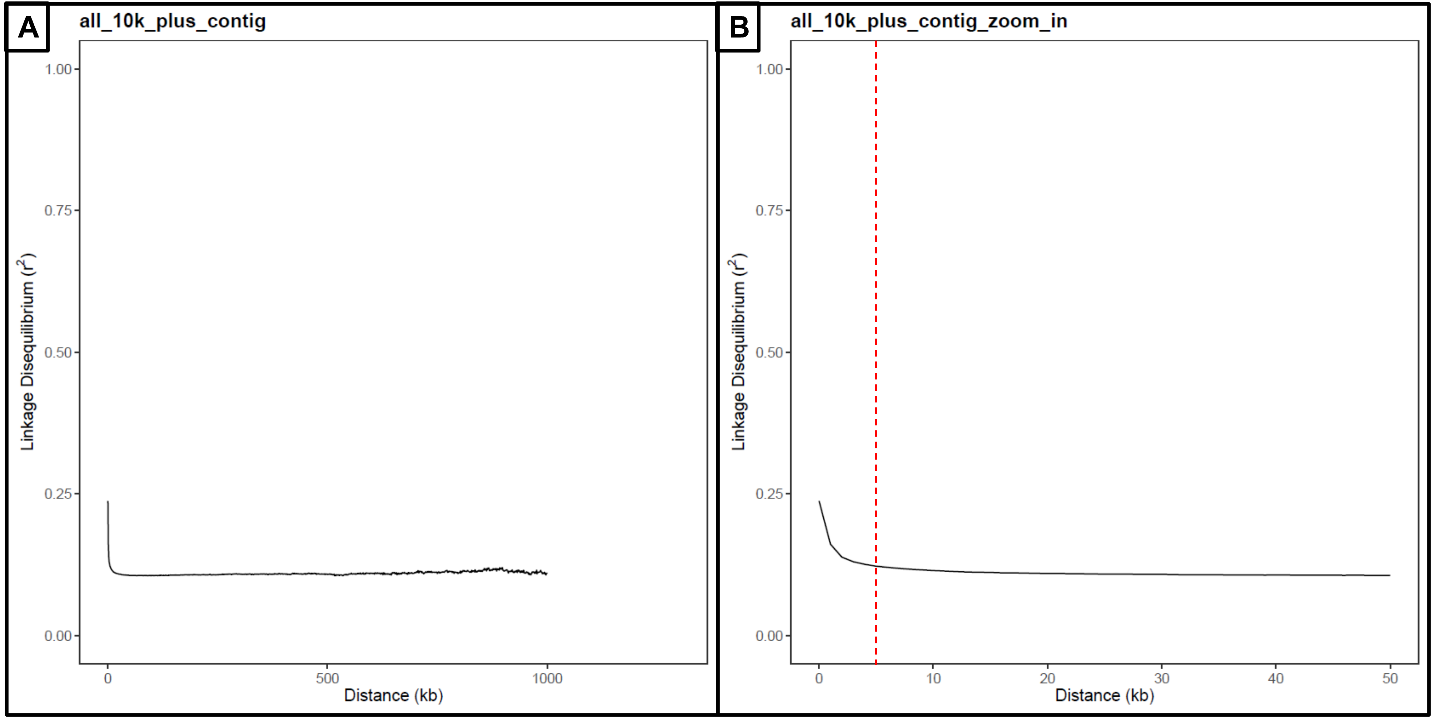

**Figure S6.** The decay of linkage disequilibrium. Mean values are shown for (A) autosomal contigs in the range of 1 Mb, as well as (B) zoomed into the first 50 Kb. The red vertical dashed line in (B) shows the threshold at 5 Kb that was used to prune SNPs for PCA and sNMF analyses.

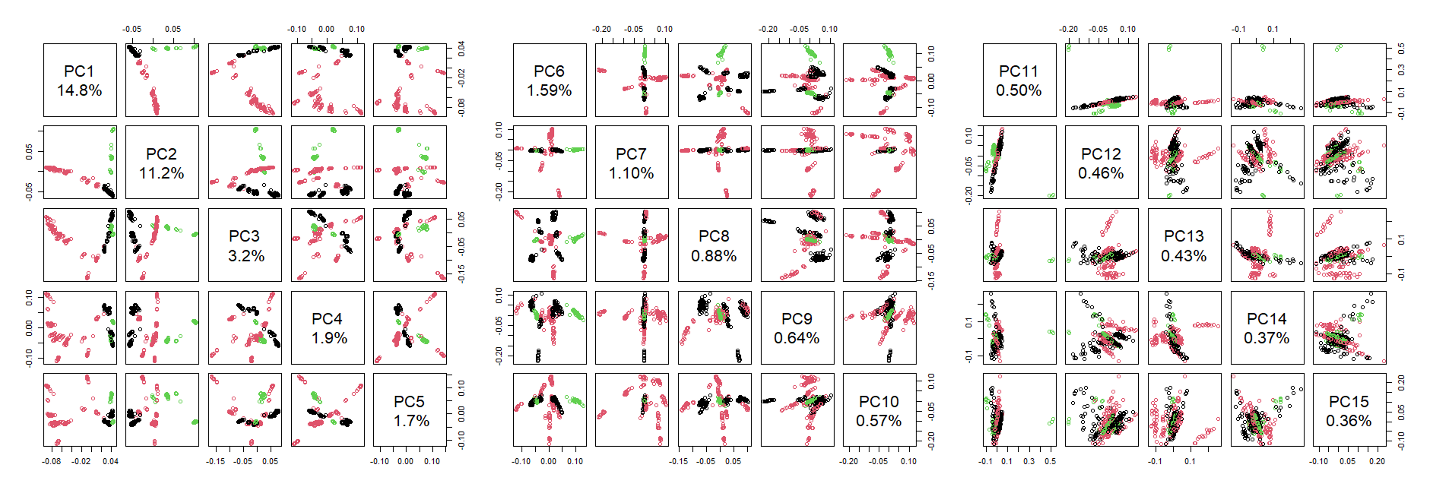

**Figure S7**. The genetic structure of *N. ingens* complex using PCA based on the maf+LD filtered dataset with 378,532 SNPs. The three lineages (*N. ingens*, imtermediate, and *N. riversi*) are colored in red, black, and green, respectively. We selected the first 15 PCs to show the genetic structure in the three panels.

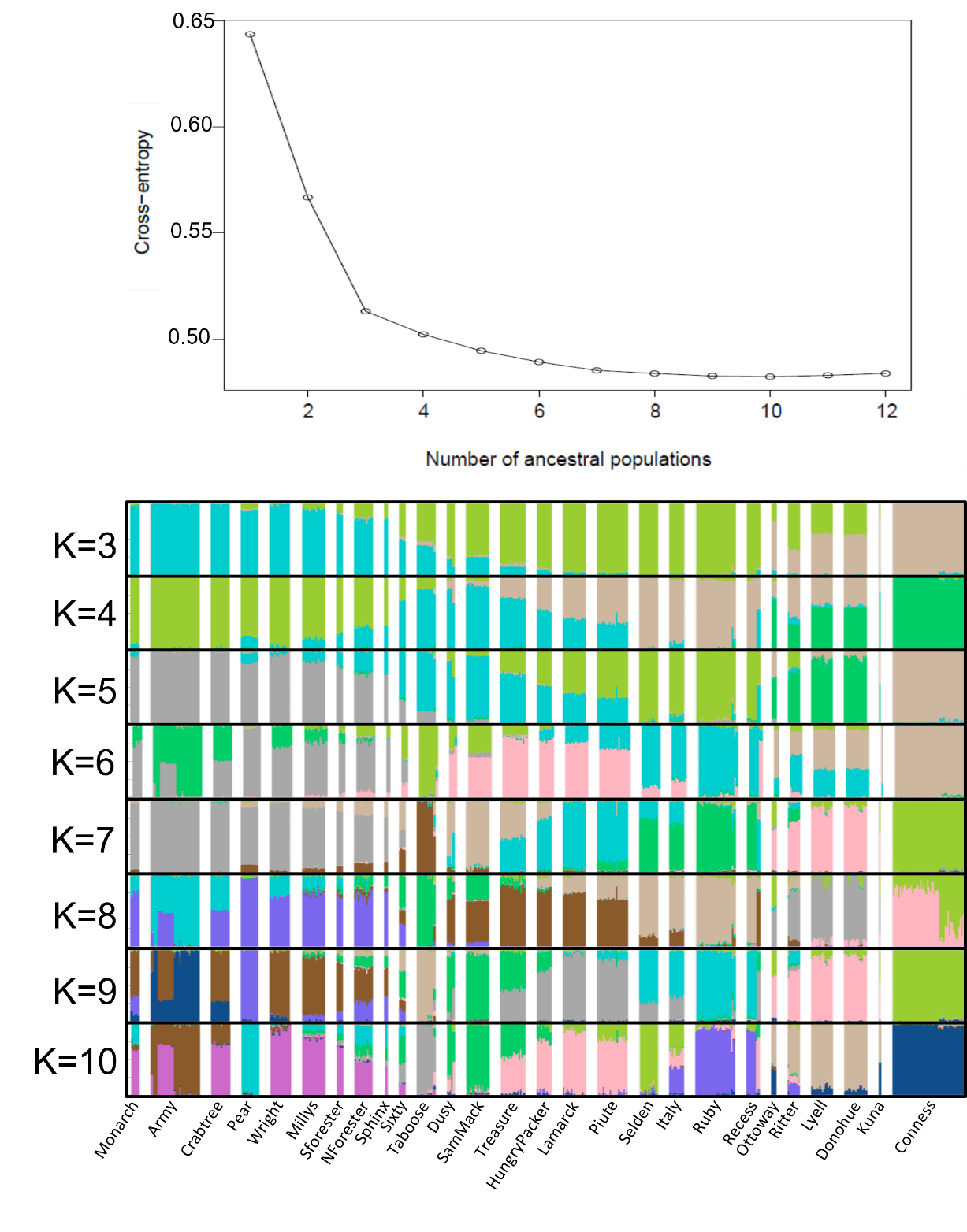

**Figure S8**. The genetic structure of *N. ingens* complex using sNMF based on the maf+LD filtered dataset. The top panel shows the cross-entropy from K=1 to K=12 and bottom panel shows the clusters from K=3 to K=10. The three lineages include *N. ingens* (from Monarch to Taboose), intermediate (Dusy to Recess), and *N. riversi* (Ottoway to Conness). The delimitation of the three lineages can be seen when K=3 but the gradual genetic changes present in many populations.

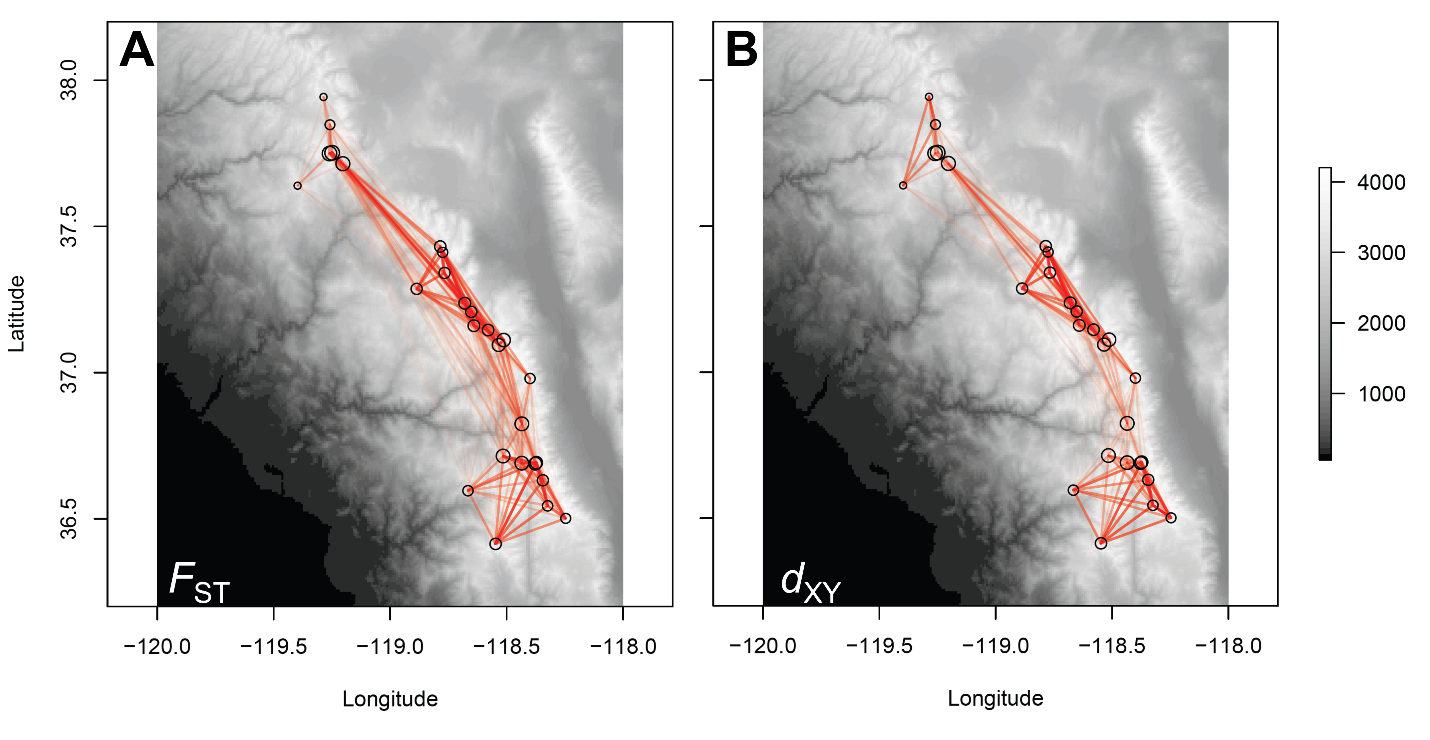

**Figure S9**. The genetic structure of *N. ingens* complex shown by pairwise *F*_ST_ (A) and *d_XY_* (B). The red line between population denotes the level of genetic divergence, with thinner and fainter line shows higher genetic divergence (i.e. higher *F*_ST_ or *d_XY_*). The circles represent the populations where larger circle means higher genetic diversity of the population. Note that the *d_XY_* is absolute measurement of genetic distance thus more independent to the effect from the genetic diversity of the populations. This is especially notable in the northern population pairs where the inflation of *F*_ST_ can be seen (thinner lines) for those populations with less genetic diversity (smaller circles), which is not seen in *d_XY_*.

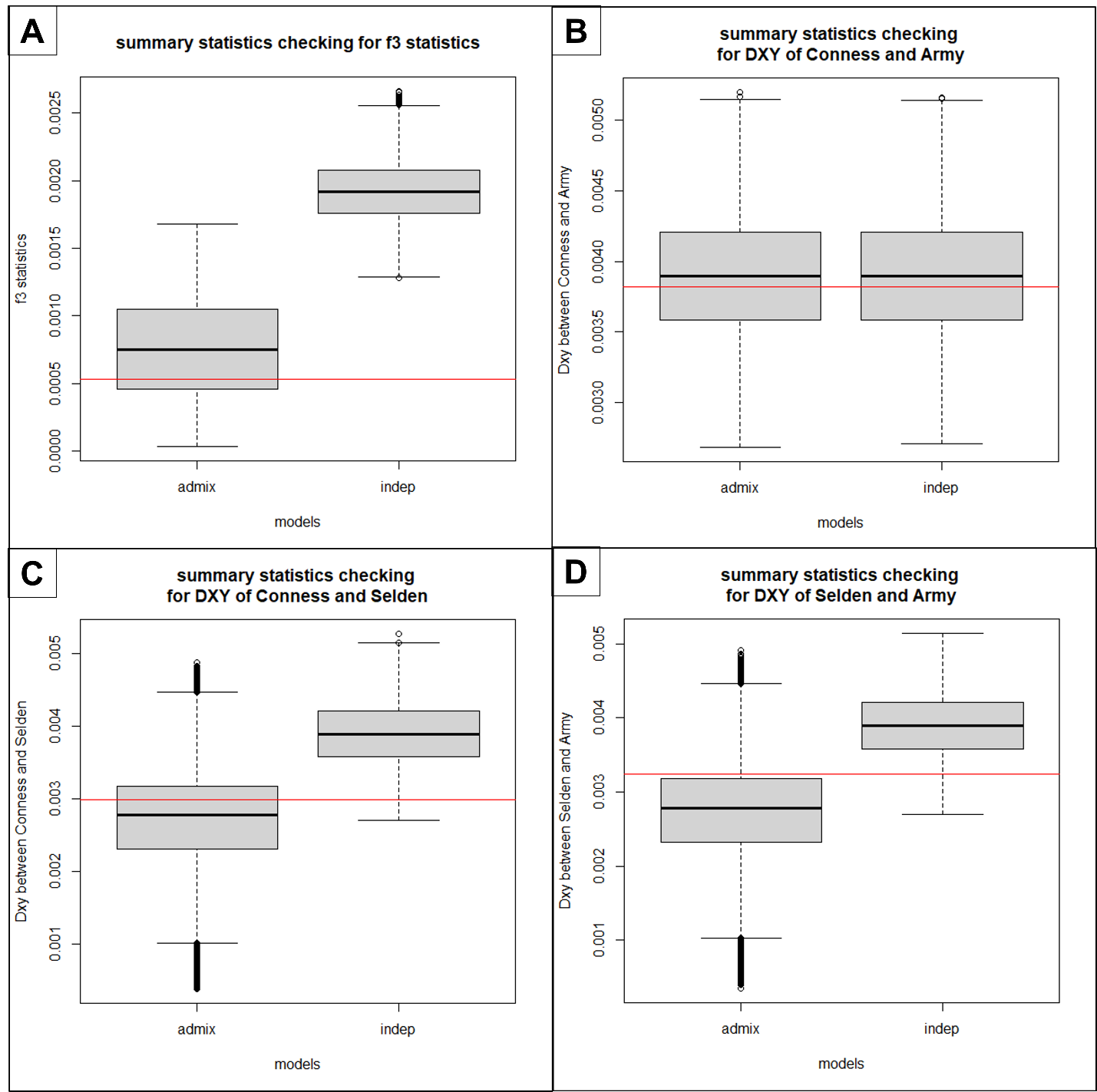

**Figure S10.** The distributions of the summary statistics from the data of two simulated models (boxes) and the observed genotypes (red line). The range of *f*3-statistics (A) from the admixture model (left) covers the observed *f*3-statistics but not for the independent model (right), implying better fit of admixture model. The *d_XY_* statistics from the three populations (B-D) have less distinctive results to support each model, as the observed *d_XY_* statistics fall into the ranges of both models.

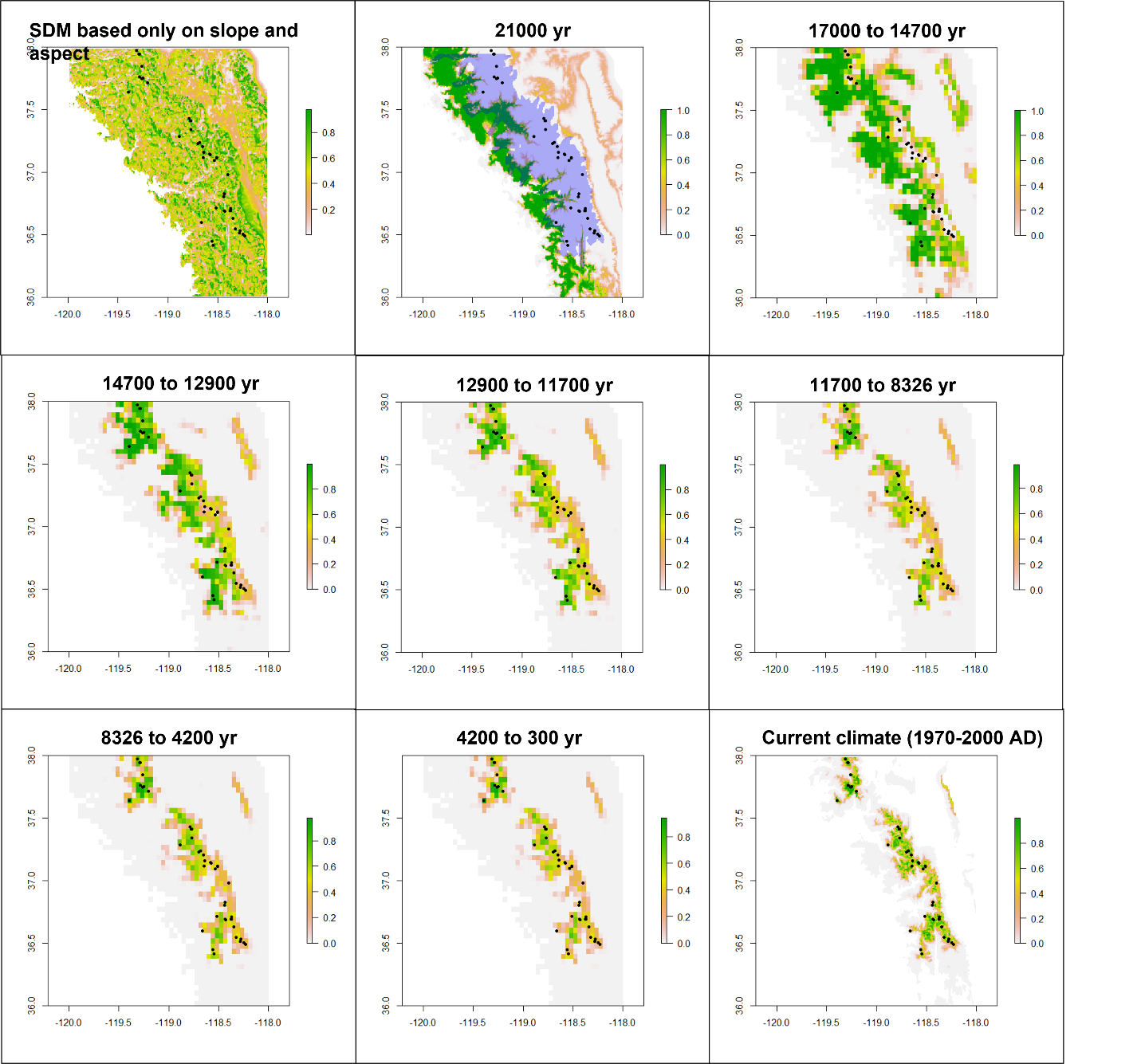

**Figure S11.** The species distribution models of different time periods. The black dots represent the sites of occurrence, and the color shows the possibility of occurrence from zero (white) to 1 (green). For the SDM of last glacial maximum period (21,000 yr), we applied the glacier boundary (the blue polygon) to the distribution map, and for the patches in the range of the glacier boundary, the suitability (possibility of occurrence) is set to be zero.

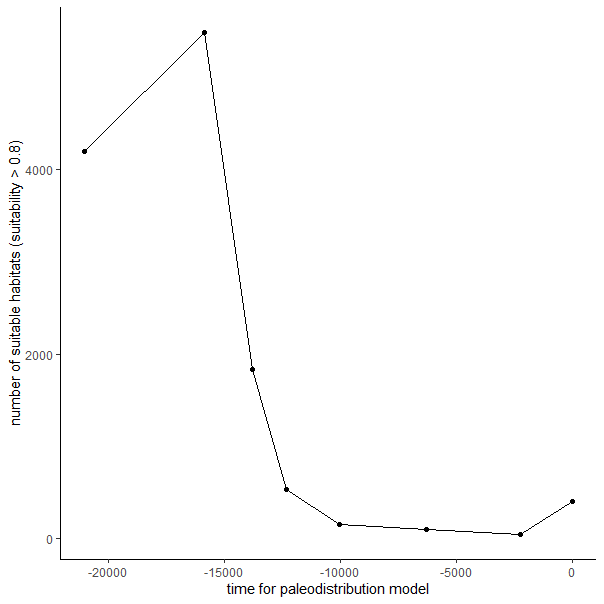

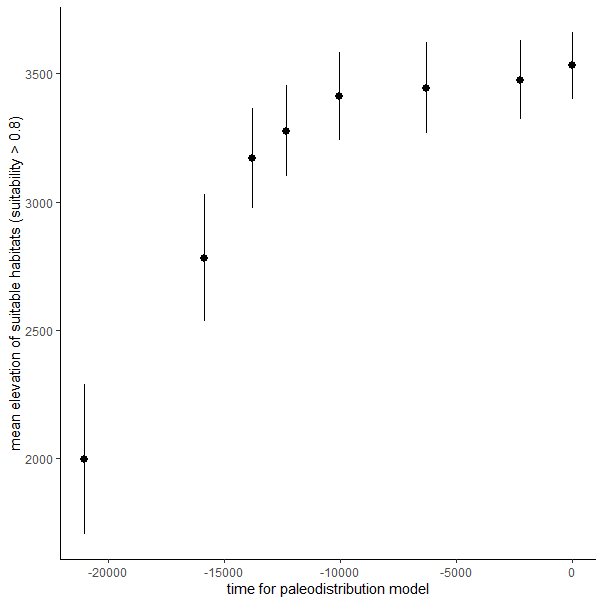

**Figure S12.** The change of suitable habitat throughout the glacial retreat from the LGM (21,000 yr) to present day, estimated from the paleodistribution models in **Figure S12**. (left) Suitability reached the lowest point around 10,000 years ago. (right) Mean elevation of suitable habitat increased over time, plateauing around 10,000 years ago. The error bars in the elevation plots shows the standard deviation of the elevation from all the suitable habitats (suitability > 0.8).

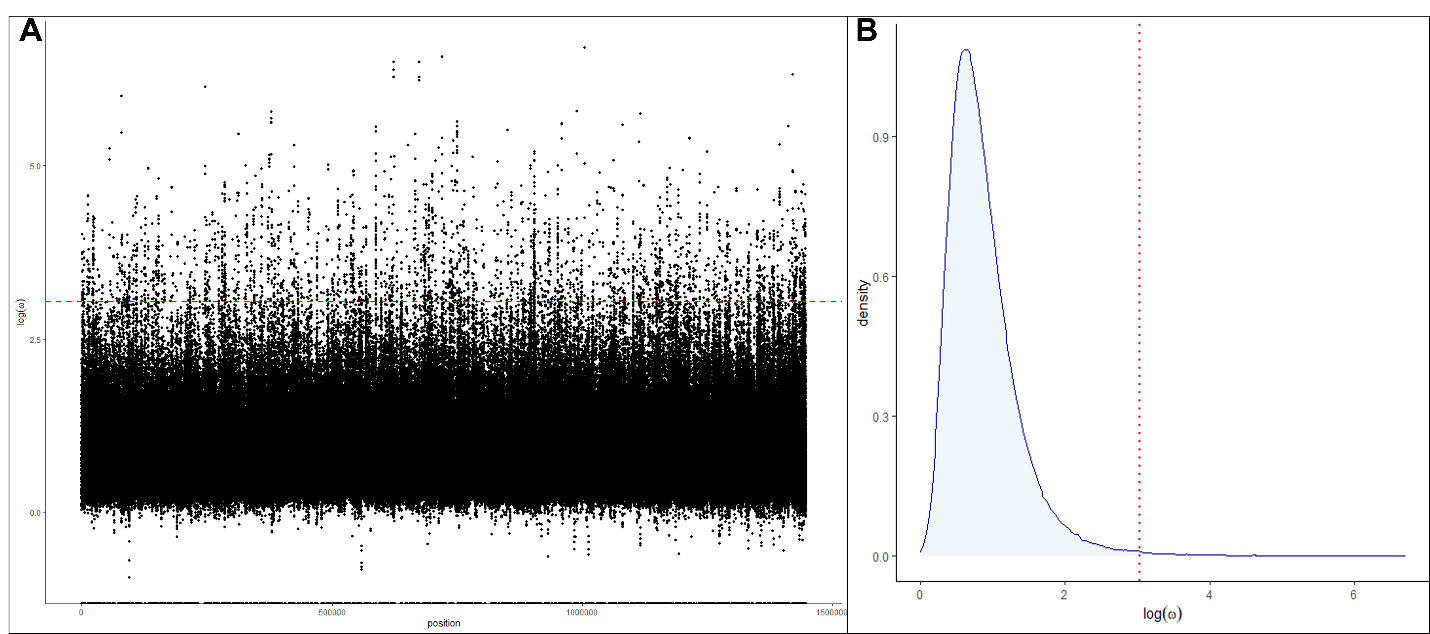

**Figure S13.** The Manhattan plot of *ω*-statistic (A) and their density distribution (B). Both plots show the *ω*-statistic with log transformation, and the red lines denote the threshold of 99.5% quantile that was used to define the significantly exceeding *ω*-statistics.

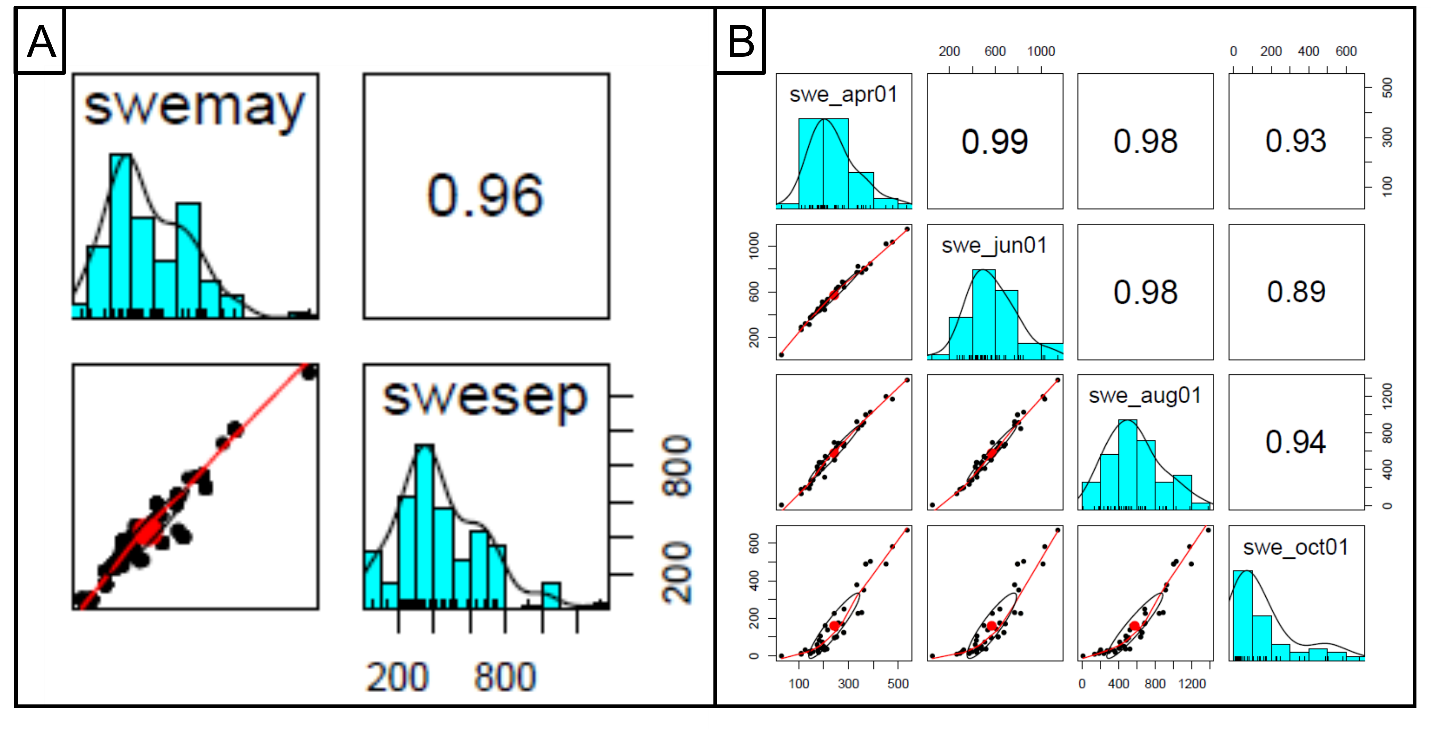

**Figure S14.** The high temporal correlations of SWE of monthly data (May and September, A) and daily data (B) during non-snowing season. The data were the averages of 31-year data extracted from the 45 unique occurrence sites of *N. ingens* complex.

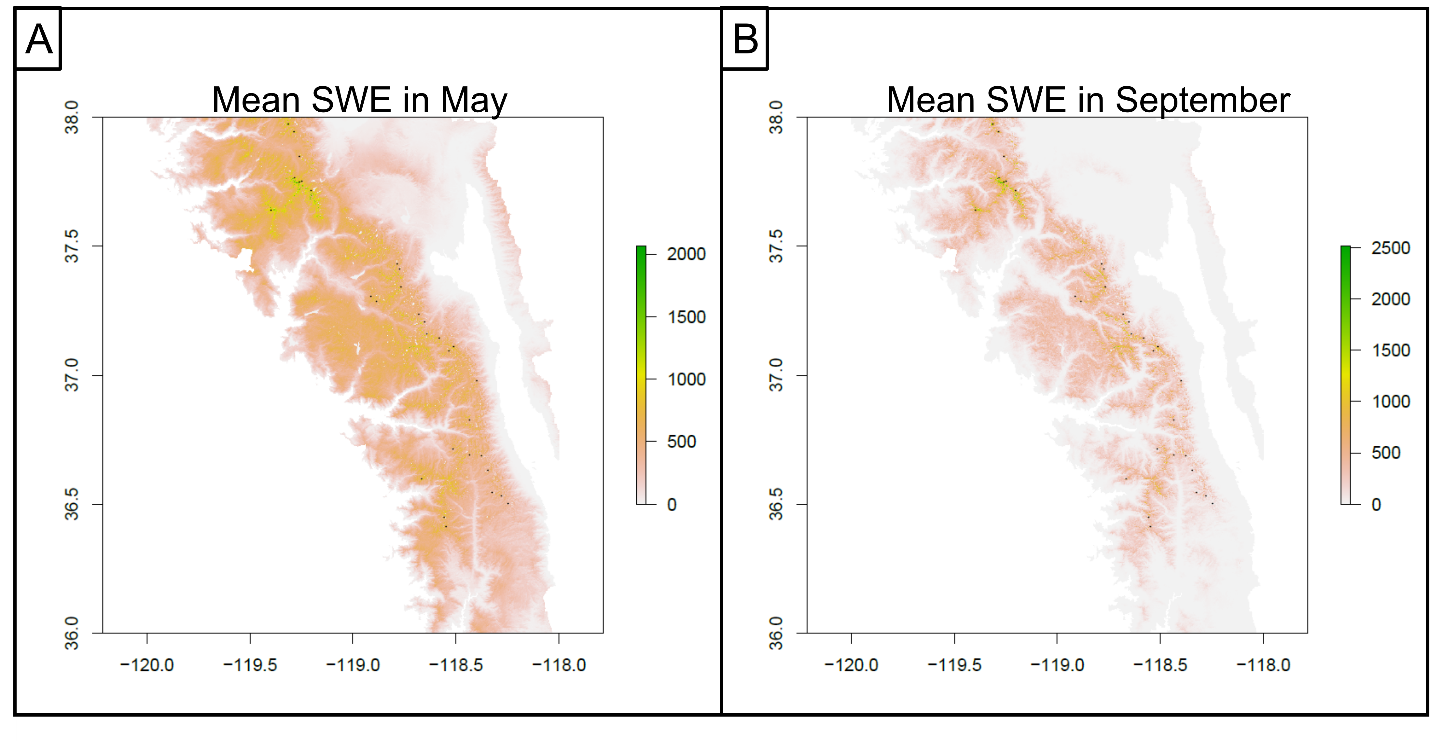

**Figure S15.** The maps show the mean SWE in the Sierra Nevada Mountain in May (A) and September (B). The black dots denote the 45 unique occurrence sites of *N. ingens* complex.

**Supplementary Methods**

#### *Variant Calling pipeline*

For most non-model organisms, accurately calling genetic variants is challenging due to the lack of a reference panel from the well-established databases. Additionally, sequence data is often low coverage, which can bias downstream population genetic analyses if it leads to a high incidence of genotyping errors (Fountain, et al. 2016; Lou, et al. 2021). To ensure that genotyping error was controlled in our study, we compared the mismatch of genotypes from low and high coverage sequence data from an individual beetle and applied different filtering methods to optimize the variant calling pipeline.

All samples were first assessed for read quality using *fastqc* (Andrews 2010). Low quality reads, barcode and adaptor sequences, the first 10 base pairs at the 5’ end with unusual GC ratio, and low-quality bases were removed from both ends of the reads using the *bbduk* tool in *bbmap* v. 38.86 (Bushnell 2014). We also used *bbduk* to remove potential contamination sequences by referencing the UniVec database from NCBI ([www.ncbi.nlm.nih.govVecScreenUniVec.html](http://www.ncbi.nlm.nih.govVecScreenUniVec.html)). The resulting clean reads were then mapped to the reference genome of *Nebria riversi* (contig level assembly, GenBank assembly accession: GCA_018344505.1) to generate the binary alignment and map (bam) files using *bwa* version 0.7.17 (Li, et al. 2009; Li 2013; Weng, Francoeur, et al. 2021). The bam files with mapped reads were sorted and marked with duplicates using *samtools* version 1.7 and *Picard* version 2.24.1 (Li, et al. 2009; Picard 2018). Variant calling steps included: 1) *HaplotypeCaller* to identify variants (both SNPs and indels) for each individual and generate a single-sample genomic variant call format (GVCF) file; 2) *GenomicsDBImport* to imports the GVCFs into a database; and 3) *GenotypeGVCFs* to combine the GVCF files into a cohort VCF file.

With the combination of with/without base quality score recalibration (BQSR) and two different variant filtering processes in the GATK variant calling pipeline (hard-filtering and variant quality score recalibration, VQSR), four cohort variant callsets were used to access the genotyping accuracy by comparing the genotype of an individual beetle (SDS06-306A) from these four cohort callsets to the same individual genotype with high coverage (**Figure S2**). Specifically, the base quality score recalibration is performed using BQSR function in the *GATK* pipeline (McKenna, et al. 2010; DePristo, et al. 2011), which generates new bam files. For both original (without BQSR) and BQSR bam files, we applied either standard hard-filtering (**Figure S3B**) or variant quality score recalibration (VQSR) at tranche 99.9 to remove the low-quality SNPs. The base quality recalibration and SNP filtering steps resulted in four datasets with different filtering approaches (original bam + hard-filtering, BQSR + hard-filtering, original bam + VQSR, and BQSR + VQSR). The final callset with highest accuracy (least mismatch in genotype comparison) was used in imputed for the downstream analyses.

To perform BQSR, a reliable reference panel with known genotypes is required. However, a reliable reference panel for the *N. ingens* complex does not exit. We thus used a complete SNP callset (loci without missing data) from the top 50 individuals with the highest coverage generated from original bam + strict hard filtering to serve as a reference panel for BQSR (**Figure S3C**). For VQSR, we used two reference panels to control the quality of SNP calls. The first is the one used for BQSR, and the second is a SNP callset from a genotype-by-sequence (GBS) sequence data downloaded from NCBI (BioProject number PRJNA645878) (Weng, Kavanaugh, et al. 2021). The GBS sequence data consists of 311 individuals partially overlapping with the 383 individuals in this study, with the mean read depth of 15.2x. Similarly, we used stricter hard filtering to remove low-quality SNPs from this GBS callset (**Figure S3D**).

The accuracy of genotypes from these 4 callsets was accessed by comparing the genotype of an individual (SDS06-306A, read depth=5.6X) in these four callsets to the high-quality genotype of same individual beetle with high coverage (read depth=171.4X). The generation of the high-quality genotype of this individual was following the GATK pipeline without BQSR and with the use of stricter hard-filtering process (**Figure S3B**). Specifically, we downloaded the high coverage sequence reads from NCBI (accession number SRX8725733; Weng et al., 2021), mapped the reads to the reference genome using bwa mem (Li 2013), and followed the standard GATK variant calling pipeline with stricter hard-filtering step to keep high-quality SNPs (**Figure S3B**). The genotype comparison was done using the customized script.

#### *Identifying autosomal contigs and their LD decay plot*

To determine parameters for pruning variants according to statistical non-independence, we examined several measures of linkage (e.g. the mean length of linkage blocks and variant correlation coefficient within linkage blocks). Adjusting for linkage improves the performance population genetic analyses that depend on statistical independence, such as PCA and sNMF. We first identified autosomal contigs as those expected to have 1:1 ratio of contig read depth when comparing males (XO) to females (XX), while the contigs from the X chromosome should show a 1:2 ratio comparing males to females. We first dropped six samples of unknown sex and seven with very low coverage, resulting in a dataset of 253 females and 116 males. The read depth of each contig for each individual was then calculated using samtools (Li, et al. 2009). We then averaged the coverage for females and males, separately, and measured the male-female ratio of the mean coverage. The contigs with read depth ratio less than 0.65 were defined as sex-linked (**Figure S5**). With the list of putative autosomal contigs, the LD decays were evaluated on these contigs by calculating pairwise correlations between SNPs within a range of 1 Mb using *plink* v1.90b6.21 (Zheng, et al. 2012). The individual contig LD decay plots as well as the average LD decay plot were subsequently generated using R.

#### *ABC model simulation*

For both evolutionary divergence models (see **Figure 1** of the main text), we set a uniform prior for the time of the ancestral split of all three lineages (*τ*_1_) to a range from 487,868 to 687,868 generations ago, assuming two years per generation. This prior was set according to the estimate of median divergence time of *N. ingens* and *N. riversi* using the mitochondrial COI gene under growth model by Schoville *et al*. (2012). The range of the divergence time was set with median ± 100,000 generations as the 95% confidence interval is large (51,626-3,162,497 generations ago) from the estimation using mitochondrial COI sequence data. The prior for the admixture event leading to the intermediate lineage (*τ*_2_) was set to a uniform distribution ranging from 57,500 to 480,000 generations ago, which is between the lower boundary of *τ*_1_ and the timing of the last glacial maximum. Based on field observations and estimates from Weng *et al*. (2021), each lineage was set to have an effective population size ranging from 1,000 to 10,000 individuals, with the ancestral effective population size set to 10x the size of descendant lineages. Population sizes were held constant to reduce the number of free parameters in each model. In the simulations, we introduced mutation rate of 2.8 × 10^-9^ per site per generation from the estimate of *Drosophila* and recombination rate of 2.48 × 10^-8^ (2.48 ± 1.31 cM/Mb) from many coleopteran species. Each model was run 1 million times through *msprime* and the summary statistics including the admixture index, outgroup-*f*3 statistic, and the genetic distance *d*_XY_ are calculated with *tskit* (Nei and Li 1979; Patterson, et al. 2012; Kelleher, et al. 2016).

#### *Distribution models of the Nebria ingens complex during different time periods*

To simulate the spatial and demographic change over time from LGM (21,000 yr) to present day, we reconstructed a series of distribution models based on both geographic and climatic variables (**Figure S11**). The geographic variables include elevation, terrain characteristics (i.e. slope and aspect), and the distance to the nearest drainage (available water resource). The slope and aspect are calculated from the elevation raster using the function “terrain” in the R package “raster” (Hijmans and van Etten 2012), where the elevation data (the 30 second digital elevation model) was downloaded from WorldClim database version 2.1 (Fick and Hijmans 2017). The distance to the nearest drainage was calculated using the function “gridDistance” in the R package “raster” with the drainage system map downloaded from Data Basin (NHD Stream and Rivers, Sierra Nevada Zones; data uploaded and owned by Conservation Biology Institute). For the climatic variables, we downloaded contemporary climate data (1970-2000 AD) at 30-second resolution from WorldClim 2.1 and paleoclimatic data at 2.5-minute resolution (except for the LGM data which is 30-second resolution and originally from CHELSA database) from PaleoClim (Brown, et al. 2018; Karger and Zimmermann 2019).

For the current species distribution model (SDM), we used 4 geographic variables including elevation, slope, aspect, and the distance to the nearest drainage, with 3 climatic variables including the mean snow water equilibrium in May (same as we used in GEA analysis, **Figure S14-S15**), annual mean temperature, and annual precipitation to predict the distribution probability of the species complex. These variables were selected based on their biological relevance (e.g. requiring cold, icy or riparian habitats) to optimize the accuracy of SDM. To find the best niche model, the model selection was performed using the function ENMevaluate in the R package ENMeval v2.0.3 with the use of maxent.jar v3.4.3 from dismo package v1.3.5 (Kass, et al. 2021). The background records were randomly sampled from the region above 1,000 meters elevation around the Sierra Nevada Mountain, and the testing models were defined with different combinations of maxent feature classes ("L", "LQ", "H", "LQH", "LQHP", "LQHPT"), with the regularization multiplier (smoothing parameter) being from to 1 to 30 with the increment of 2. The occurrence data is described in the published work by Weng, Kavanaugh, & Schoville, 2021. Best model was then chosen based on minimum corrected Akaike Information Criterion (AICc). This model, however, can not be used to predict the historical distributions because the snow water equilibrium is unavailable for paleoclimatic data, and the elevation should not be considered as determining variable as it has been assumed that the altitudinal range of distribution could change with the change of climate. Accordingly, a subset of variables (slope, aspect, distance to the nearest drainage, annual mean temperature, and annual precipitation) was considered in searching the best niche model based on minimum AICc for predicting the historical SDMs (**Figure S11**). For the SDM of last glacial maximum period (21,000 yr), we applied the glacier boundary of LGM to the species distribution map by zeroing the possibility of occurrence (suitability) for the patches within the glacier boundary (Rood, et al. 2011), as the environment within the range is extremely harsh and unlikely to provide large habitats for the species (**Figure S11**). However, the same data is not available for other 7 time periods, thus this glacier boundary was applied only on the SDM of last glacial maximum period. Finally, to provide the baseline distribution model in the initial step of the simulation, we created the distribution model with the use of only terrain characteristics (**Figure S11**).

#### *Calculating mean snow water equilibrium for GEA analyses from the database*

For the snow water equilibrium (SWE) data used in SDM and genotype-environment association (GEA) analyses, we first downloaded the SWE data from the research website of Margulis Research Group (<https://margulis-group.github.io/data/>). The detail for SWE modeling is described in the webpage and the related publications (Margulis, et al. 2016). The data includes daily SWE data for 31 years (1985–2015) for the Sierra Nevada Mountain area and we focused only on the mean SWE in May over the 31 years for both biological and computational reasons. For biological reason, we use SWE in May because the last snow usually occurs before May and the adults of *N. ingens* complex emerges starting from late May to early June. For the computational reason, we found that the SWE data shows high temporal correlation in the period between last snow (April) and first snow of the year (usually in October) (**Figure S14**). Thus, we used the mean SWE in May as proxy of the mean SWE of non-snowing season in the purpose of reducing the computational demand. For the data extraction from the database, we used H5Dump to convert the daily SWE data of Mays in each year from hdf format to ascii format. We then calculated the average of SWE over the 930 days (30 days in May times 31 years in the database) and crop the range to be with 36-38 degree in latitude and -120 to -118 in longitude with the R package “raster” (Hijmans and van Etten 2012) (**Figure S15**).

**Table S3.** Enriched biological process terms from the 18 elevation associated genes

| Term ID | Name | log(*p*-value) |
| --- | --- | --- |
| GO:0051656 | establishment of organelle localization | -3.5087 |
| GO:0043303 | mast cell degranulation | -2.2254 |
| GO:0002279 | mast cell activation involved in immune response | -2.2254 |
| GO:0045576 | mast cell activation | -2.2254 |
| GO:0060376 | positive regulation of mast cell differentiation | -2.9234 |
| GO:0002699 | positive regulation of immune effector process | -2.2241 |
| GO:0033005 | positive regulation of mast cell activation | -2.0798 |
| GO:0033008 | positive regulation of mast cell activation involved in immune response | -2.0798 |
| GO:0002763 | positive regulation of myeloid leukocyte differentiation | -2.022 |
| GO:0060375 | regulation of mast cell differentiation | -2.9234 |
| GO:0019372 | lipoxygenase pathway | -2.6227 |
| GO:0033275 | actin-myosin filament sliding | -2.4468 |
| GO:0030049 | muscle filament sliding | -2.4468 |
| GO:0010927 | cellular component assembly involved in morphogenesis | -2.3335 |
| GO:0036462 | TRAIL-activated apoptotic signaling pathway | -2.4468 |
| GO:1903353 | regulation of nucleus organization | -2.4468 |
| GO:1901568 | fatty acid derivative metabolic process | -2.2327 |
| GO:0035995 | detection of muscle stretch | -2.022 |
| GO:0034374 | low-density lipoprotein particle remodeling | -2.1465 |
| GO:0031394 | positive regulation of prostaglandin biosynthetic process | -2.9234 |
| GO:2001280 | positive regulation of unsaturated fatty acid biosynthetic process | -2.9234 |
| GO:0031392 | regulation of prostaglandin biosynthetic process | -2.9234 |
| GO:2001279 | regulation of unsaturated fatty acid biosynthetic process | -2.9234 |
| GO:2001235 | positive regulation of apoptotic signaling pathway | -2.2072 |
| GO:0061886 | negative regulation of mini excitatory postsynaptic potential | -2.4468 |
| GO:0099149 | regulation of postsynaptic neurotransmitter receptor internalization | -2.2254 |
| GO:1903422 | negative regulation of synaptic vesicle recycling | -2.1465 |
| GO:1902804 | negative regulation of synaptic vesicle transport | -2.022 |
| GO:0051640 | organelle localization | -3.0874 |
| GO:2001053 | regulation of mesenchymal cell apoptotic process | -2.0798 |
| GO:0010743 | regulation of macrophage derived foam cell differentiation | -2.022 |
| GO:0048769 | sarcomerogenesis | -2.3221 |
| GO:0030240 | skeletal muscle thin filament assembly | -2.0798 |
| GO:2001054 | negative regulation of mesenchymal cell apoptotic process | -2.0798 |
| GO:0042759 | long-chain fatty acid biosynthetic process | -2.3221 |
| GO:0010744 | positive regulation of macrophage derived foam cell differentiation | -2.9234 |
| GO:1903530 | regulation of secretion by cell | -2.1043 |
| GO:0060627 | regulation of vesicle-mediated transport | -2.0052 |
| GO:1903593 | regulation of histamine secretion by mast cell | -2.3221 |
| GO:1903595 | positive regulation of histamine secretion by mast cell | -2.3221 |
| GO:0043302 | positive regulation of leukocyte degranulation | -2.022 |
| GO:0043306 | positive regulation of mast cell degranulation | -2.0798 |
| GO:0055013 | cardiac muscle cell development | -2.2254 |

**Table S4.** Enriched biological process terms from the nine precipitation associated genes

| Term ID | Name | log(p-value) |
| --- | --- | --- |
| GO:0032528 | microvillus organization | -2.0179 |
| GO:0032868 | response to insulin | -2.65 |
| GO:1901652 | response to peptide | -2.179 |
| GO:0051412 | response to corticosterone | -2.0477 |
| GO:0043434 | response to peptide hormone | -2.3266 |
| GO:0060676 | ureteric bud formation | -2.8914 |
| GO:0061005 | cell differentiation involved in kidney development | -2.0798 |
| GO:0072202 | cell differentiation involved in metanephros development | -2.239 |
| GO:0035850 | epithelial cell differentiation involved in kidney development | -2.29 |
| GO:0072172 | mesonephric tubule formation | -2.29 |
| GO:0072079 | nephron tubule formation | -2.239 |
| GO:1900157 | regulation of bone mineralization involved in bone maturation | -2.8914 |
| GO:0070168 | negative regulation of biomineral tissue development | -2.3479 |
| GO:1900158 | negative regulation of bone mineralization involved in bone maturation | -2.8914 |
| GO:0030502 | negative regulation of bone mineralization | -2.4938 |
| GO:1901143 | insulin catabolic process | -2.29 |
| GO:0090291 | negative regulation of osteoclast proliferation | -2.8914 |
| GO:0070664 | negative regulation of leukocyte proliferation | -2.0477 |
| GO:0090289 | regulation of osteoclast proliferation | -2.7154 |
| GO:0090330 | regulation of platelet aggregation | -2.0477 |
| GO:0009953 | dorsal/ventral pattern formation | -2.3896 |
| GO:0038098 | sequestering of BMP from receptor via BMP binding | -2.29 |
| GO:0060394 | negative regulation of pathway-restricted SMAD protein phosphorylation | -2.0179 |
| GO:0035581 | sequestering of extracellular ligand from receptor | -2.1521 |
| GO:1900116 | extracellular negative regulation of signal transduction | -2.1521 |
| GO:0003257 | positive regulation of transcription from RNA polymerase II promoter involved in myocardial precursor cell differentiation | -2.8914 |
| GO:0003256 | regulation of transcription from RNA polymerase II promoter involved in myocardial precursor cell differentiation | -2.7154 |
| GO:1900115 | extracellular regulation of signal transduction | -2.1521 |
| GO:0060231 | mesenchymal to epithelial transition | -2.4147 |
| GO:0051973 | positive regulation of telomerase activity | -2.1521 |
| GO:1900086 | positive regulation of peptidyl-tyrosine autophosphorylation | -2.5906 |
| GO:0031954 | positive regulation of protein autophosphorylation | -2.1934 |
| GO:1900084 | regulation of peptidyl-tyrosine autophosphorylation | -2.5906 |
| GO:0090189 | regulation of branching involved in ureteric bud morphogenesis | -2.0798 |
| GO:0061217 | regulation of mesonephros development | -2.0798 |
| GO:0090190 | positive regulation of branching involved in ureteric bud morphogenesis | -2.0798 |
| GO:1905209 | positive regulation of cardiocyte differentiation | -2.4147 |
| GO:2000727 | positive regulation of cardiac muscle cell differentiation | -2.4938 |
| GO:0046851 | negative regulation of bone remodeling | -2.7154 |
| GO:0034104 | negative regulation of tissue remodeling | -2.4938 |
| GO:0046850 | regulation of bone remodeling | -2.1521 |
| GO:1900154 | regulation of bone trabecula formation | -2.5906 |
| GO:2000273 | positive regulation of signaling receptor activity | -2.1144 |
| GO:0048263 | determination of dorsal identity | -2.4147 |
| GO:0045647 | negative regulation of erythrocyte differentiation | -2.8914 |
| GO:0002042 | cell migration involved in sprouting angiogenesis | -2.29 |
| GO:0033688 | regulation of osteoblast proliferation | -2.1934 |
| GO:1900155 | negative regulation of bone trabecula formation | -2.5906 |
| GO:1903010 | regulation of bone development | -2.0477 |
| GO:0055025 | positive regulation of cardiac muscle tissue development | -2.1144 |
| GO:1903011 | negative regulation of bone development | -2.29 |
| GO:0033689 | negative regulation of osteoblast proliferation | -2.4147 |
| GO:0003337 | mesenchymal to epithelial transition involved in metanephros morphogenesis | -2.5906 |
| GO:1901228 | positive regulation of transcription from RNA polymerase II promoter involved in heart development | -2.8914 |
| GO:1901213 | regulation of transcription from RNA polymerase II promoter involved in heart development | -2.4938 |
| GO:0061213 | positive regulation of mesonephros development | -2.0798 |

**Table S5.** Enriched biological process terms from the 18 snow water equivalent associated genes

| Term ID | Name | log(p-value) |
| --- | --- | --- |
| GO:0006406 | mRNA export from nucleus | -2.4304 |
| GO:0006405 | RNA export from nucleus | -2.1087 |
| GO:0031110 | regulation of microtubule polymerization or depolymerization | -2.381 |
| GO:0060481 | lobar bronchus epithelium development | -2.8613 |
| GO:0060428 | lung epithelium development | -2.2601 |
| GO:0060052 | neurofilament cytoskeleton organization | -2.1634 |
| GO:0042840 | D-glucuronate catabolic process | -2.2601 |
| GO:0042839 | D-glucuronate metabolic process | -2.2601 |
| GO:0006064 | glucuronate catabolic process | -2.0845 |
| GO:0008592 | regulation of Toll signaling pathway | -2.3691 |
| GO:1900052 | regulation of retinoic acid biosynthetic process | -2.2601 |
| GO:0019747 | regulation of isoprenoid metabolic process | -2.0179 |
| GO:0030656 | regulation of vitamin metabolic process | -2.0845 |
| GO:0007541 | sex determination, primary response to X:A ratio | -2.0845 |
| GO:0043568 | positive regulation of insulin-like growth factor receptor signaling pathway | -2.1634 |
| GO:0016358 | dendrite development | -2.7911 |
| GO:0042501 | serine phosphorylation of STAT protein | -2.2601 |
| GO:0019800 | peptide cross-linking via chondroitin 4-sulfate glycosaminoglycan | -2.0845 |
| GO:0061056 | sclerotome development | -2.1634 |
| GO:0007399 | nervous system development | -2.4041 |

### **Additional References**

Andrews S. 2010. FastQC: a quality control tool for high throughput sequence data. In: Babraham Bioinformatics, Babraham Institute, Cambridge, United Kingdom.

Brown JL, Hill DJ, Dolan AM, Carnaval AC, Haywood AM. 2018. PaleoClim, high spatial resolution paleoclimate surfaces for global land areas. Scientific Data 5:180254.

Browning BL, Browning SR. 2016. Genotype Imputation with Millions of Reference Samples. The American Journal of Human Genetics 98:116-126.

Bushnell B. 2014. BBTools software package. URL <http://sourceforge>. net/projects/bbmap.

DePristo MA, Banks E, Poplin R, Garimella KV, Maguire JR, Hartl C, Philippakis AA, del Angel G, Rivas MA, Hanna M, et al. 2011. A framework for variation discovery and genotyping using next-generation DNA sequencing data. Nat Genet 43:491-498.

Fick SE, Hijmans RJ. 2017. WorldClim 2: new 1‐km spatial resolution climate surfaces for global land areas. International journal of climatology 37:4302-4315.

Fountain ED, Pauli JN, Reid BN, Palsbøll PJ, Peery MZ. 2016. Finding the right coverage: the impact of coverage and sequence quality on single nucleotide polymorphism genotyping error rates. Molecular Ecology Resources 16:966-978.

Hijmans RJ, van Etten J. 2012. raster: Geographic analysis and modeling with raster data. R package version 2.0-12.

Karger DN, Zimmermann NE. 2019. Climatologies at high resolution for the earth land surface areas CHELSA V1. 2: Technical specification. Scientific Data.

Kass JM, Muscarella R, Galante PJ, Bohl CL, Pinilla‐Buitrago GE, Boria RA, Soley‐Guardia M, Anderson RP. 2021. ENMeval 2.0: redesigned for customizable and reproducible modeling of species’ niches and distributions. Methods in Ecology and Evolution.

Kelleher J, Etheridge AM, McVean G. 2016. Efficient Coalescent Simulation and Genealogical Analysis for Large Sample Sizes. PLOS Computational Biology 12:e1004842.

Li H. 2013. Aligning sequence reads, clone sequences and assembly contigs with BWA-MEM. arXiv preprint arXiv:1303.3997.

Li H, Handsaker B, Wysoker A, Fennell T, Ruan J, Homer N, Marth G, Abecasis G, Durbin R. 2009. The sequence alignment/map format and SAMtools. Bioinformatics 25:2078-2079.

Lou RN, Jacobs A, Wilder AP, Therkildsen NO. 2021. A beginner's guide to low-coverage whole genome sequencing for population genomics. Molecular Ecology 30:5966-5993.

Margulis SA, Cortés G, Girotto M, Durand M. 2016. A Landsat-Era Sierra Nevada Snow Reanalysis (1985–2015). Journal of Hydrometeorology 17:1203-1221.

McKenna A, Hanna M, Banks E, Sivachenko A, Cibulskis K, Kernytsky A, Garimella K, Altshuler D, Gabriel S, Daly M. 2010. The Genome Analysis Toolkit: a MapReduce framework for analyzing next-generation DNA sequencing data. Genome research 20:1297-1303.

Nei M, Li WH. 1979. Mathematical model for studying genetic variation in terms of restriction endonucleases. Proceedings of the National Academy of Sciences 76:5269-5273.

Patterson N, Moorjani P, Luo Y, Mallick S, Rohland N, Zhan Y, Genschoreck T, Webster T, Reich D. 2012. Ancient admixture in human history. Genetics 192:1065-1093.

Picard. 2018. Picard toolkit. <http://broadinstitute.github.io/picard/>: Broad Institute, GitHub repository.

Rood DH, Burbank DW, Finkel RC. 2011. Chronology of glaciations in the Sierra Nevada, California, from 10Be surface exposure dating. Quaternary Science Reviews 30:646-661.

Weng YM, Francoeur CB, Currie CR, Kavanaugh DH, Schoville SD. 2021. A high-quality carabid genome assembly provides insights into beetle genome evolution and cold adaptation. Molecular Ecology Resources 21:2145-2165.

Weng YM, Kavanaugh DH, Schoville SD. 2021. Drainage basins serve as multiple glacial refugia for alpine habitats in the Sierra Nevada Mountains, California. Molecular Ecology 30:826-843.

Zheng X, Levine D, Shen J, Gogarten SM, Laurie C, Weir BS. 2012. A high-performance computing toolset for relatedness and principal component analysis of SNP data. Bioinformatics 28:3326-3328.
