## supplementary_file_3 for "Evidence for admixture and rapid evolution during glacial climate change in an alpine specialist"

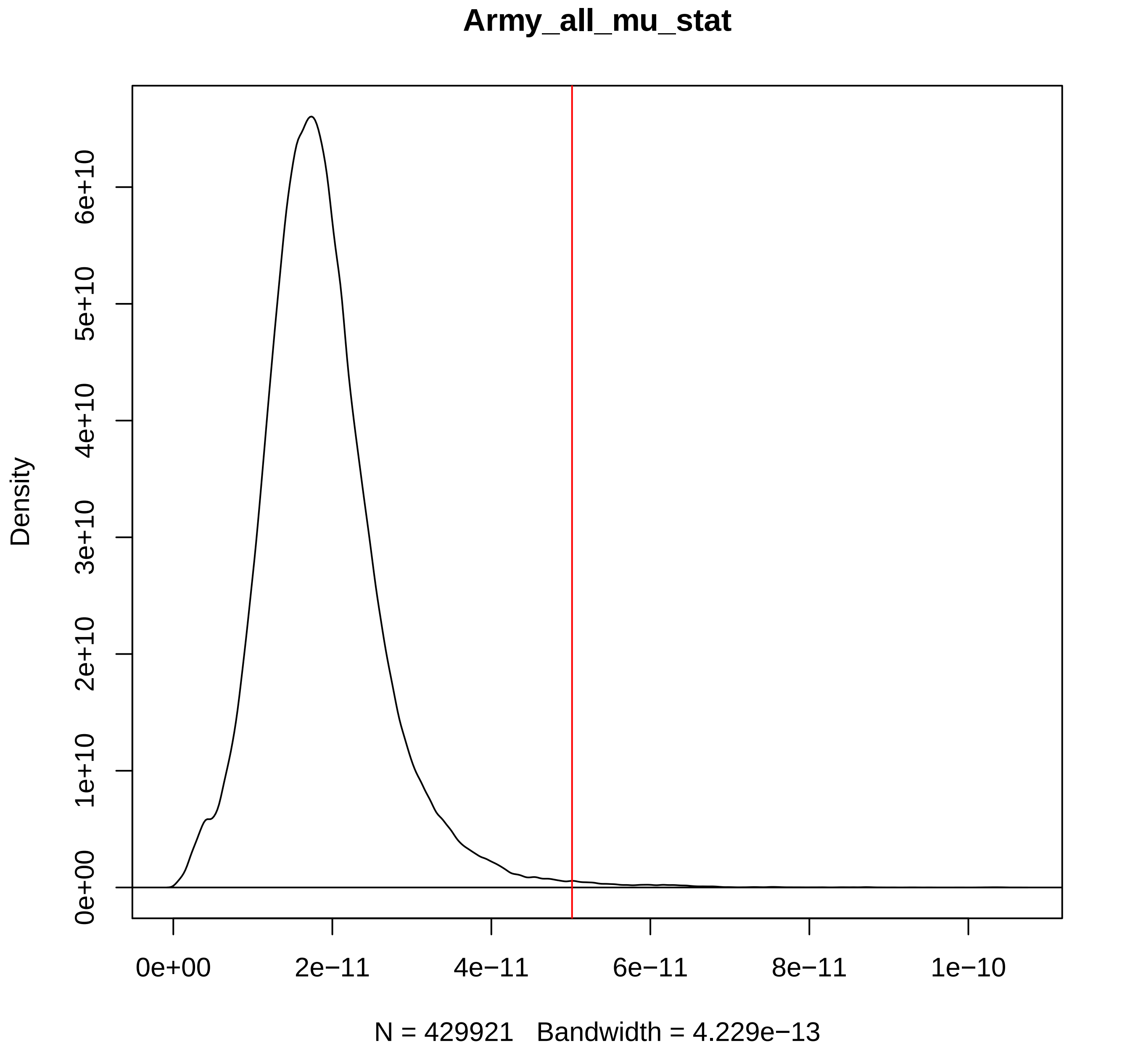

Army_all_mu_stat
Density
0e+00
1e+10
2e+10
3e+10
4e+10
5e+10
6e+10
0e+00
2e−11
4e−11
6e−11
8e−11
1e−10
N = 429921 Bandwidth = 4.229e−13

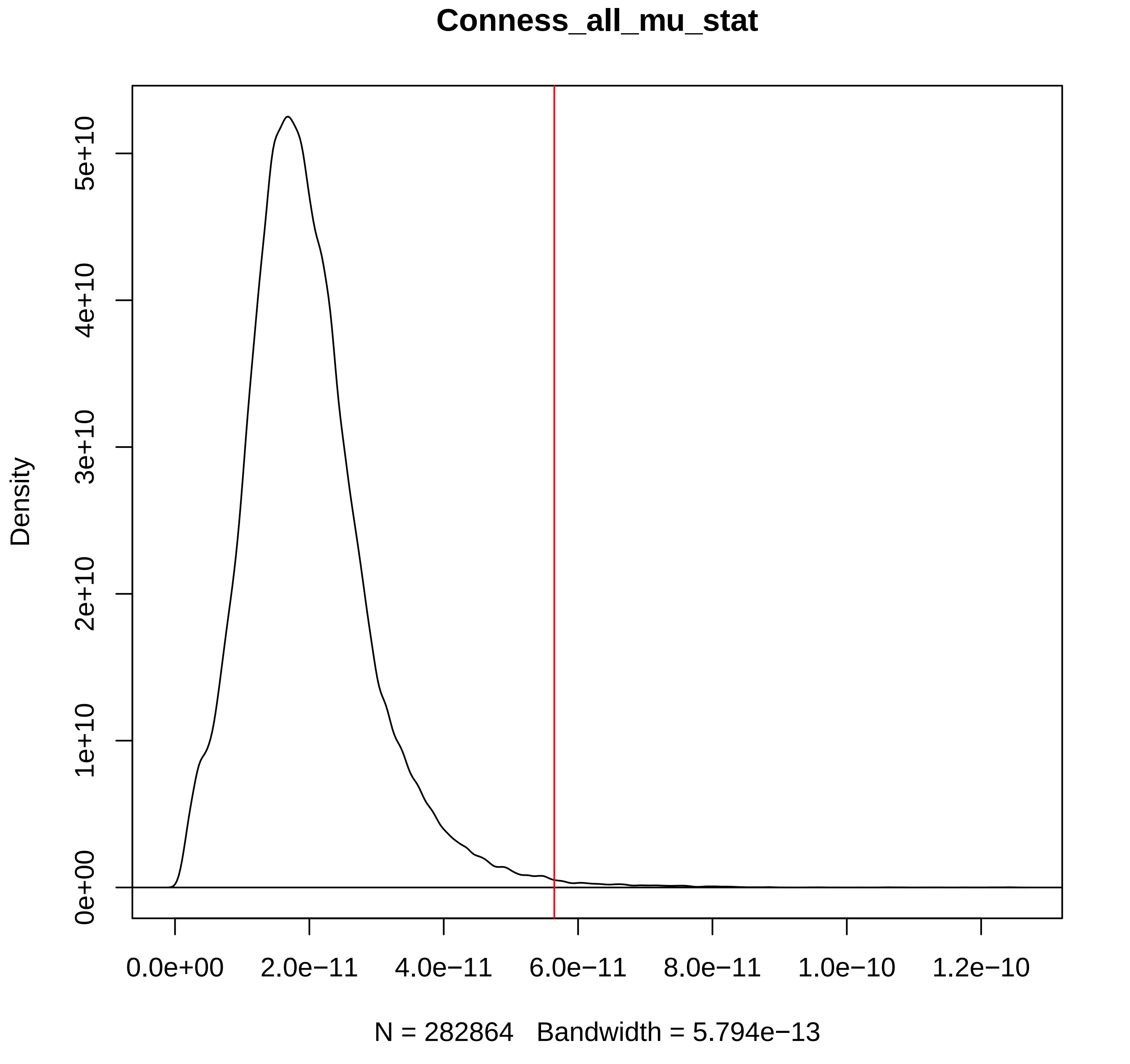
Conness_all_mu_stat
Density
0e+00
1e+10
2e+10
3e+10
4e+10
5e+10
0.0e+00
2.0e−11
4.0e−11
6.0e−11
8.0e−11
1.0e−10
1.2e−10
N = 282864   Bandwidth = 5.794e−13

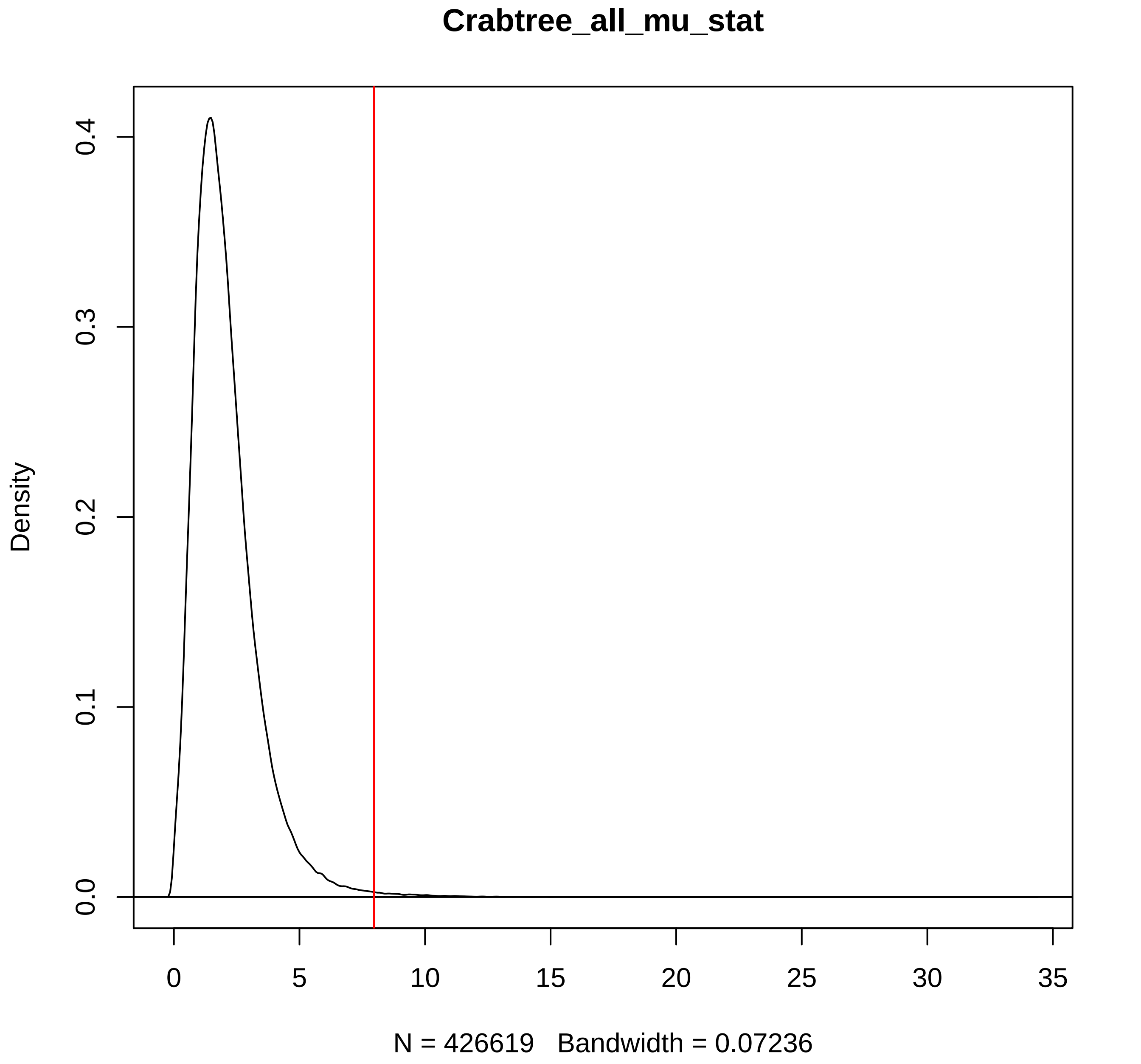
Crabtree_all_mu_stat
Density
0.0
0.1
0.2
0.3
0.4
0
5
10
15
20
25
30
35
N = 426619   Bandwidth = 0.07236

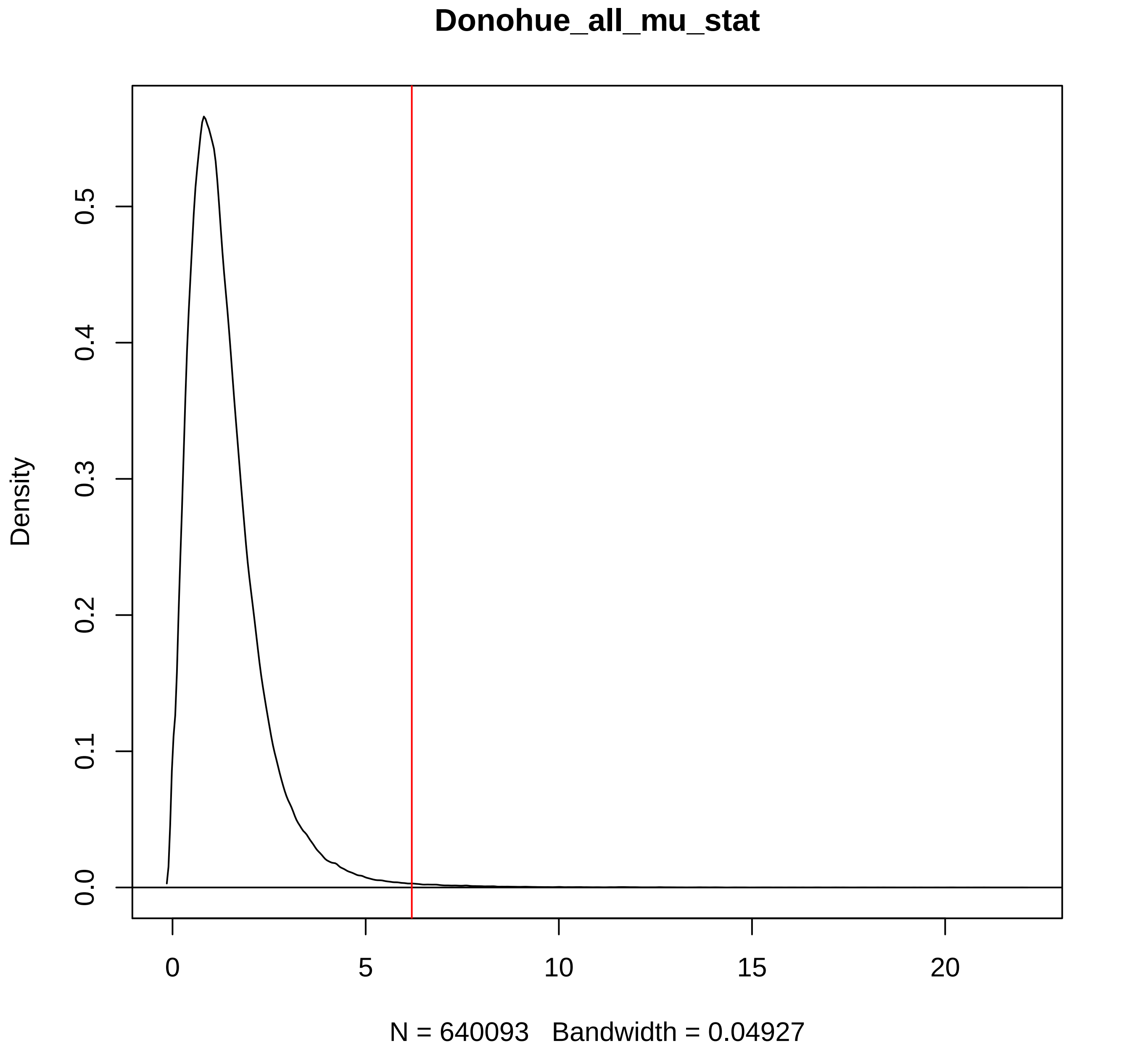
Donohue_all_mu_stat
Density
0.0
0.1
0.2
0.3
0.4
0.5
0
5
10
15
20
N = 640093   Bandwidth = 0.04927

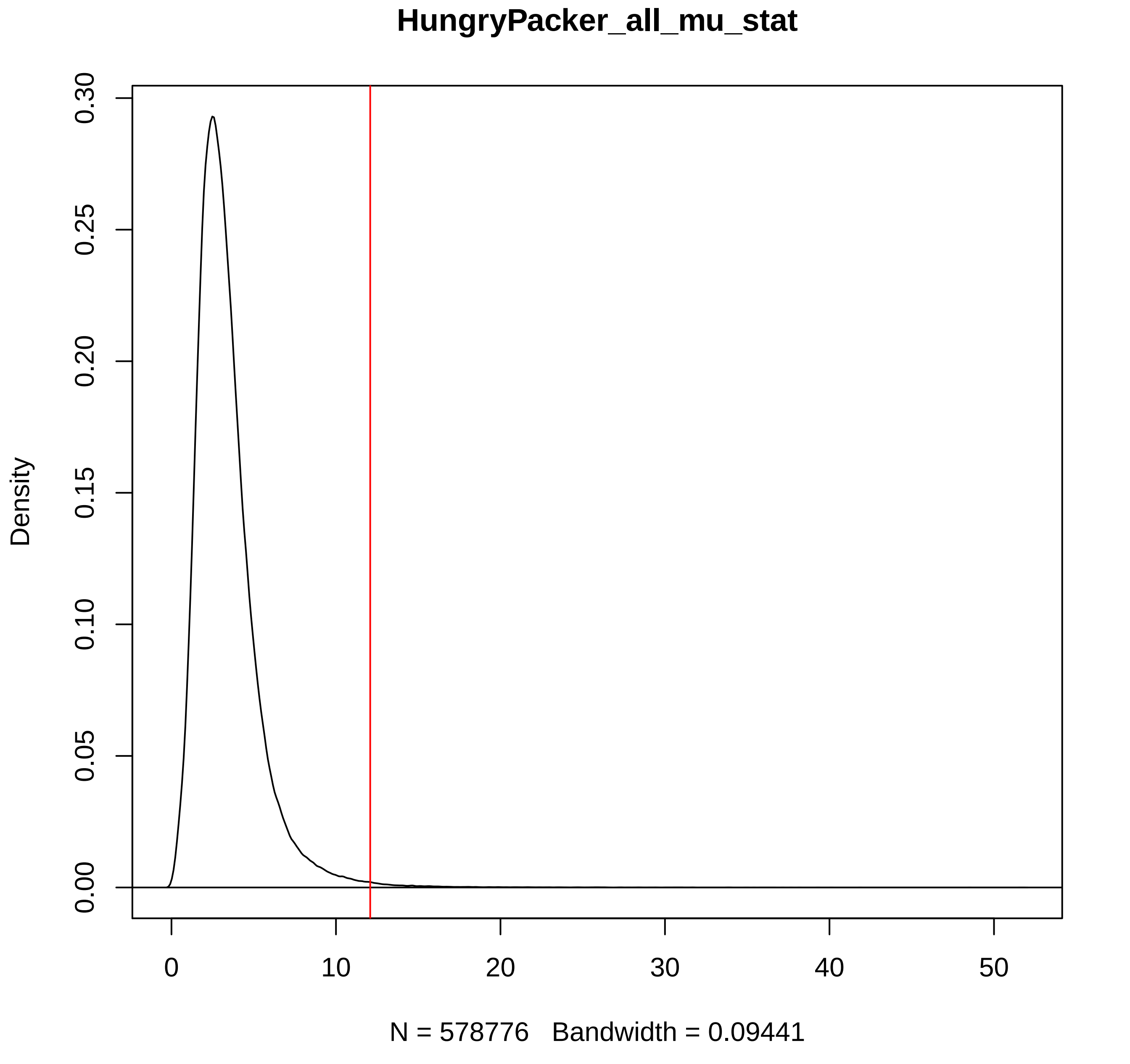
HungryPacker_all_mu_stat
Density
0.00
0.05
0.10
0.15
0.20
0.25
0.30
0
10
20
30
40
50
N = 578776   Bandwidth = 0.09441

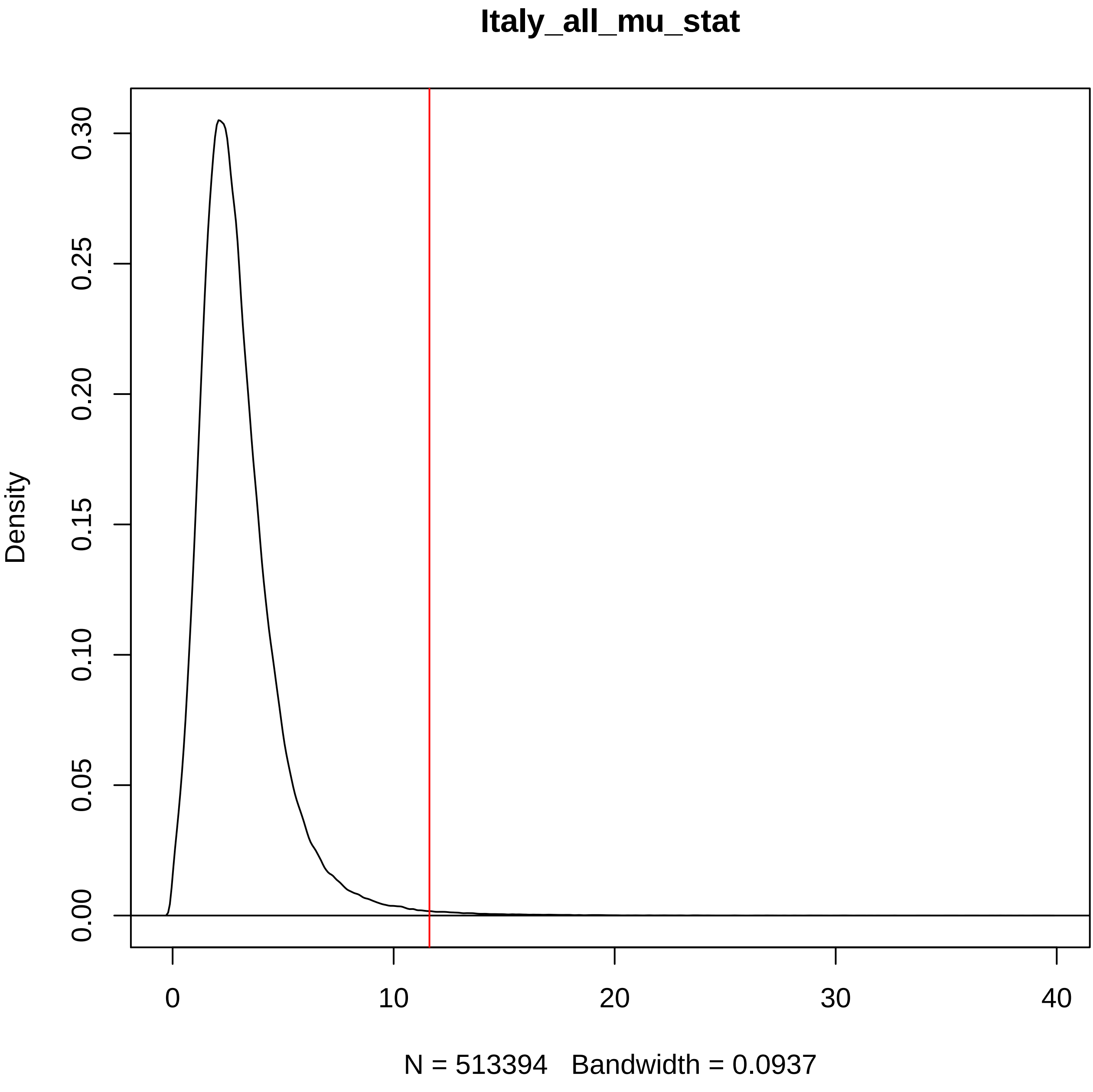
Italy_all_mu_stat
Density
0.00
0.05
0.10
0.15
0.20
0.25
0.30
0
10
20
30
40
N = 513394   Bandwidth = 0.0937

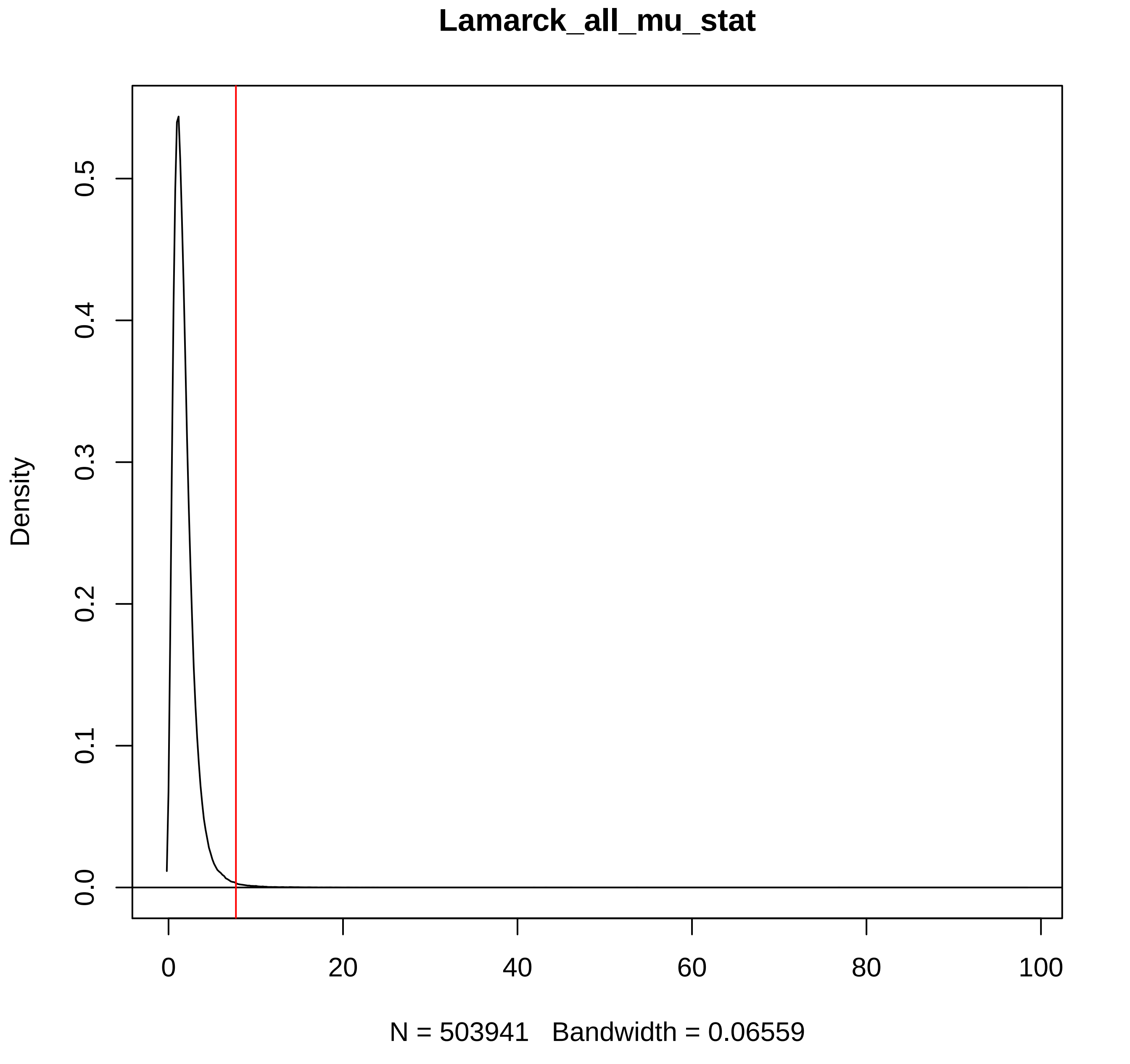

Lamarck_all_mu_stat
Density
0.0
0.1
0.2
0.3
0.4
0.5
0
20
40
60
80
100
N = 503941 Bandwidth = 0.06559

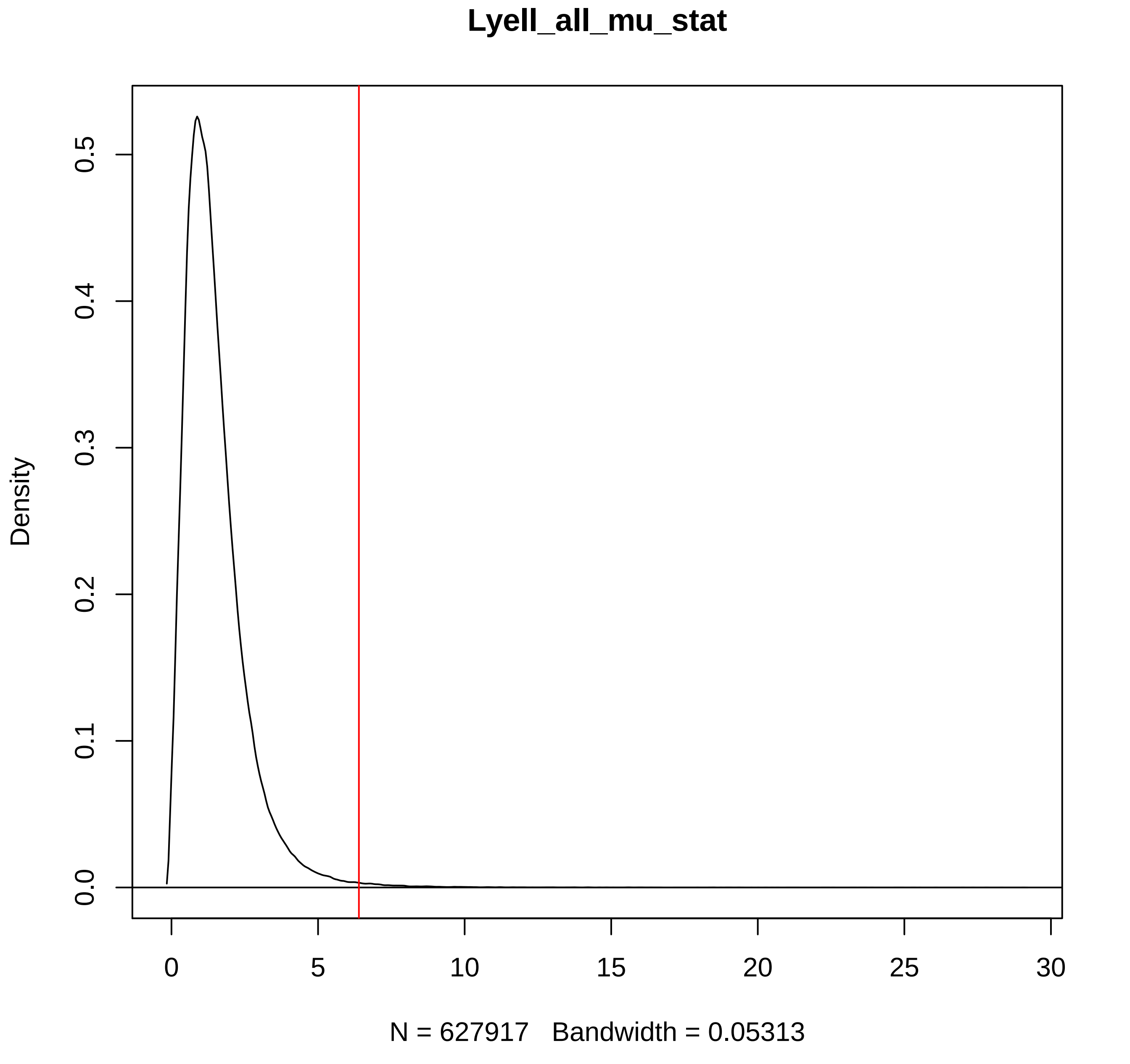

Lyell_all_mu_stat
Density
0.0
0.1
0.2
0.3
0.4
0.5
0
5
10
15
20
25
30
N = 627917 Bandwidth = 0.05313

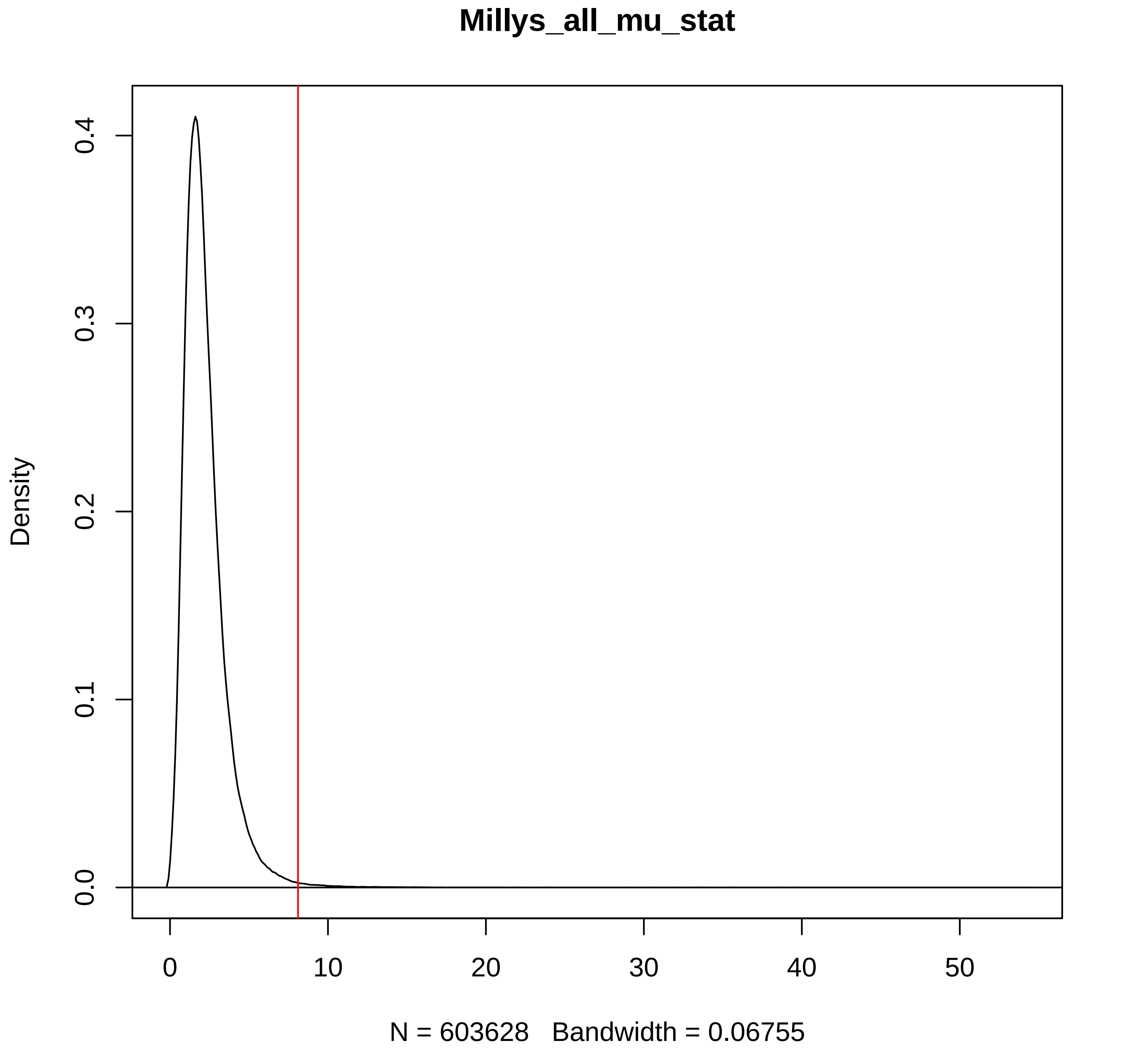

Millys_all_mu_stat
Density
0.0
0.1
0.2
0.3
0.4
0
10
20
30
40
50
N = 603628   Bandwidth = 0.06755

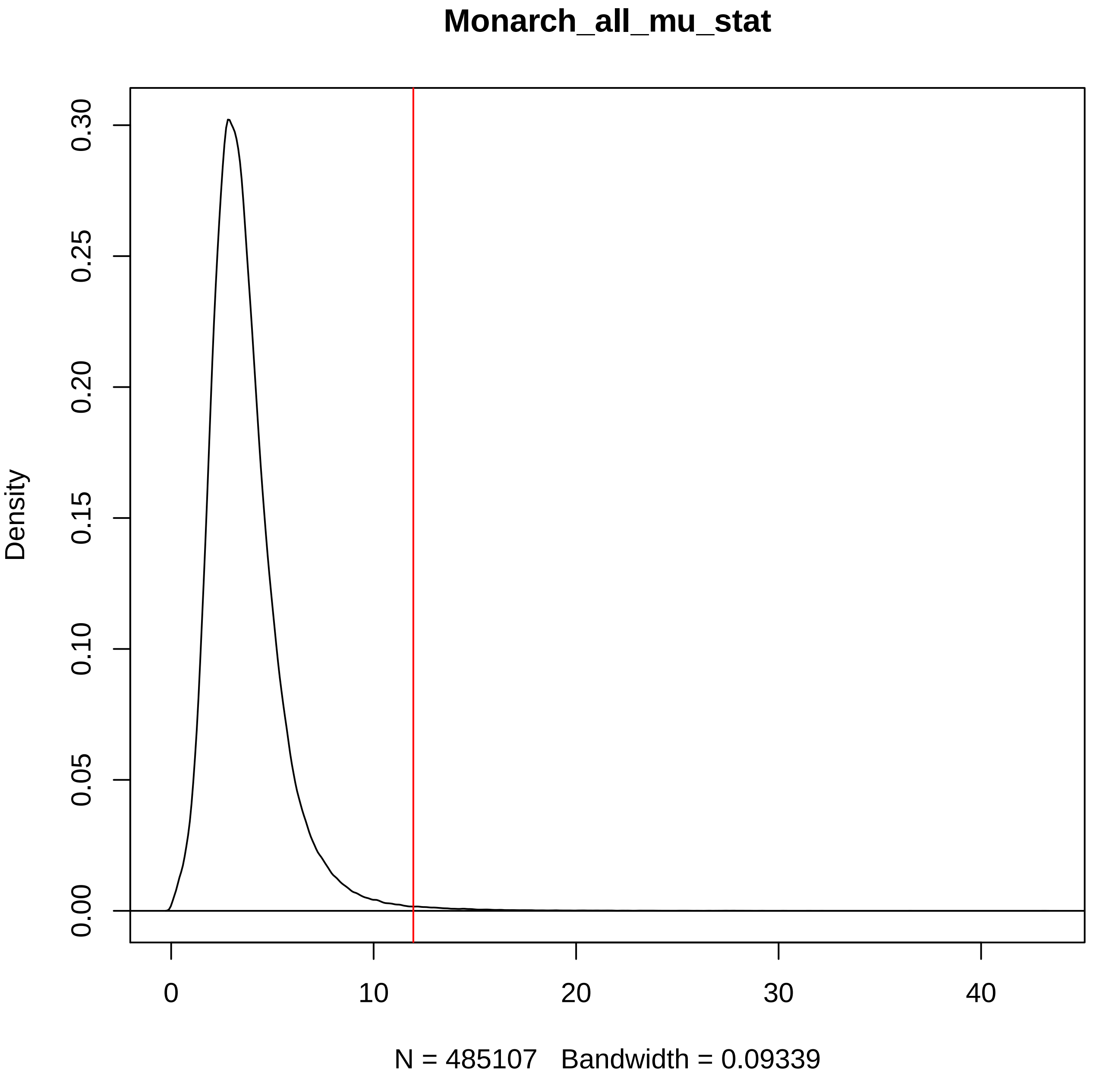

Monarch_all_mu_stat
Density
0.00
0.05
0.10
0.15
0.20
0.25
0.30
0
10
20
30
40
N = 485107   Bandwidth = 0.09339

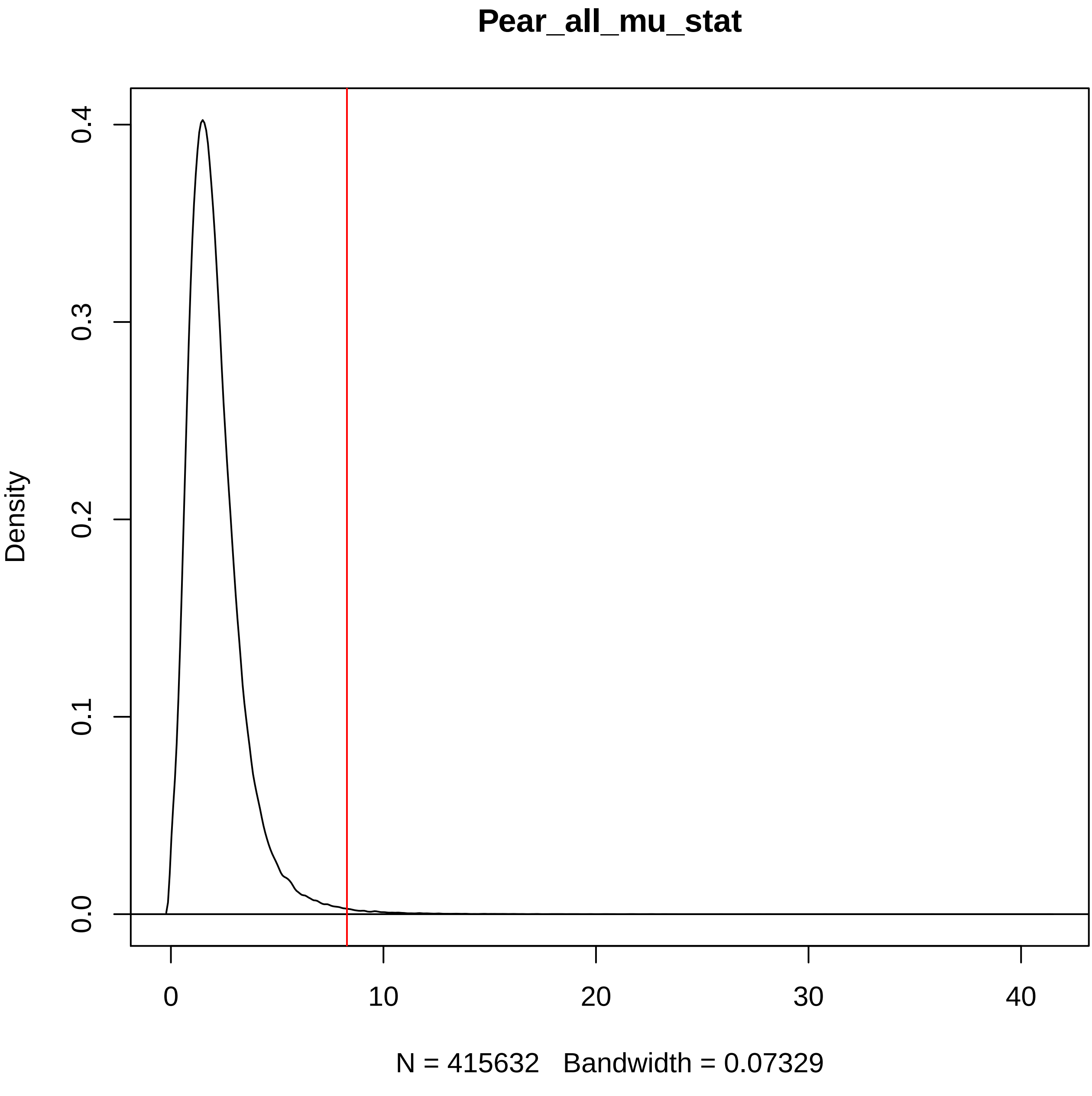

Pear_all_mu_stat
Density
0.0
0.1
0.2
0.3
0.4
0
10
20
30
40
N = 415632   Bandwidth = 0.07329

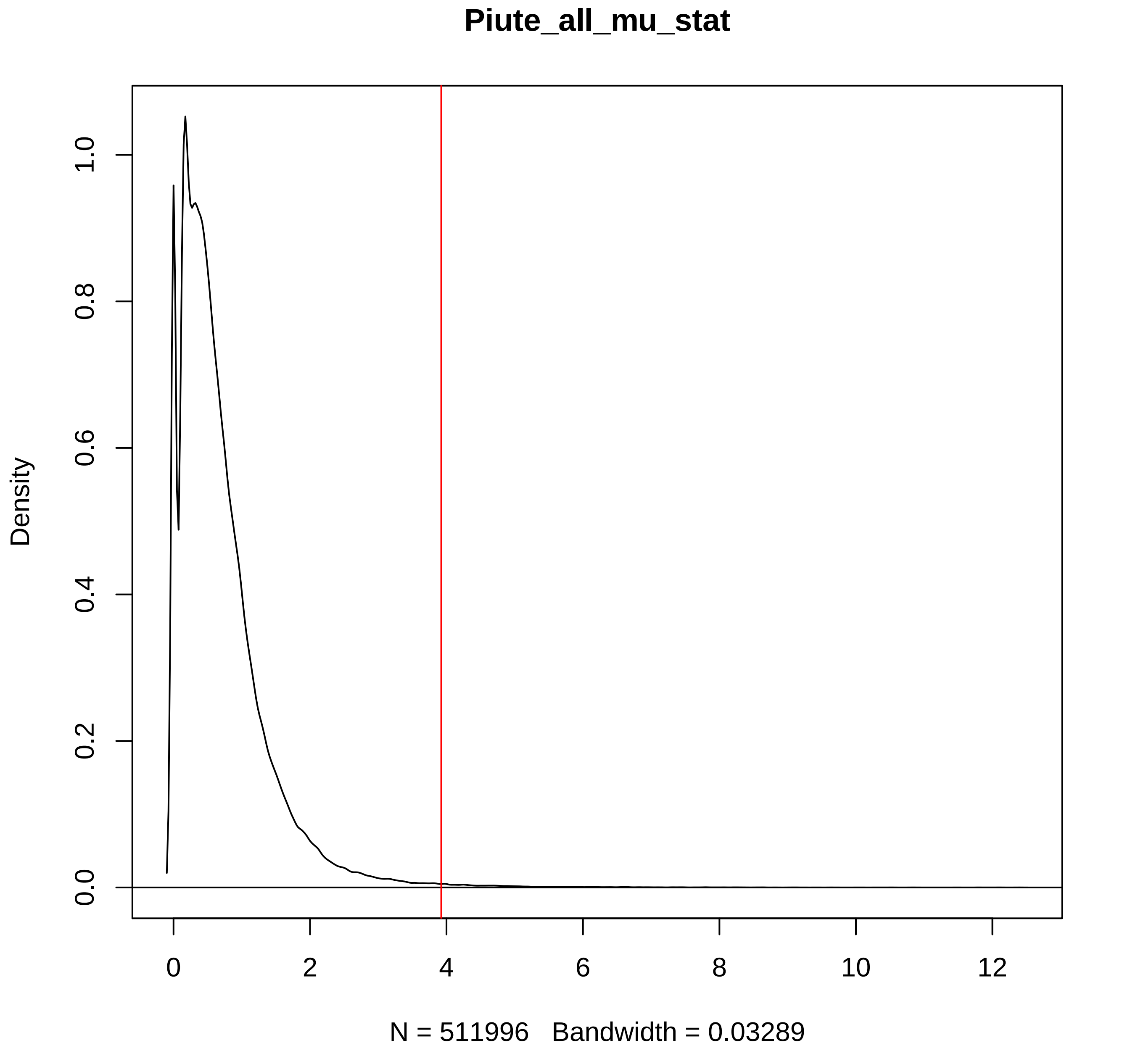
Piute_all_mu_stat
Density
0.0
0.2
0.4
0.6
0.8
1.0
0
2
4
6
8
10
12
N = 511996   Bandwidth = 0.03289

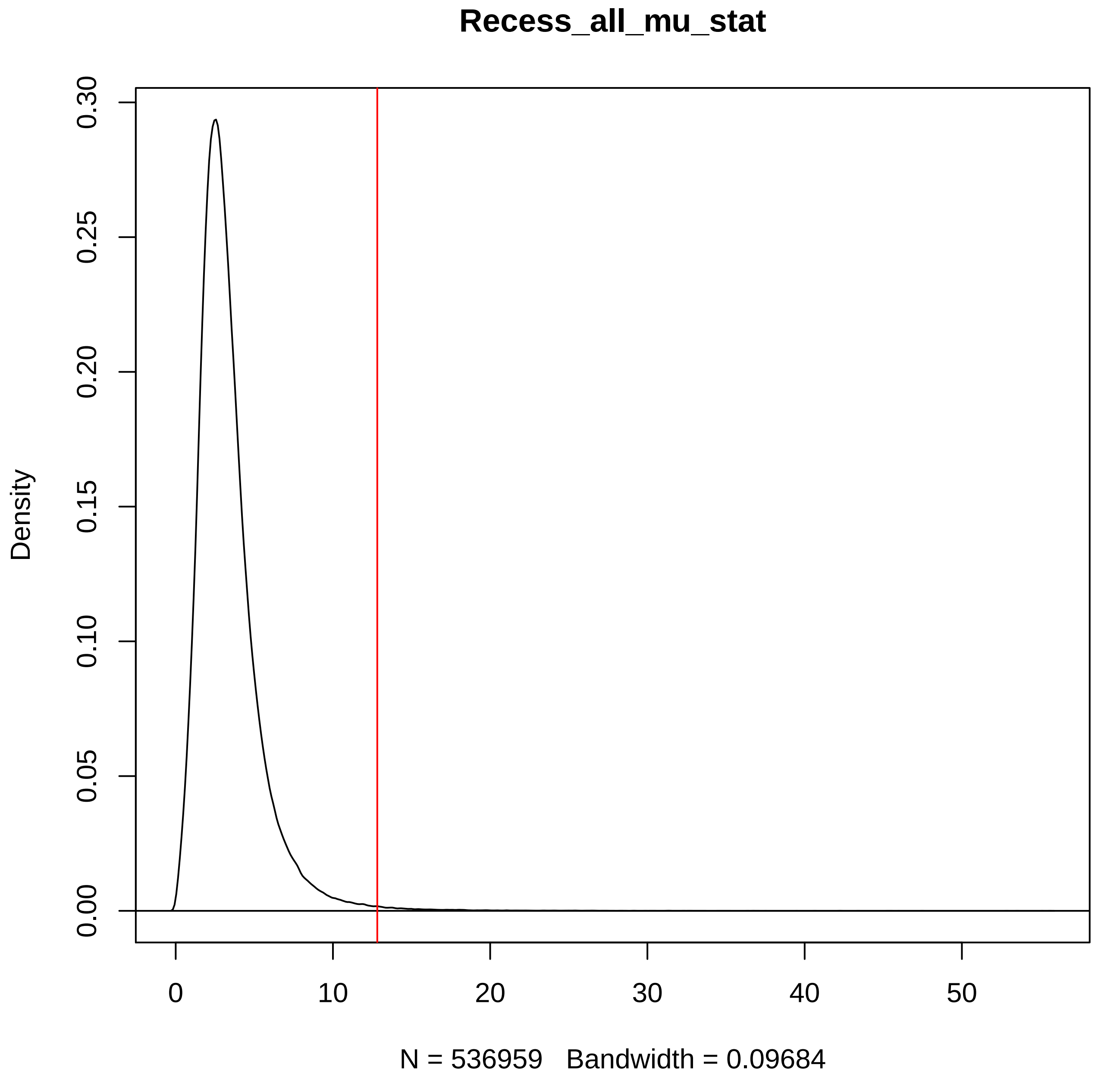
Recess_all_mu_stat
Density
0.00
0.05
0.10
0.15
0.20
0.25
0.30
0
10
20
30
40
50
N = 536959   Bandwidth = 0.09684

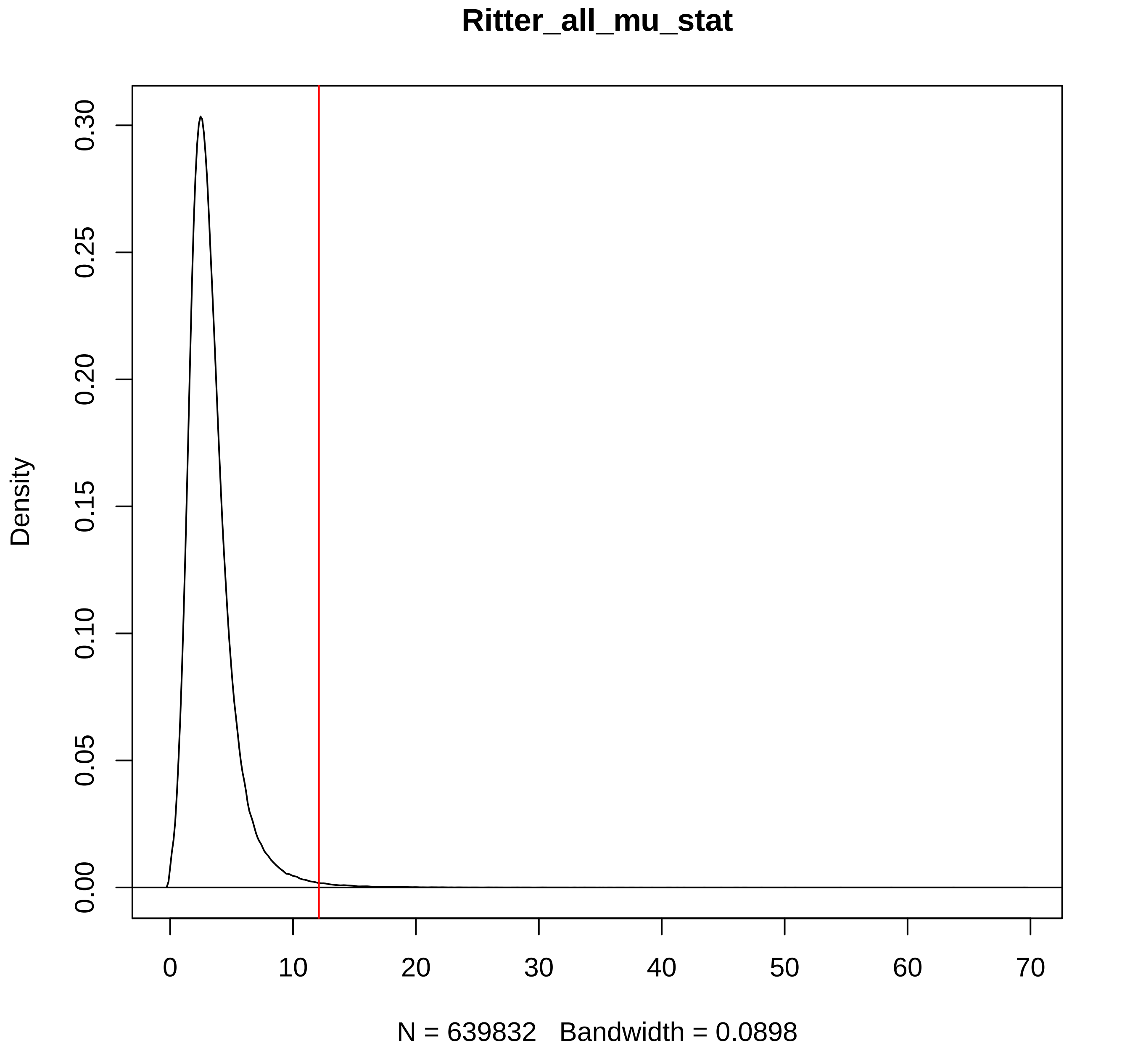
Ritter_all_mu_stat
Density
0.00
0.05
0.10
0.15
0.20
0.25
0.30
0
10
20
30
40
50
60
70
N = 639832   Bandwidth = 0.0898

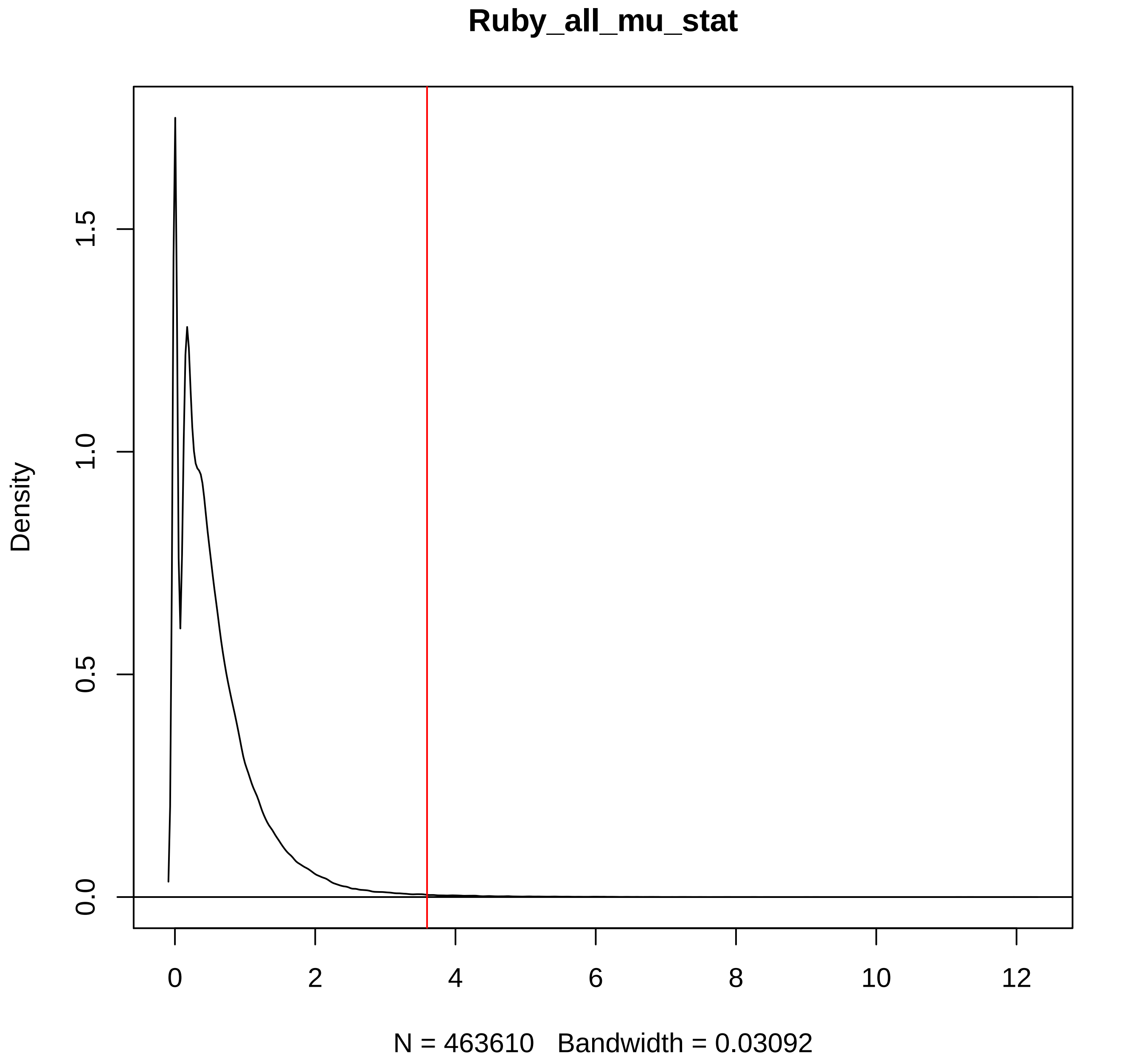

Ruby_all_mu_stat
Density
0.0
0.5
1.0
1.5
0
2
4
6
8
10
12
N = 463610 Bandwidth = 0.03092

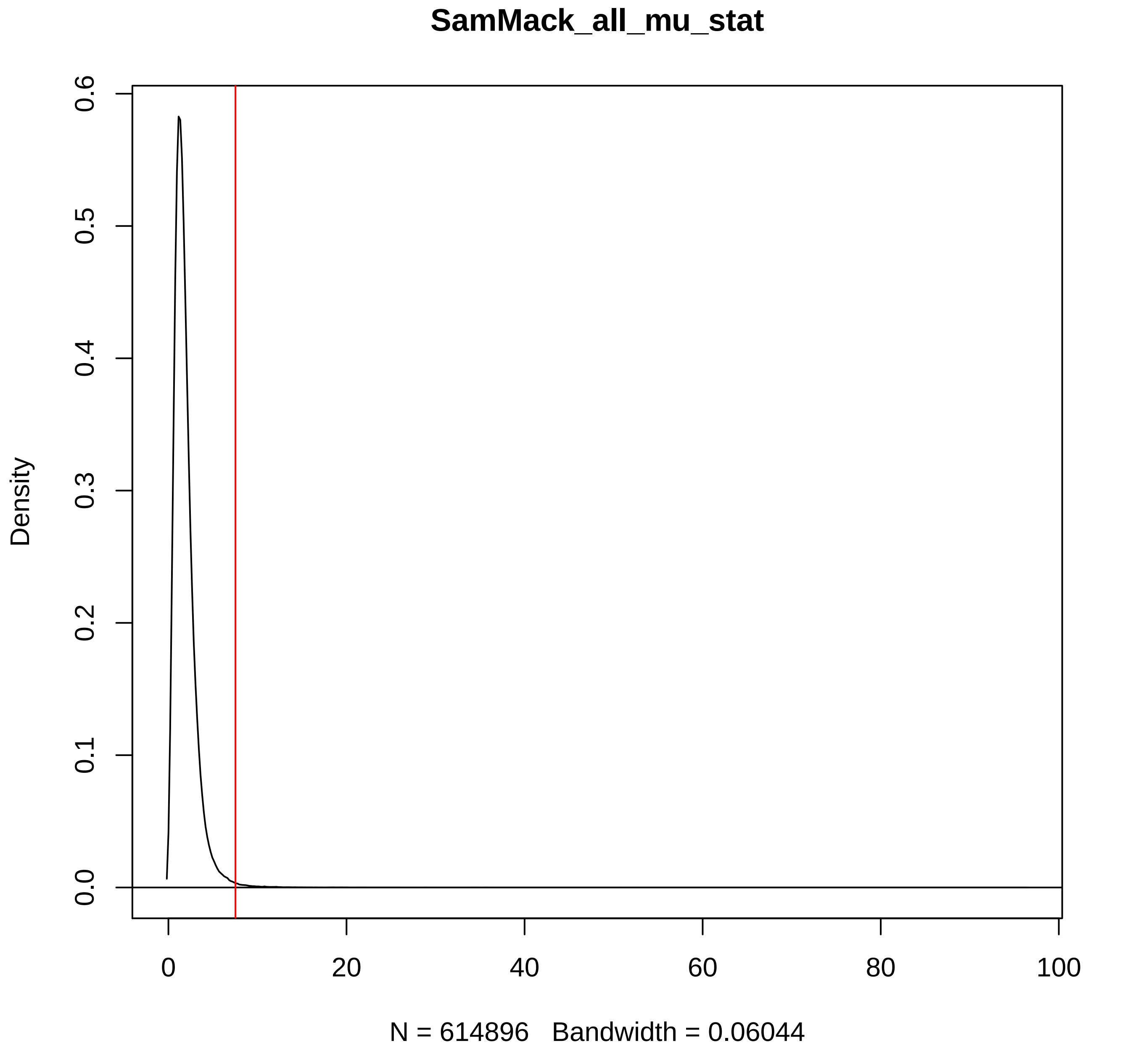

SamMack_all_mu_stat
Density
0.0
0.1
0.2
0.3
0.4
0.5
0.6
0
20
40
60
80
100
N = 614896 Bandwidth = 0.06044

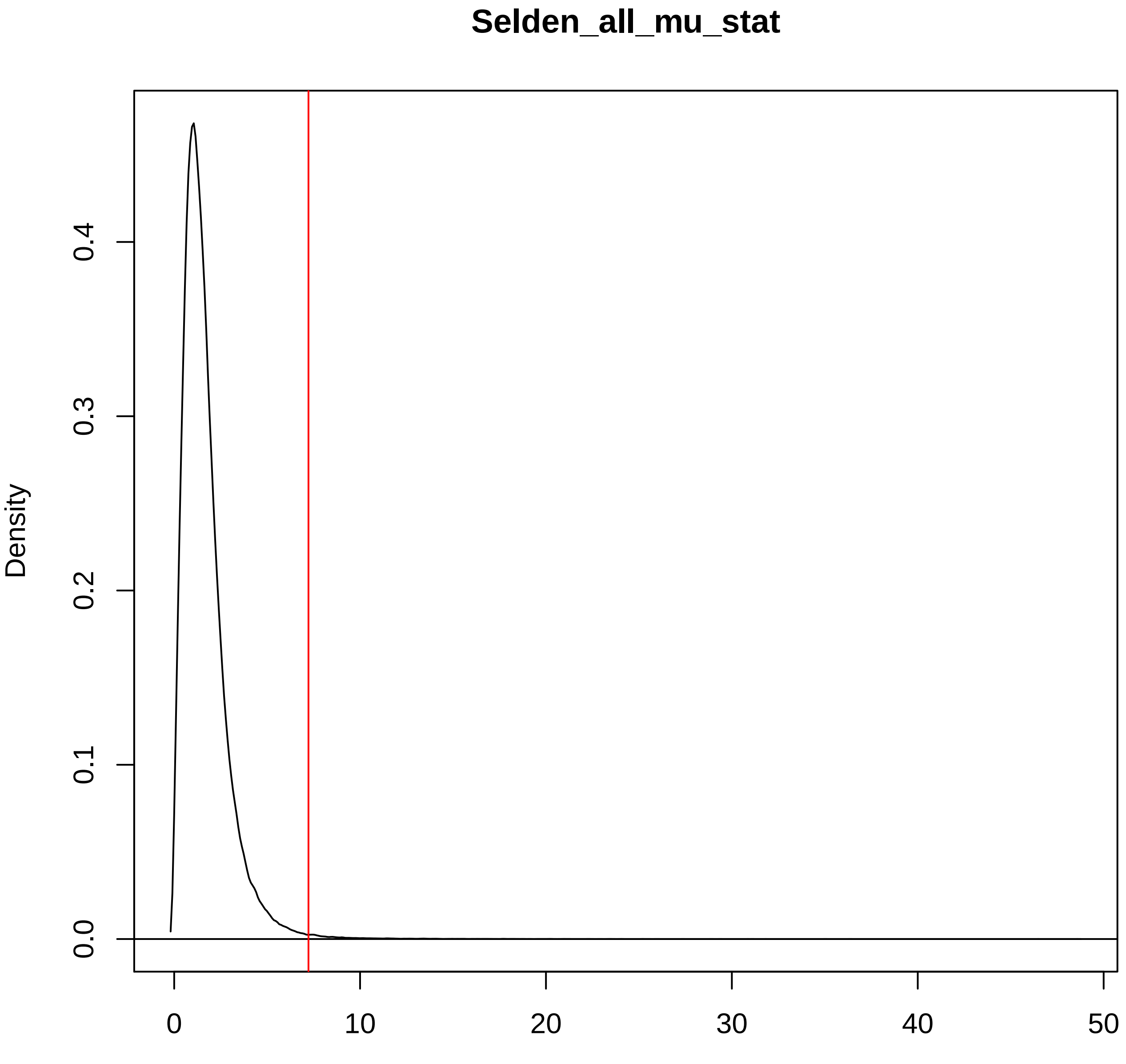

Selden_all_mu_stat
Density
0.0
0.1
0.2
0.3
0.4
0
10
20
30
40
50
N = 452781   Bandwidth = 0.06399

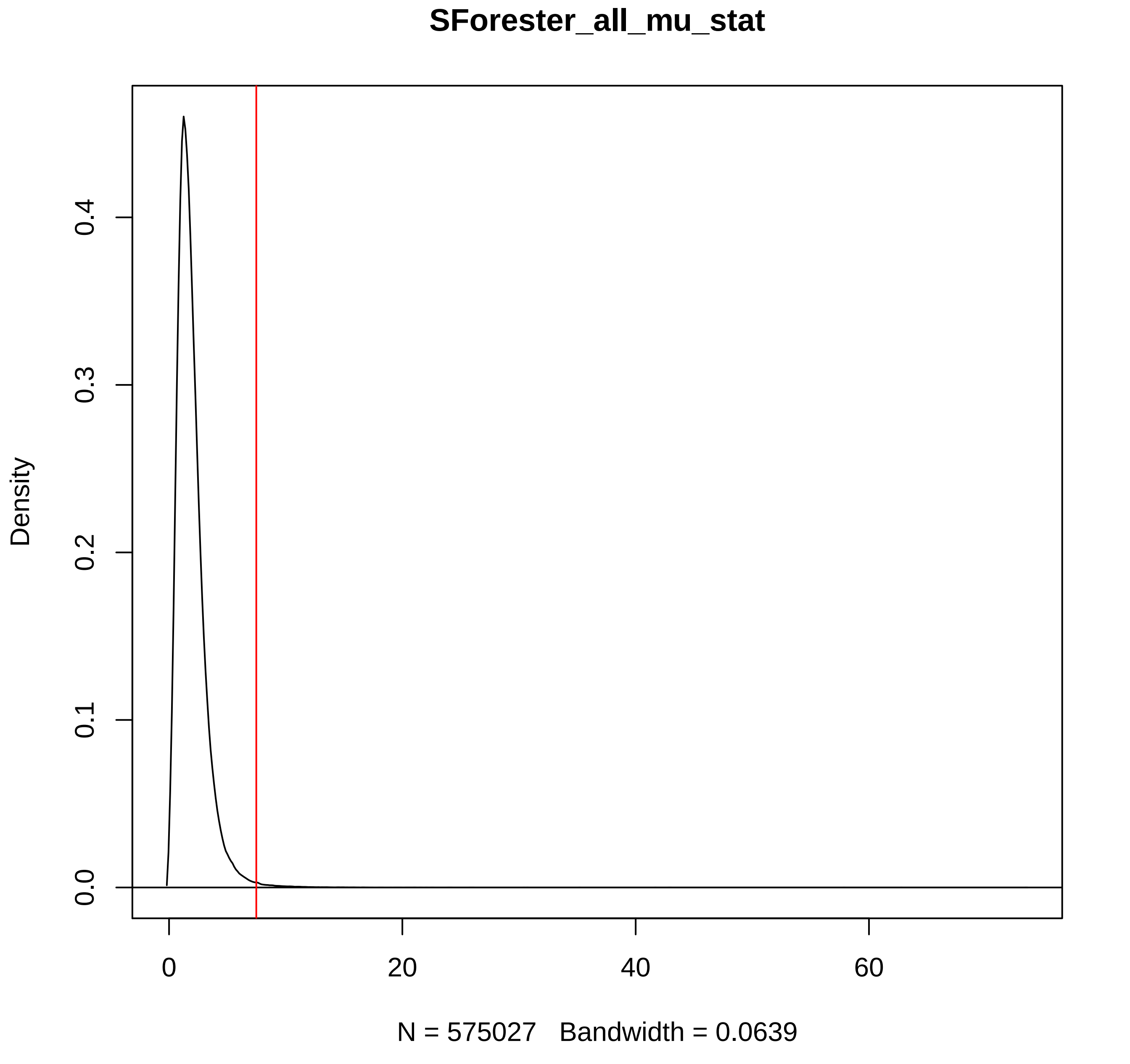
SForester_all_mu_stat
Density
0.0
0.1
0.2
0.3
0.4
0
20
40
60
N = 575027 Bandwidth = 0.0639

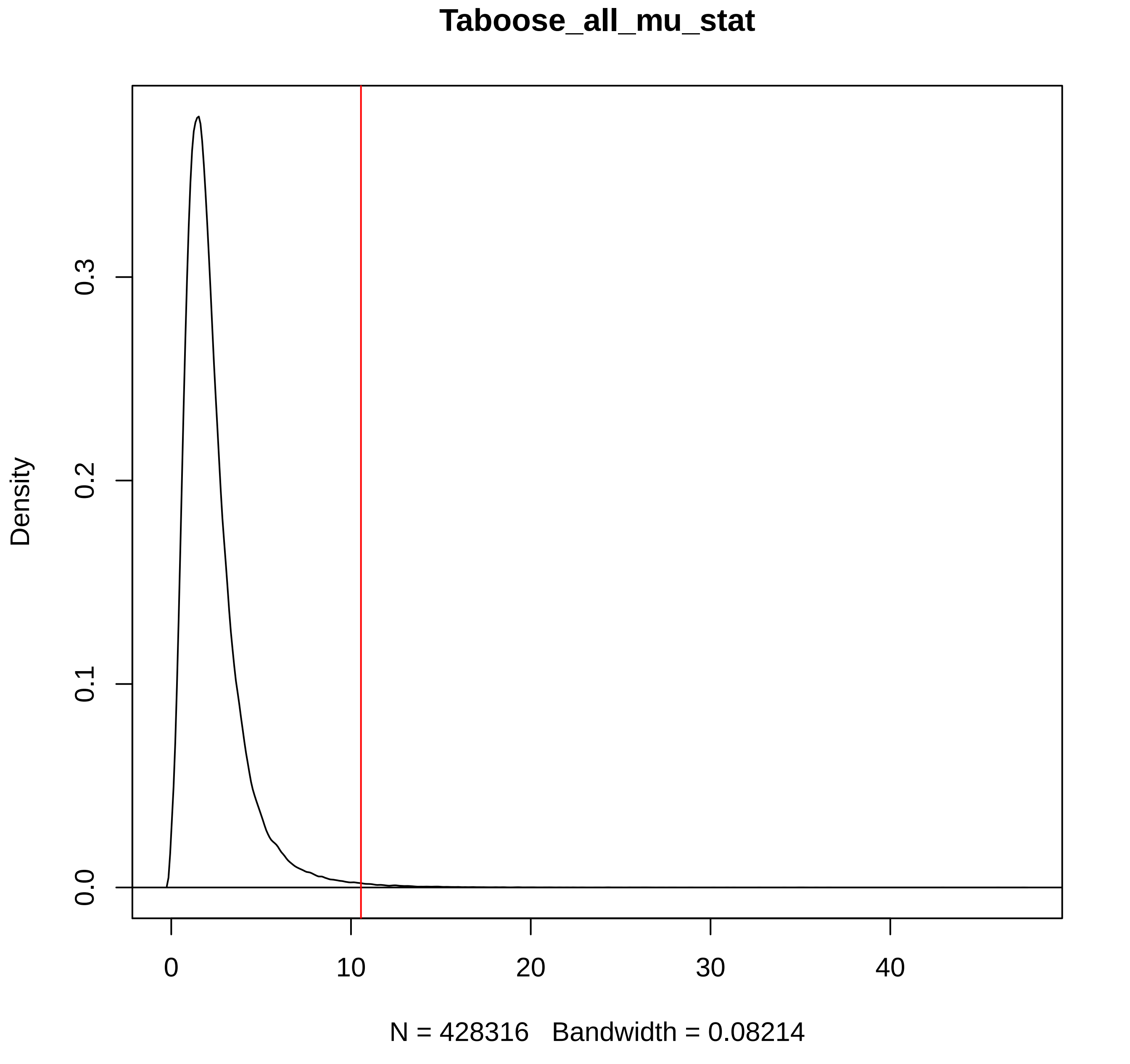
Taboose_all_mu_stat
Density
0.0
0.1
0.2
0.3
0
10
20
30
40
N = 428316   Bandwidth = 0.08214

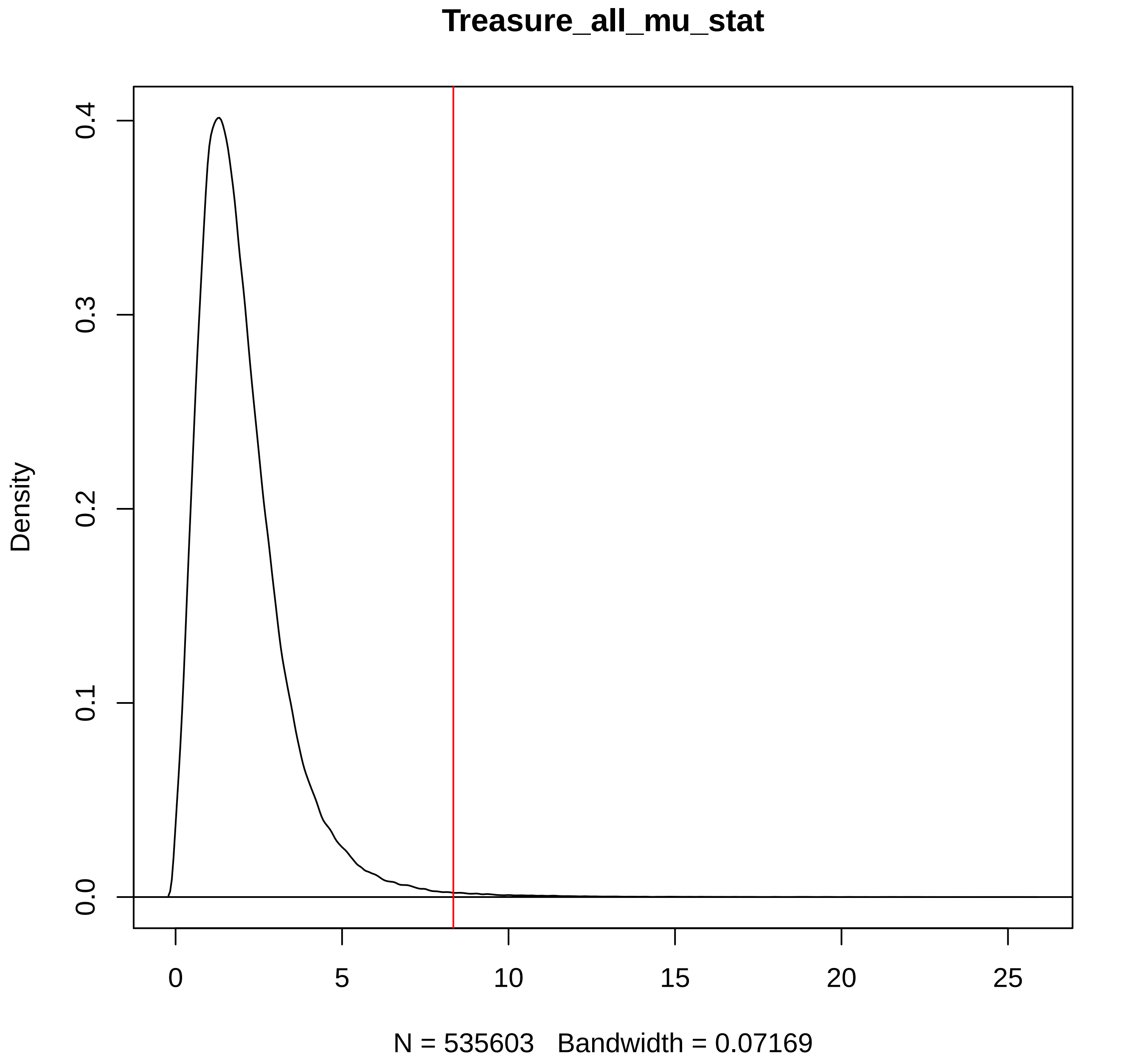
Treasure_all_mu_stat
Density
0.0
0.1
0.2
0.3
0.4
0
5
10
15
20
25
N = 535603   Bandwidth = 0.07169

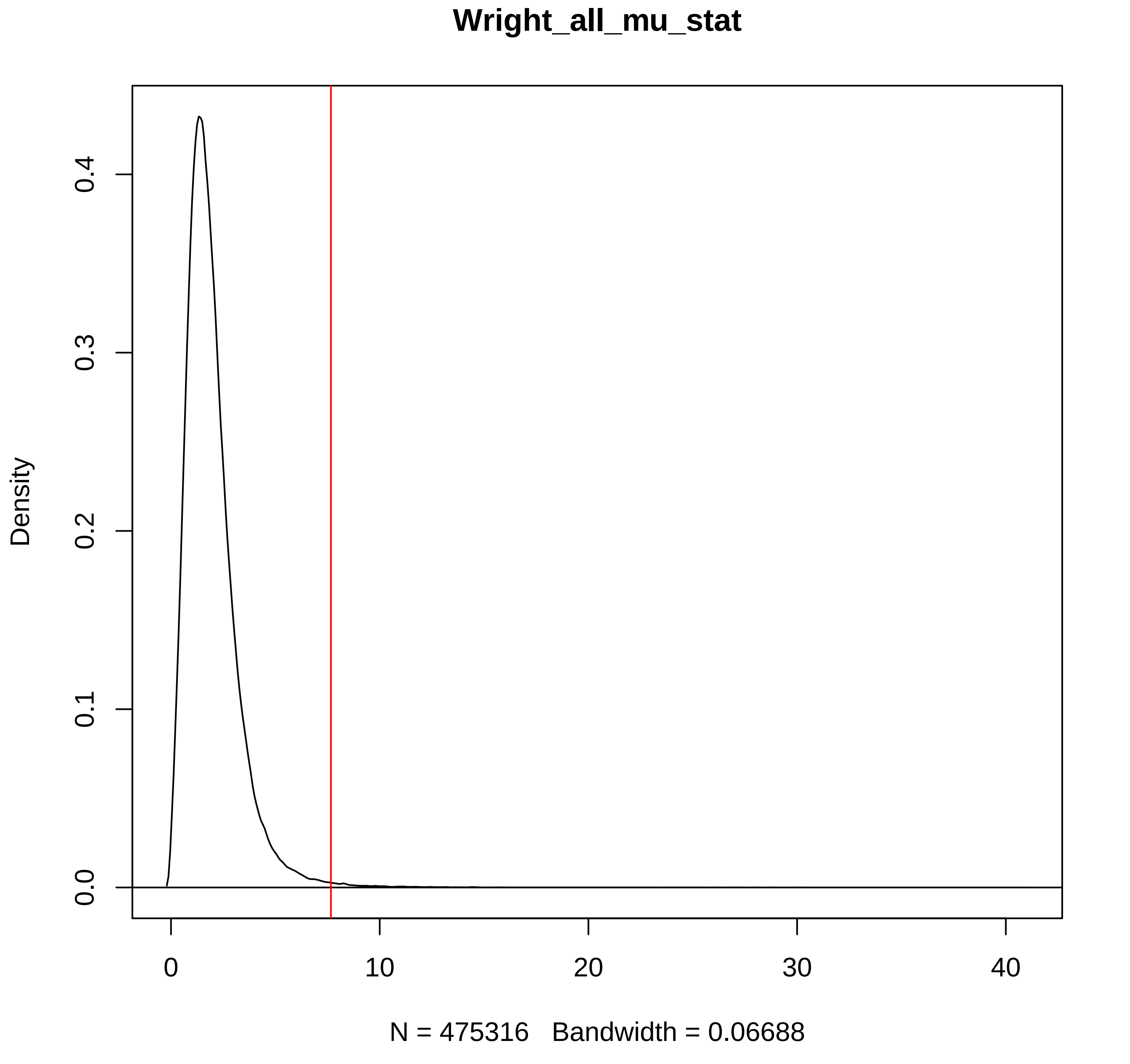

Wright_all_mu_stat
Density
0.0
0.1
0.2
0.3
0.4
0
10
20
30
40
N = 475316   Bandwidth = 0.06688

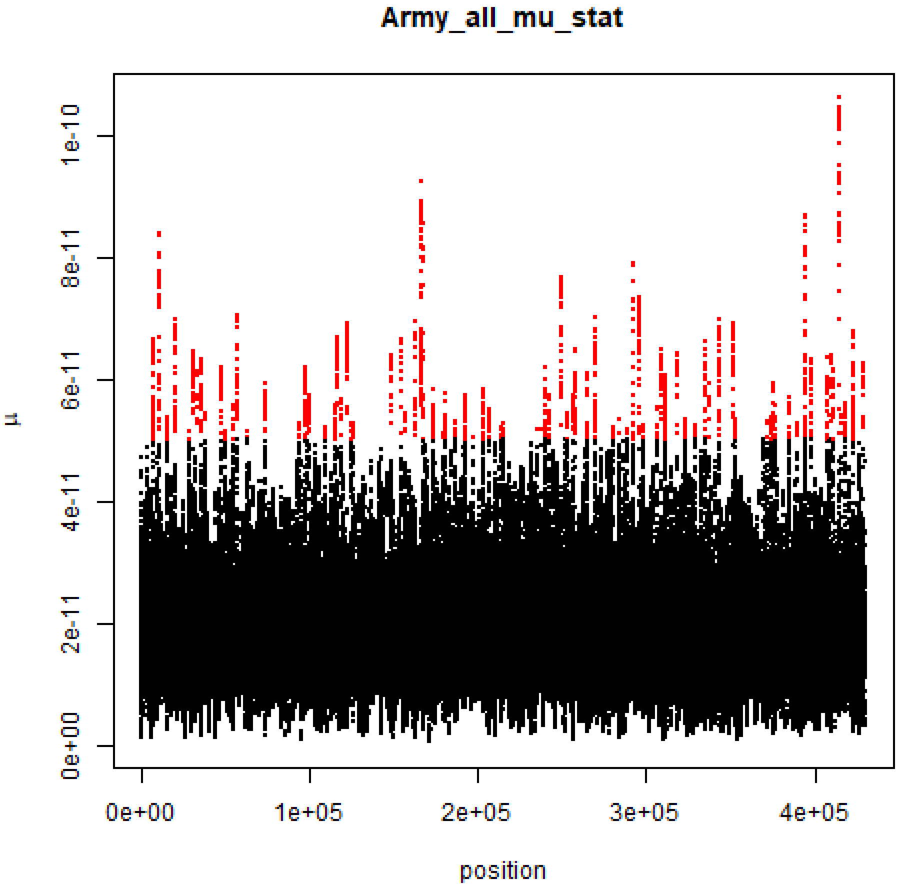
Army_all_mu_stat
μ
1e-10
8e-11
6e-11
4e-11
2e-11
0e+00
0e+00
1e+05
2e+05
3e+05
4e+05
position

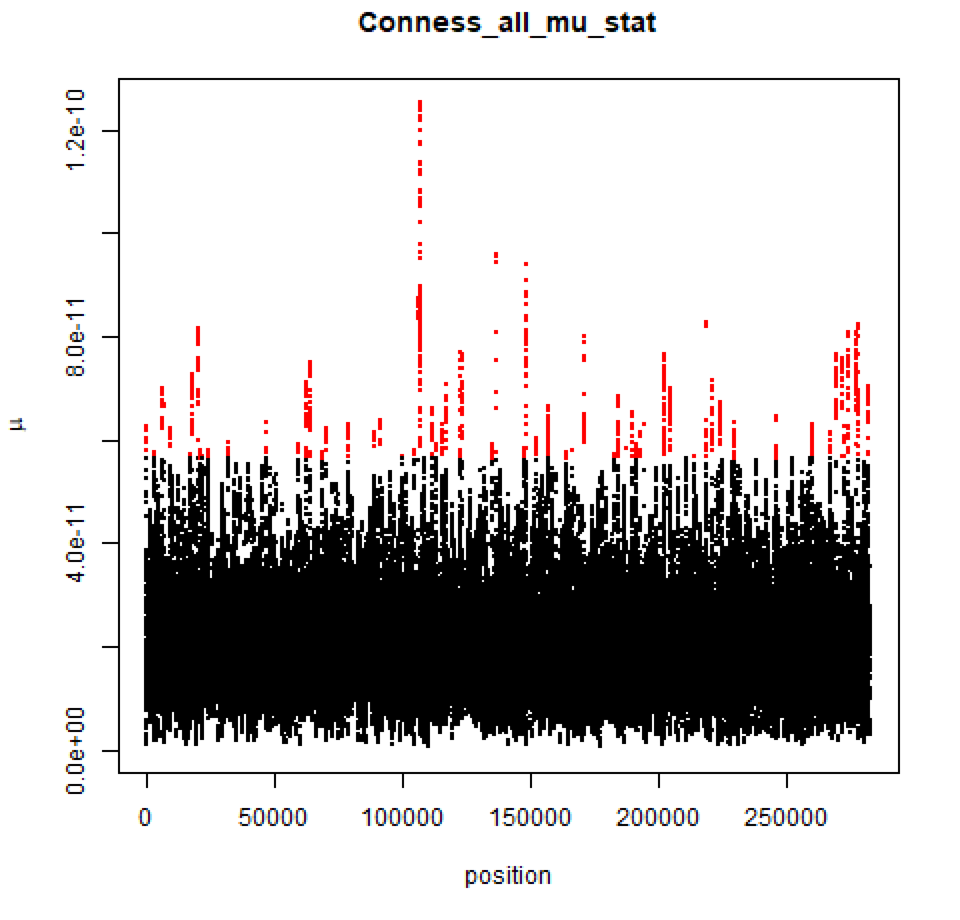
Conness_all_mu_stat
μ
0.0e+00
4.0e-11
8.0e-11
1.2e-10
0
50000
100000
150000
200000
250000
position

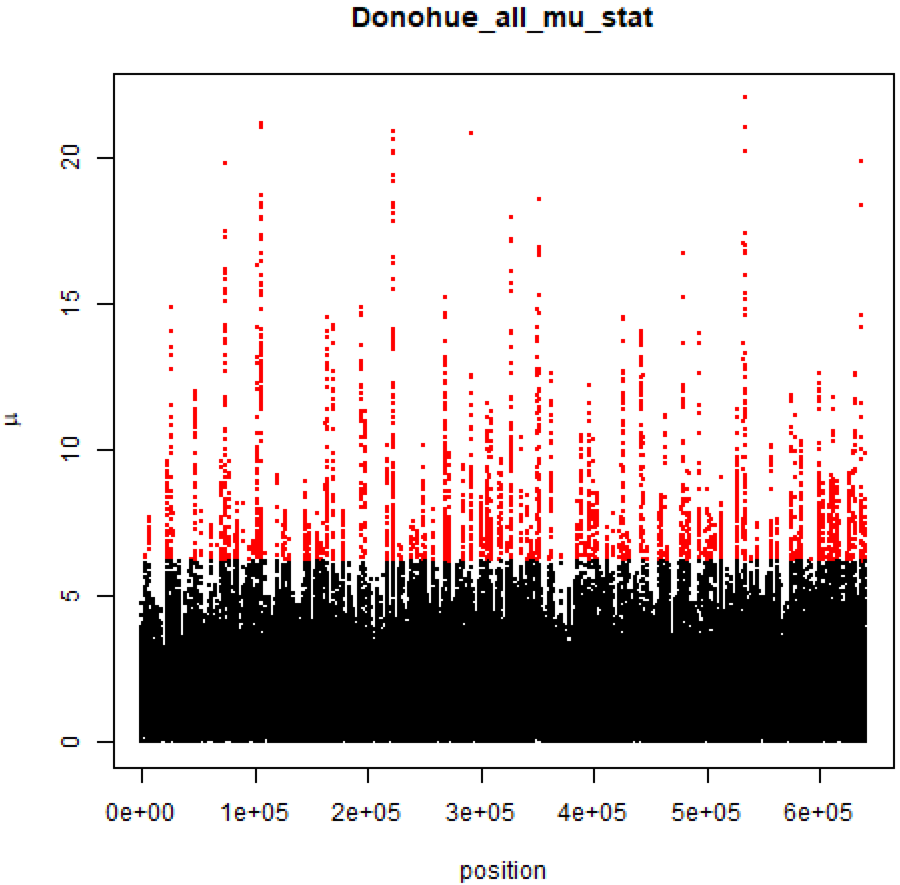

Donohue_all_mu_stat
μ
0
5
10
15
20
0e+00
1e+05
2e+05
3e+05
4e+05
5e+05
6e+05
position

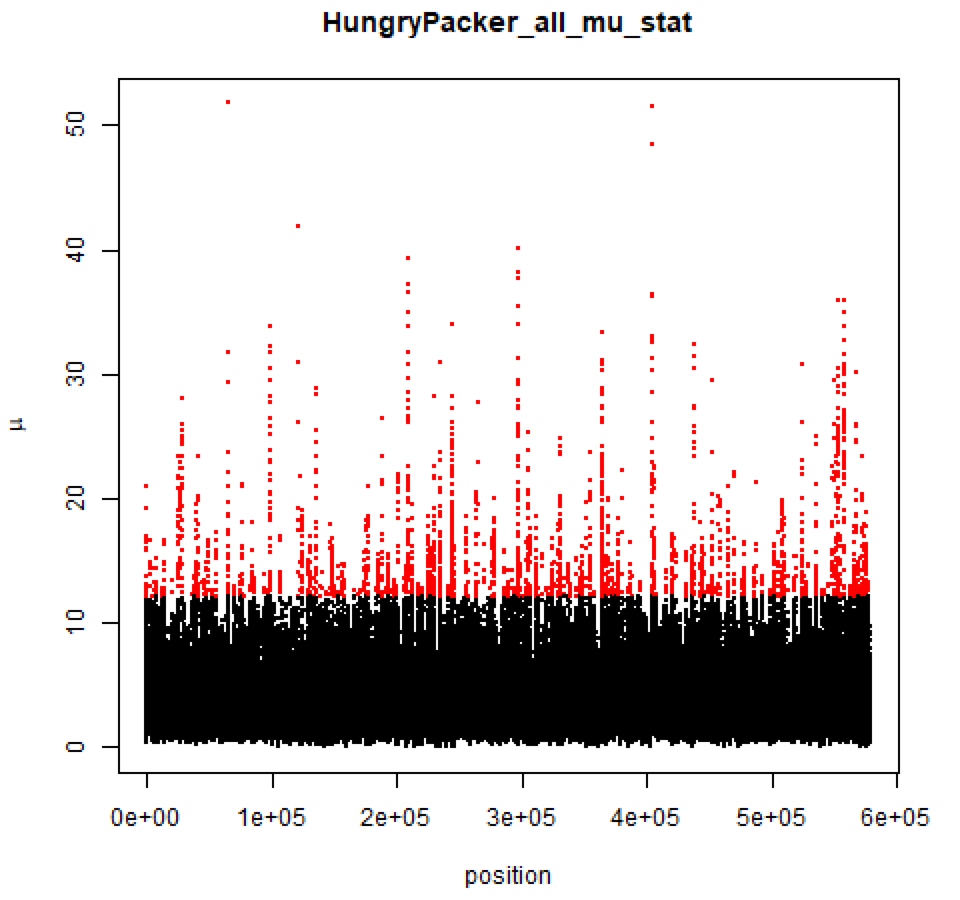

HungryPacker_all_mu_stat
μ
0
10
20
30
40
50
0e+00
1e+05
2e+05
3e+05
4e+05
5e+05
6e+05
position

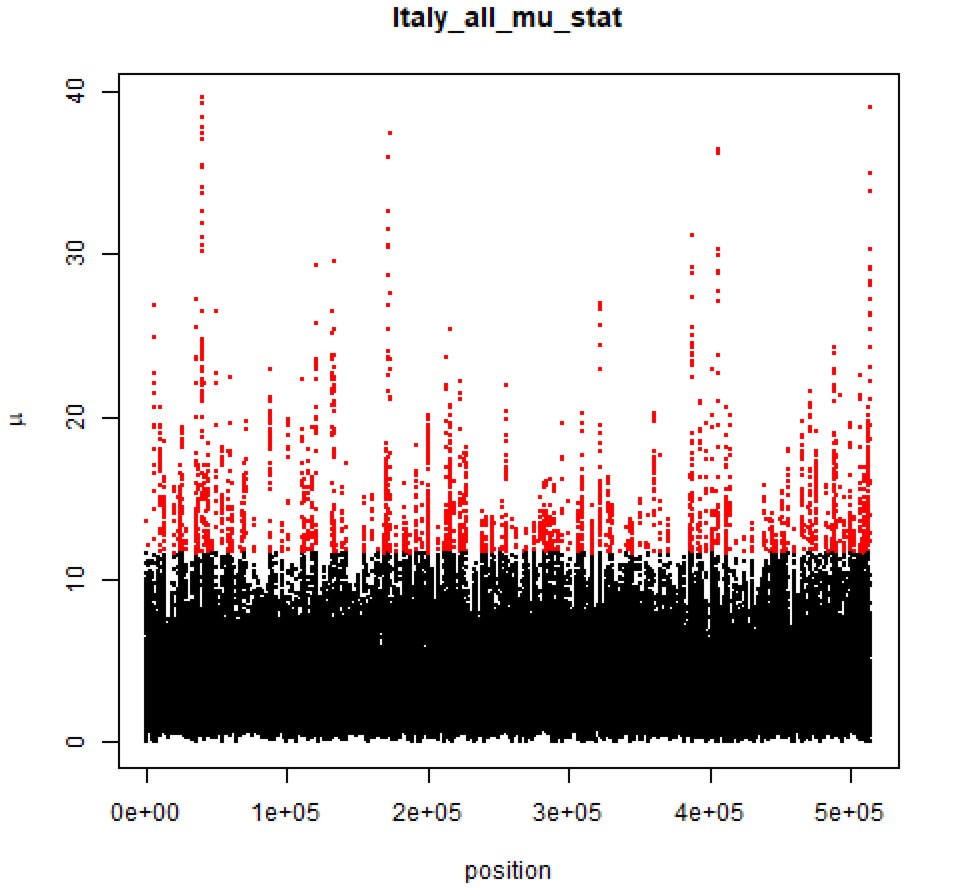

Italy_all_mu_stat
μ
40
30
20
10
0
0e+00
1e+05
2e+05
3e+05
4e+05
5e+05
position

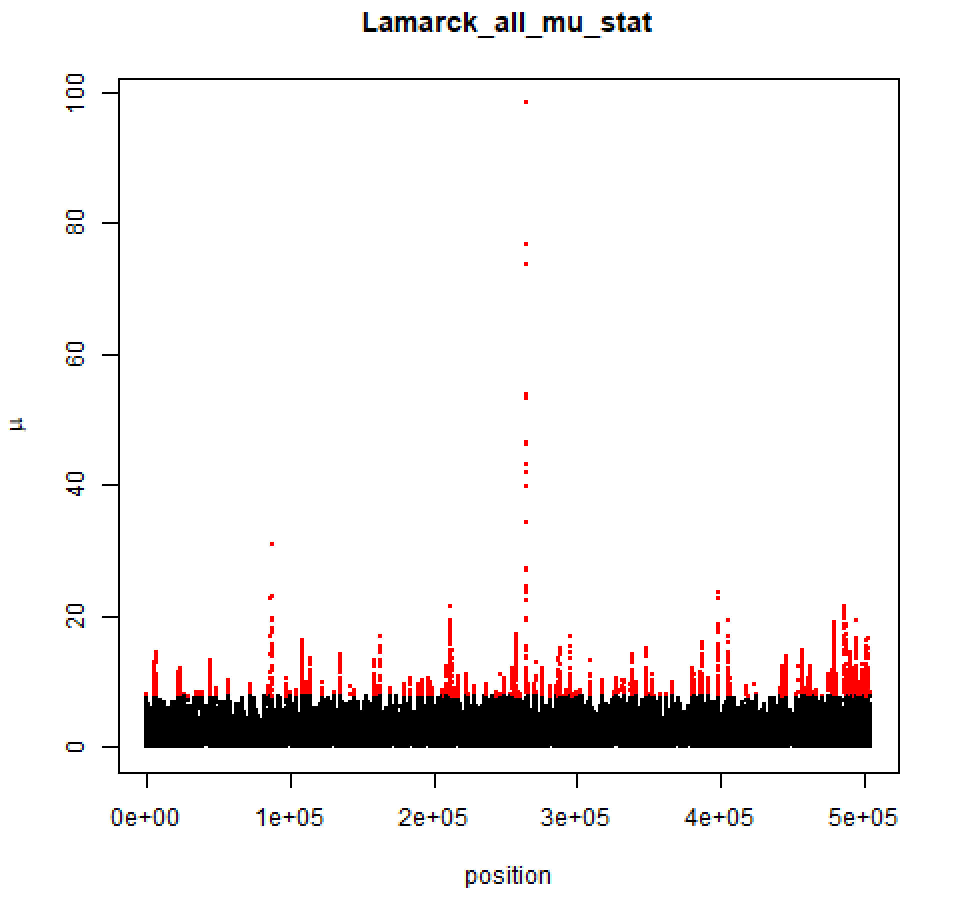

Lamarck_all_mu_stat
μ
position
0
20
40
60
80
100
0e+00
1e+05
2e+05
3e+05
4e+05
5e+05

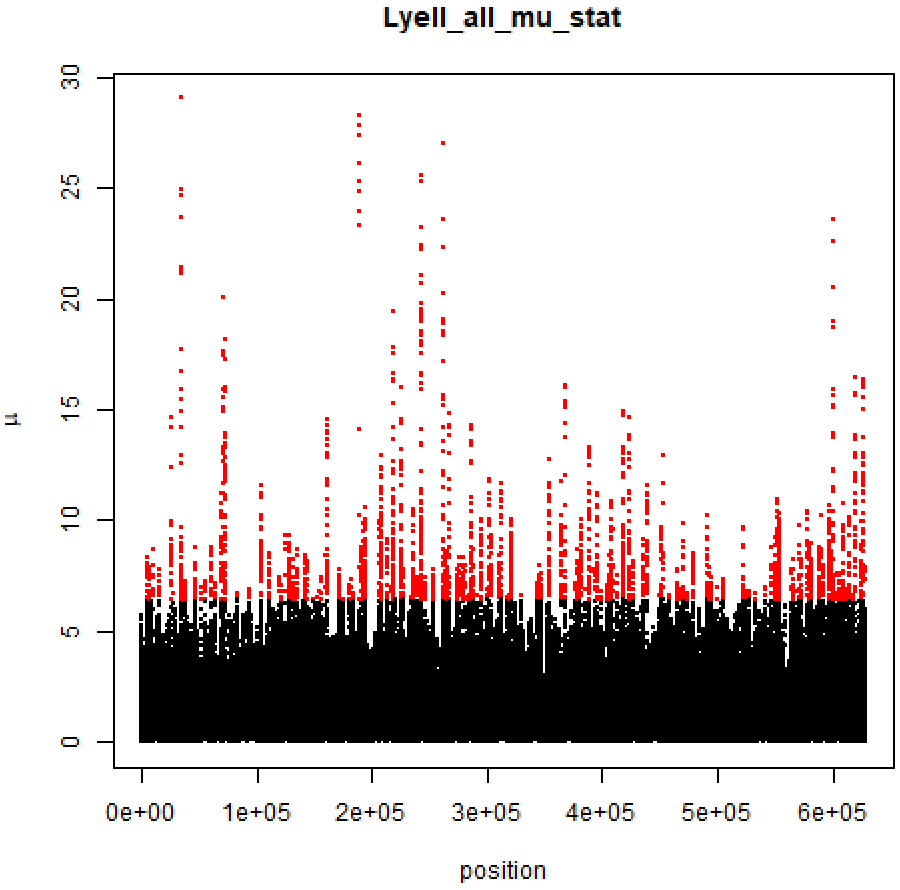

Lyell_all_mu_stat
μ
position
0
5
10
15
20
25
30
0e+00
1e+05
2e+05
3e+05
4e+05
5e+05
6e+05

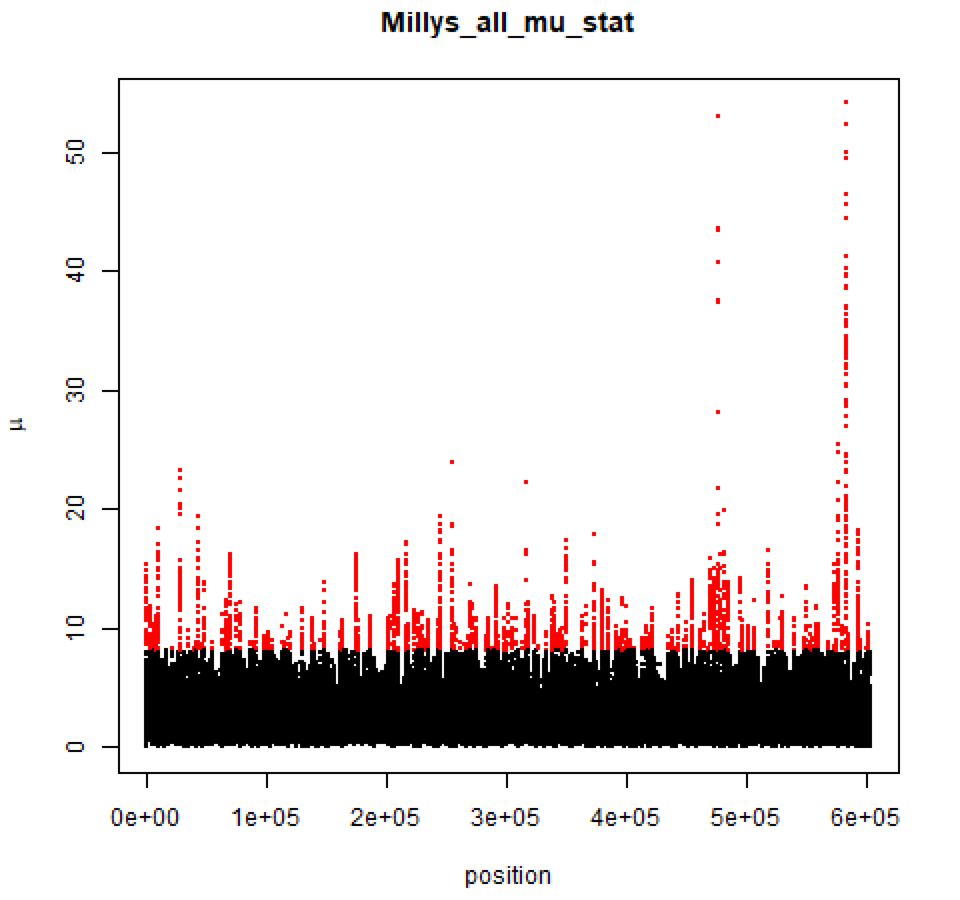

Millys_all_mu_stat
μ
position
0
10
20
30
40
50
0e+00
1e+05
2e+05
3e+05
4e+05
5e+05
6e+05

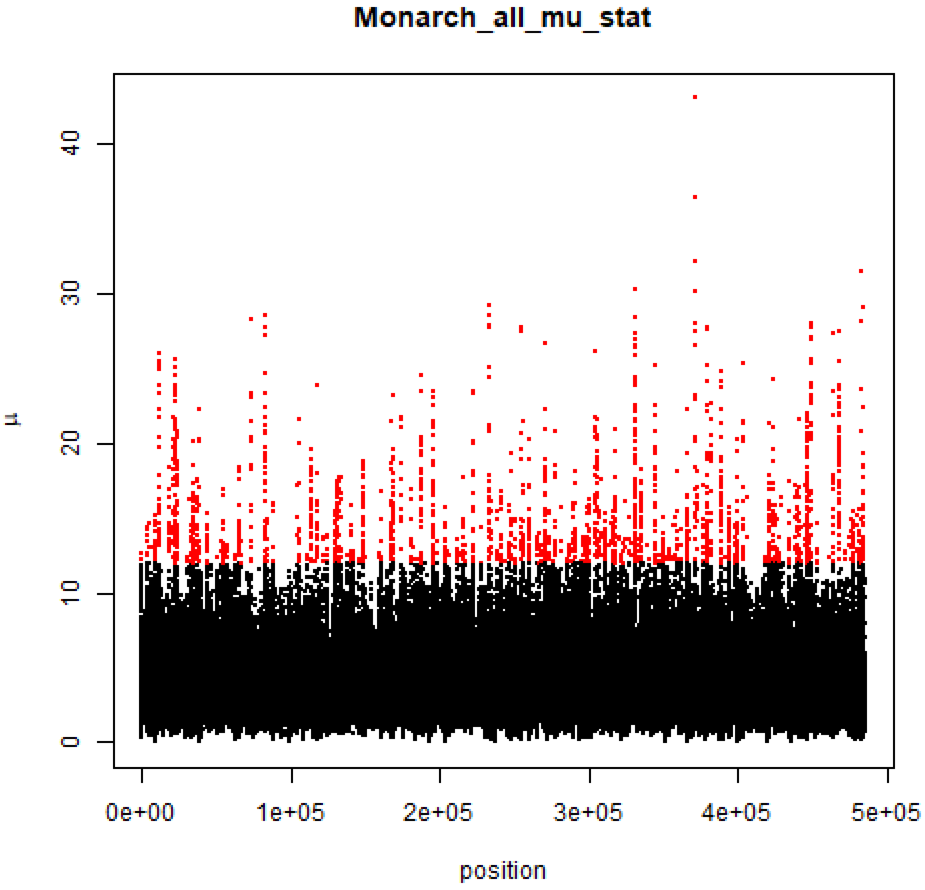

Monarch_all_mu_stat
μ
0
10
20
30
40
0e+00
1e+05
2e+05
3e+05
4e+05
5e+05
position

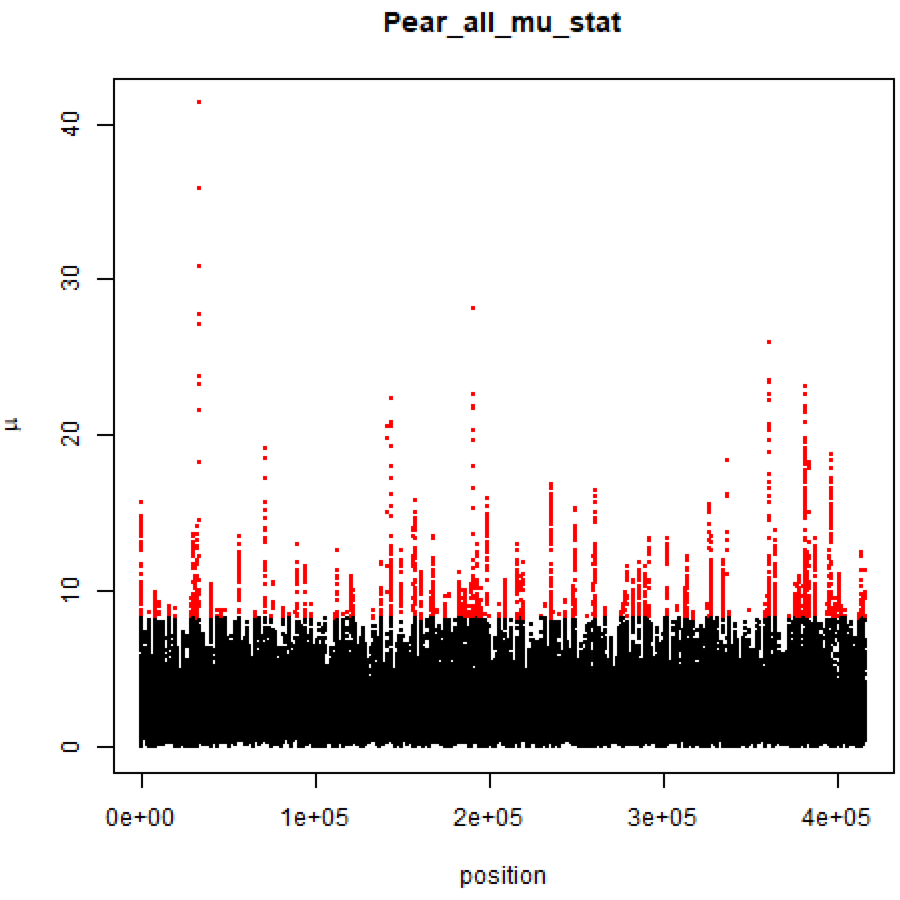

Pear_all_mu_stat
μ
0
10
20
30
40
position
0e+00
1e+05
2e+05
3e+05
4e+05

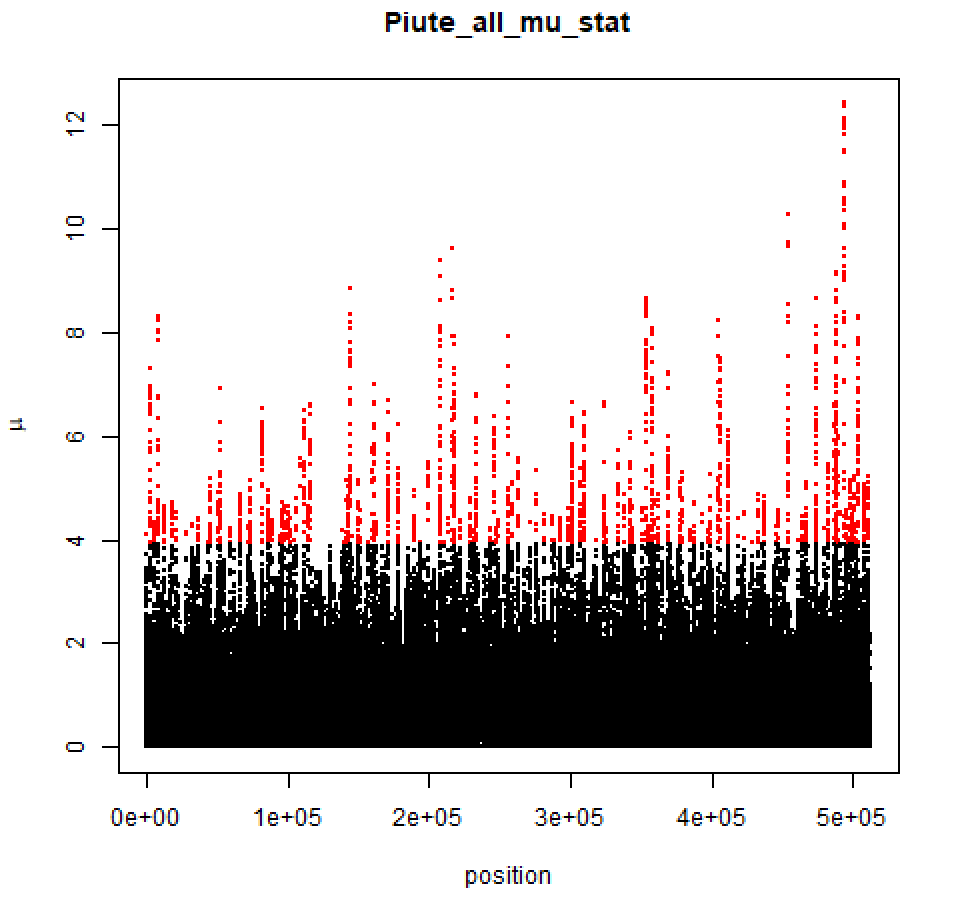
Piute_all_mu_stat
μ
0
2
4
6
8
10
12
0e+00
1e+05
2e+05
3e+05
4e+05
5e+05
position

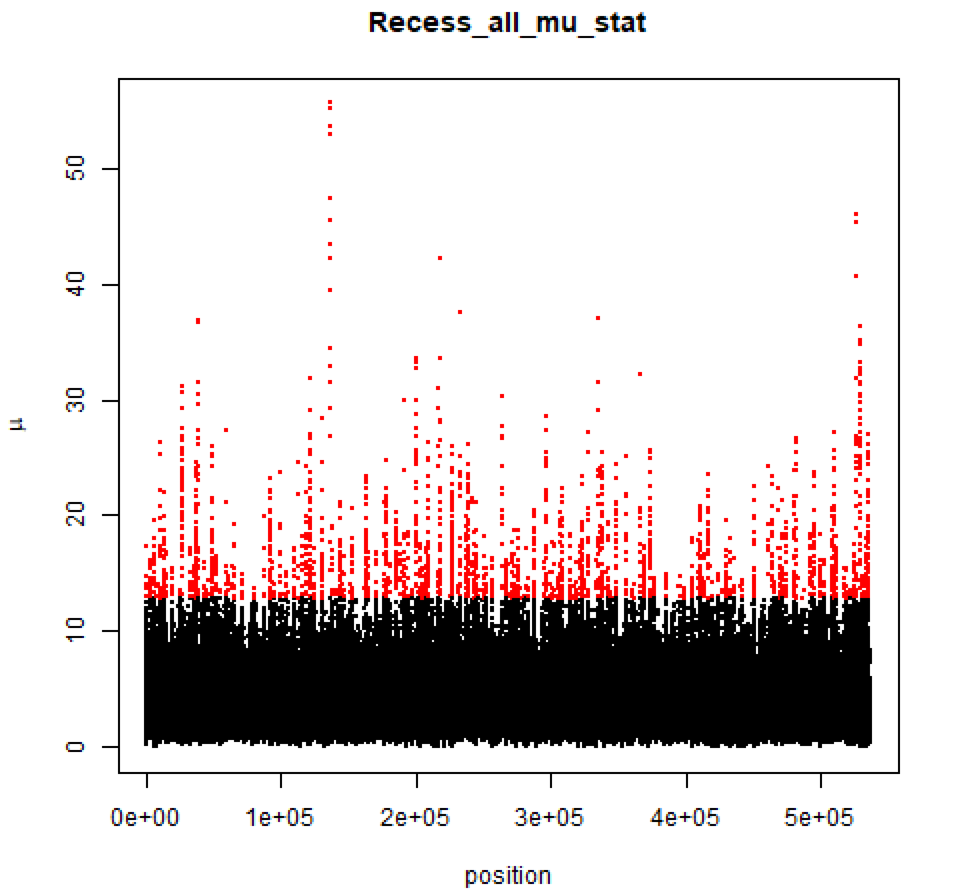
Recess_all_mu_stat
μ
0
10
20
30
40
50
0e+00
1e+05
2e+05
3e+05
4e+05
5e+05
position

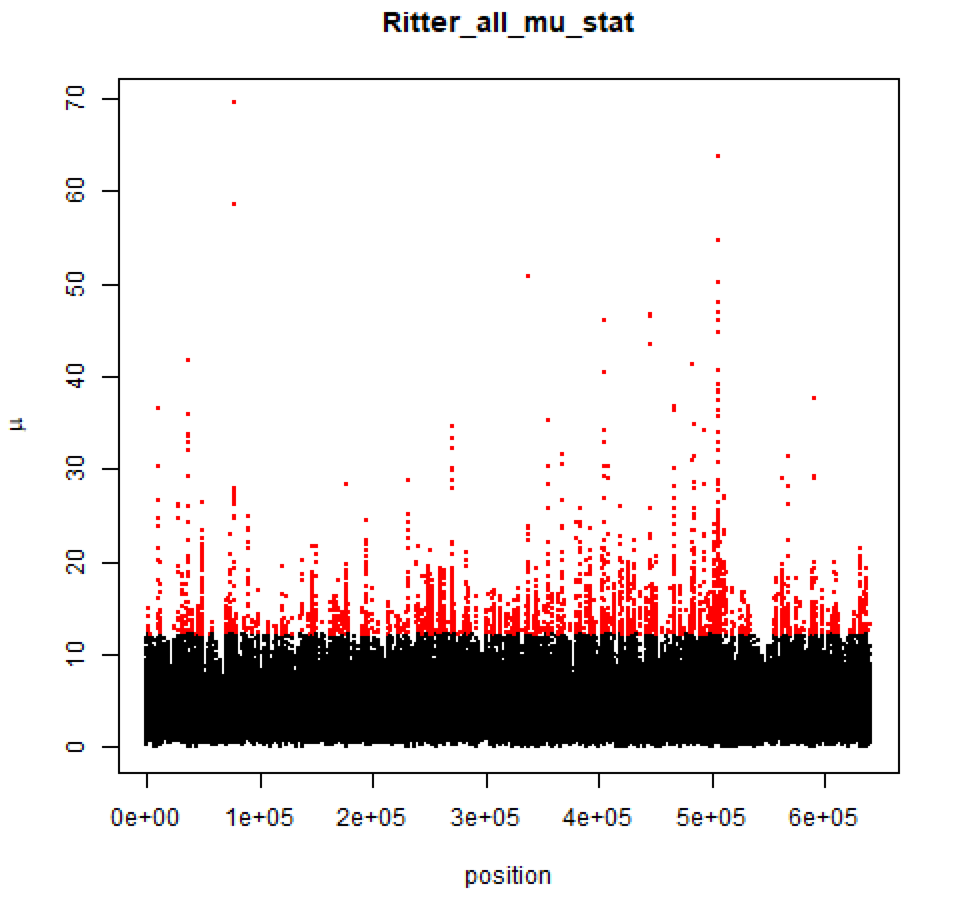
Ritter_all_mu_stat
μ
0
10
20
30
40
50
60
70
0e+00
1e+05
2e+05
3e+05
4e+05
5e+05
6e+05
position

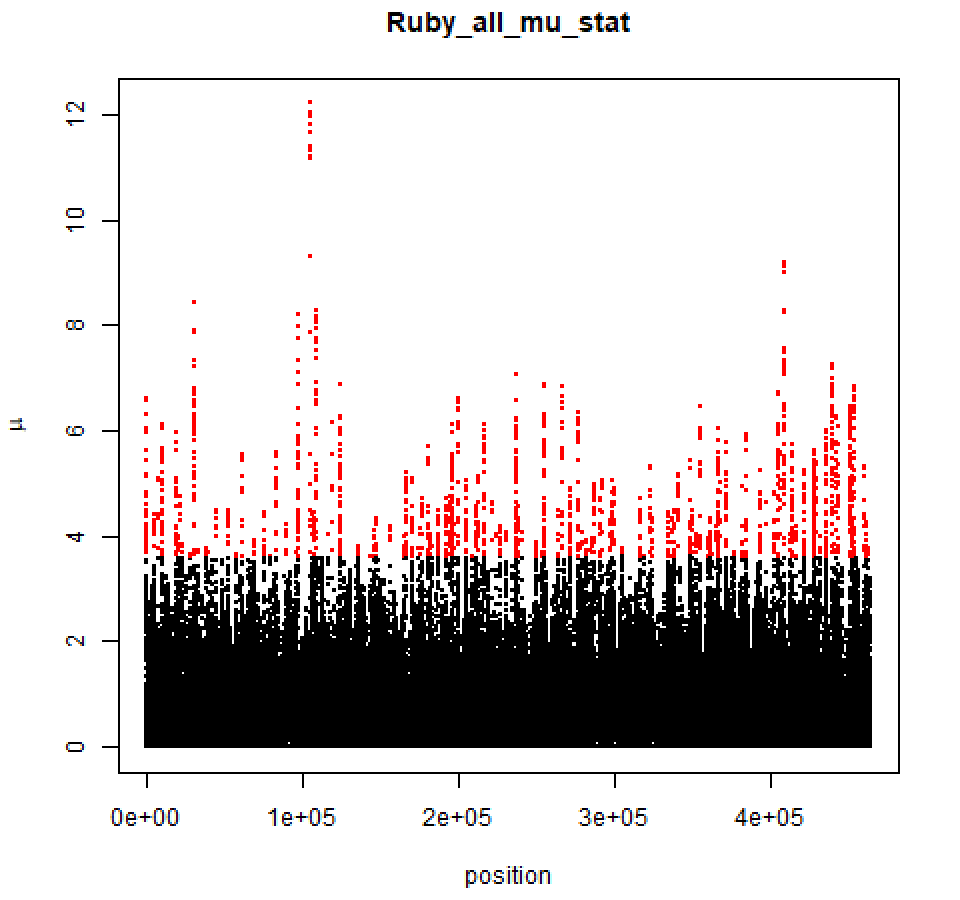

Ruby_all_mu_stat
12
10
8
6
4
2
0
μ
0e+00
1e+05
2e+05
3e+05
4e+05
position

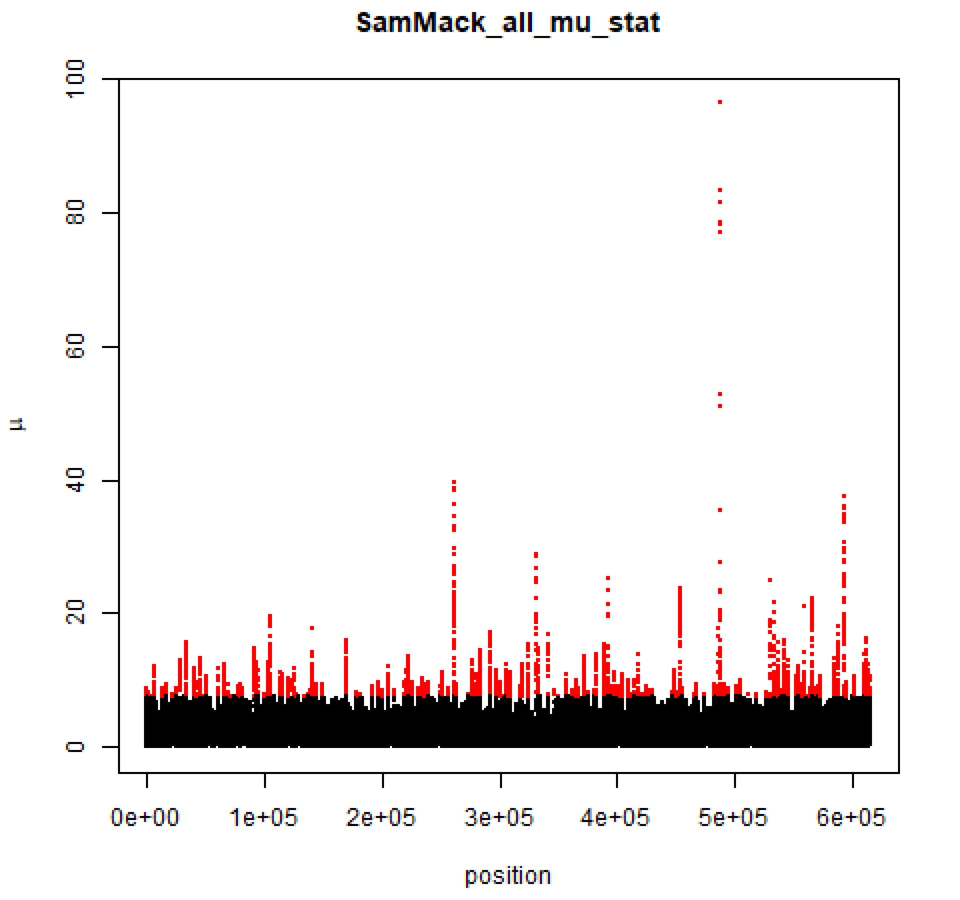
SamMack_all_mu_stat
100
80
60
40
20
0
μ
0e+00
1e+05
2e+05
3e+05
4e+05
5e+05
6e+05
position

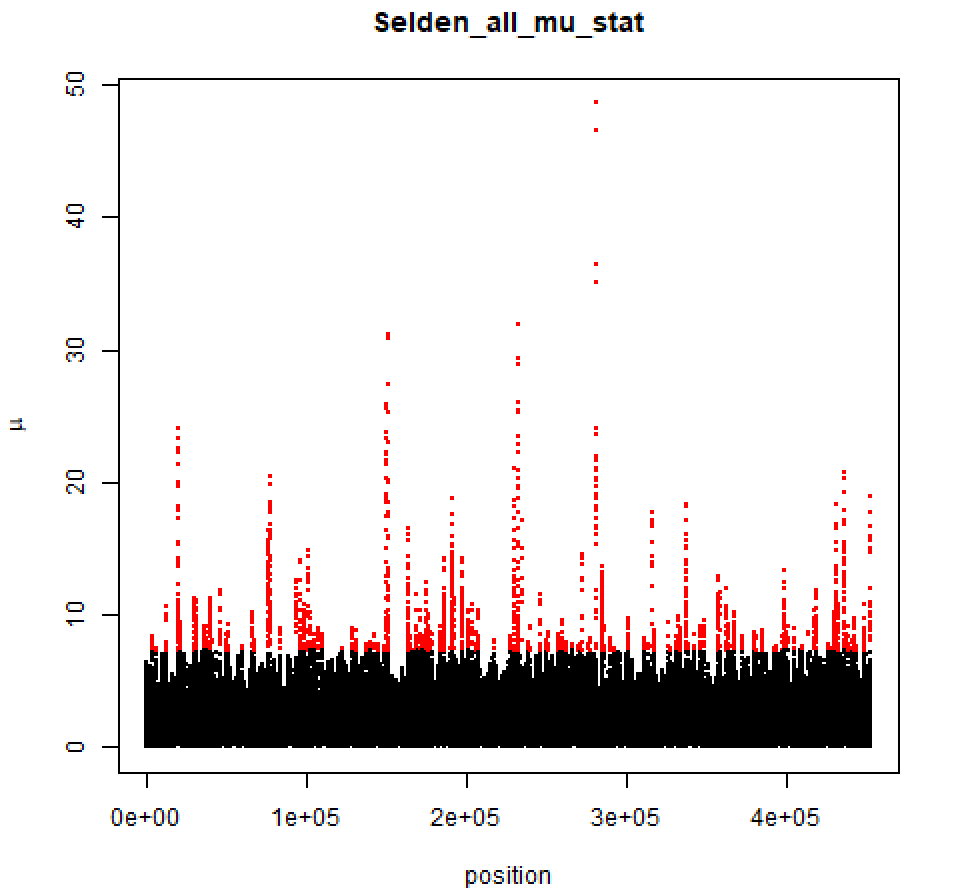
Selden_all_mu_stat
μ
0
10
20
30
40
50
0e+00
1e+05
2e+05
3e+05
4e+05
position

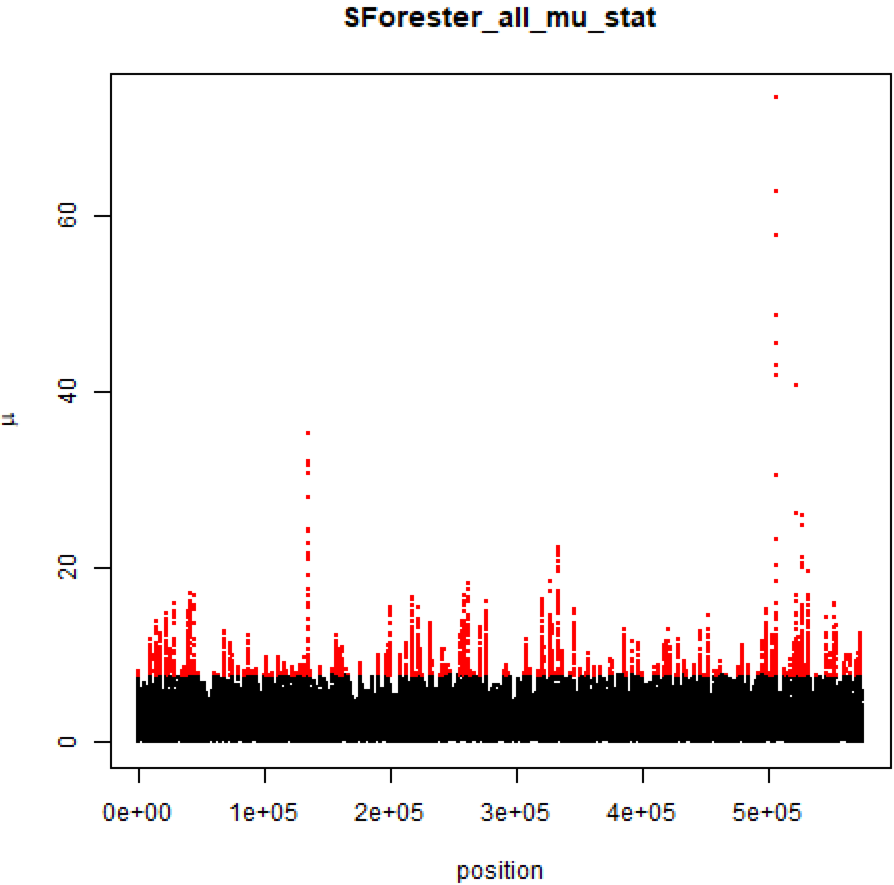

SForester_all_mu_stat
µ
0
20
40
60
0e+00
1e+05
2e+05
3e+05
4e+05
5e+05
position

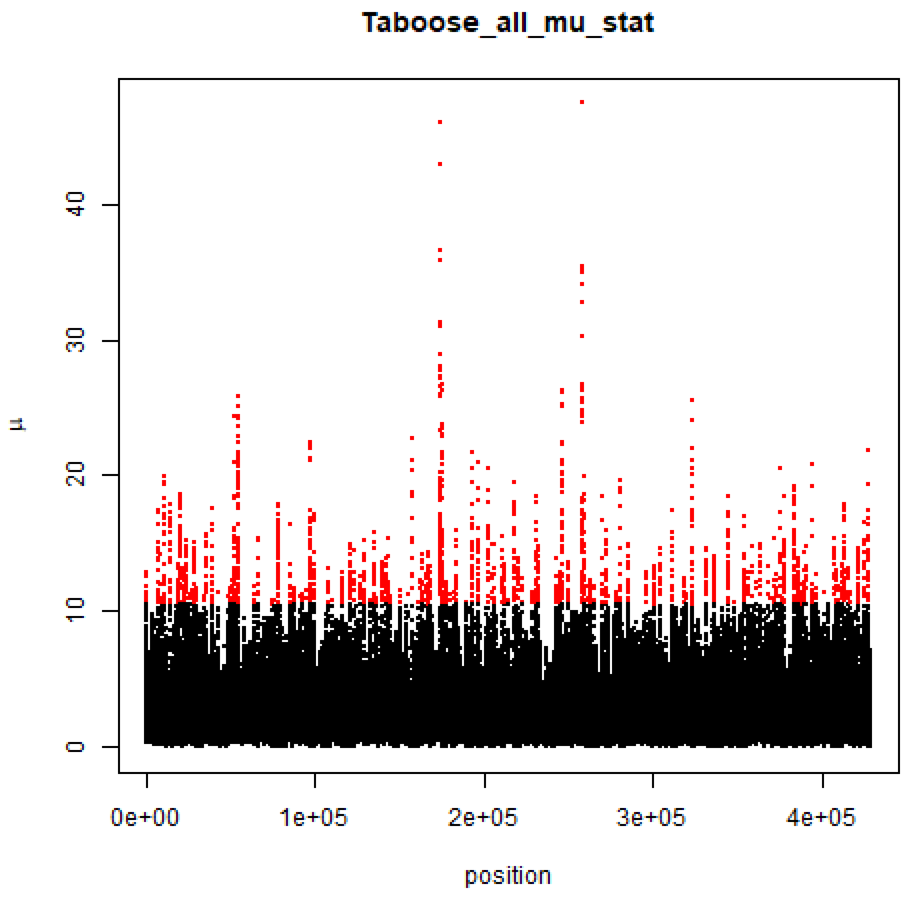
Taboose_all_mu_stat
μ
0
10
20
30
40
0e+00
1e+05
2e+05
3e+05
4e+05
position

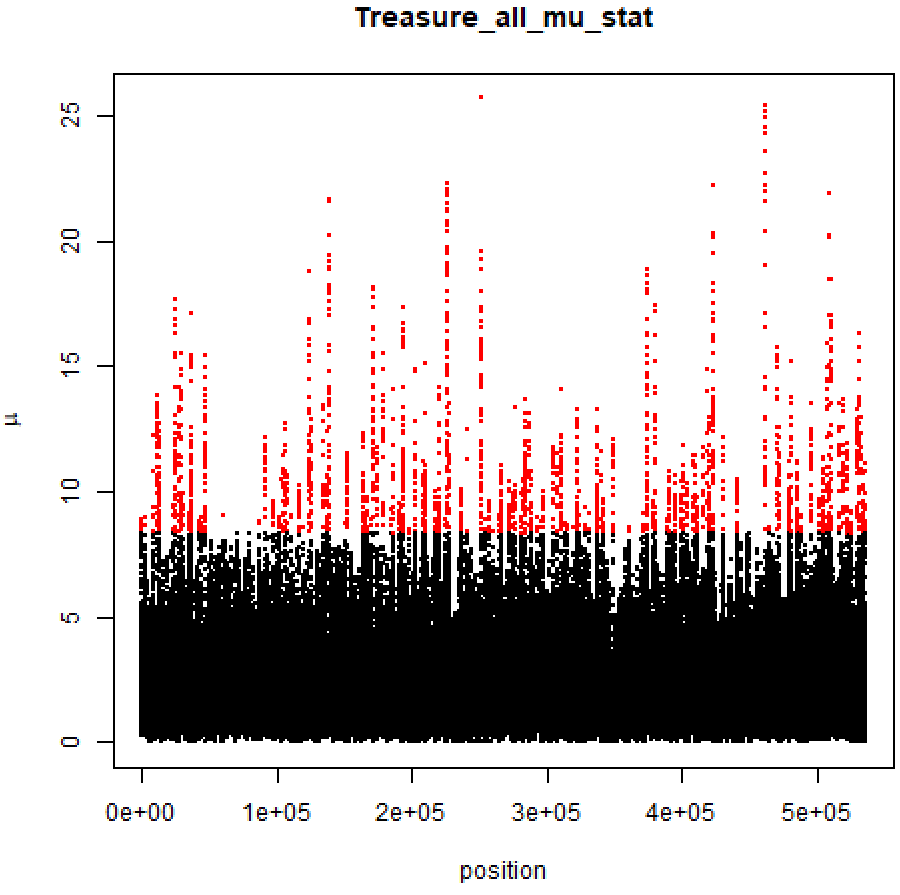

Treasure_all_mu_stat
μ
0
5
10
15
20
25
0e+00
1e+05
2e+05
3e+05
4e+05
5e+05
position

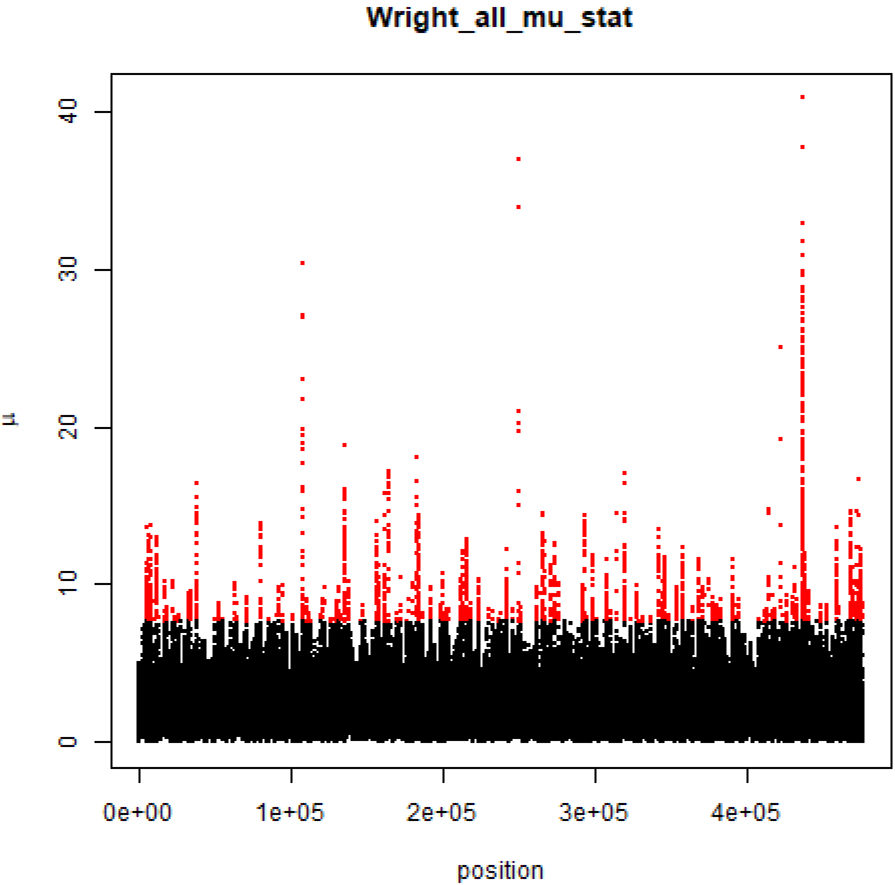

Wright_all_mu_stat
μ
0
10
20
30
40
0e+00
1e+05
2e+05
3e+05
4e+05
position
